# AggreCount: An unbiased image analysis tool for identifying and quantifying cellular aggregates in a spatial manner

**DOI:** 10.1101/2020.07.25.221267

**Authors:** Jacob Aaron Klickstein, Sirisha Mukkavalli, Malavika Raman

**Author notes:** Corresponding author: Malavika Raman.

## Abstract

Protein quality control is maintained by a number of integrated cellular pathways that monitor the folding and functionality of the cellular proteome. Defects in these pathways lead to the accumulation of misfolded or faulty proteins that may become insoluble and aggregate over time. Protein aggregates significantly contribute to the development of a number of human diseases such as Amyotrophic lateral sclerosis, Huntington’s and Alzheimer’s Disease. *In vitro*, imaging-based, cellular studies have defined key components that recognize and clear aggregates; however, no unifying method is available to quantify cellular aggregates. Here we describe an ImageJ macro called AggreCount to identify and measure protein aggregates in cells. AggreCount is designed to be intuitive, easy to use and customizable for different types of aggregates observed in cells. Minimal experience in coding is required to utilize the script. Based on a user defined image, AggreCount will report a number of metrics: (i) total number of cellular aggregates, (ii) percent cells with aggregates, (iii) aggregates per cell, (iv) area of aggregates and (v) localization of aggregates (cytosol, perinuclear or nuclear). A data table of aggregate information on a per cell basis as well as a summary table is provided for further data analysis. We demonstrate the versatility of AggreCount by analyzing a number of different cellular aggregates including aggresomes, stress granules and inclusion bodies caused by Huntingtin polyQ expansion.

## Introduction

The dynamic nature of protein folding necessitates constant surveillance to ensure the production of functional proteins (1). Errors in transcription, translation, or exposure to chemical or oxidative stress exacerbates protein misfolding (2). Protein homeostasis (proteostasis) is monitored during and after protein synthesis to promote refolding by chaperones or clearance of terminally misfolded proteins by either the ubiquitin proteasome system or autophagy (3). Components of protein quality control (PQC) work together to sense and ameliorate cellular stress caused by protein misfolding to maintain proteostasis.

Recent studies have identified an age-associated decline in the quantity, capacity, and efficiency of PQC components (4,5). Thus, maintaining proteostasis is especially important in postmitotic cells such as neurons. This decline is often manifested by the occurrence of age-related neurodegenerative disorders, where misfolded proteins accumulate and overwhelm proteostasis pathways (6). The formation of aggregates is observed in a wide range of neurodegenerative diseases such as Huntington’s disease (HD)(7), amyotrophic lateral sclerosis (ALS)(8), Parkinson’s (9) and Alzheimer’s disease (10), and contributes significantly to disease phenotypes. The morphology, localization, and composition of aggregates in neurodegenerative diseases is complex and heterogenous. Aggregates can be seeded by protein or RNA-based mechanisms and arise in distinct cellular compartments (11). Certain types of aggregates such as aggresomes and inclusion bodies are believed to be protective by sequestering misfolded proteins into a single structure for disposal (12). Furthermore, the morphology of aggregates can be quite distinct and impacts the cellular response. For example, aggregates caused by *C9orf72* gene expansions in ALS form ribbon-like structures that trap proteasomes and exclude ribosomes (13). In contrast, CAG (glutamine, Q) expansions in the Huntingtin gene that form polyQ aggregates in HD are fibrillar and exclude proteasomes (14).

In order to study molecular mechanisms behind protein aggregation, *in vitro* models of protein aggregation have been crucial. Microscopy-based studies investigating cellular aggregates have identified numerous factors that regulate the formation and dissipation of these structures and have informed our understanding of disease (15,16,17). Quantification of protein aggregates has therefore become a convenient and widely accepted measure of disease phenotype severity (18). Our survey of literature relevant to this area suggests that current approaches for quantifying protein aggregates are often focused on manual image analysis (19,20,21). Notably, it is not always clear if the images were analyzed in a blinded fashion (22,23). Manual analysis of aggregates is typically limited to scoring a single cellular feature, such as, area, number, or localization of the aggregate. Such an approach is incapable of capturing multiple features at single cell resolution from hundreds of images. Furthermore, because manual analysis often reports a population phenotype, it is unable to detect subtle differences in sub-populations of cells. Finally, manual image analysis is subjective, error prone and labor-intensive. An increasing number of automatic and semi-automatic image analysis software such as ImageJ and CellProfiler are available to evaluate fluorescence images (24,25). Machine learning algorithms are also increasingly used to aid in the analysis of multi-parametric, complex cellular phenotypes (26). However, incorporating various image analysis modules within these systems to create an analysis pipeline can be time-intensive and unintuitive. These disadvantages highlight a clear need for a user-friendly, flexible, high-throughput cellular image analysis method to quantify aggregates.

In order to improve ease, precision, speed and reproducibility of aggregate quantification, we developed AggreCount, a novel image processing macro, using ImageJ as a platform, to process, analyze and quantify images. This automated tool uses common stains for nuclei, cell bodies and aggregates as input and quantifies aggregate number, area and cellular localization on a cell-by-cell basis. In addition to delivering an unbiased analysis of images, this level of detail provides increased analytical depth and better evaluation of subtle phenotypes that may otherwise be overlooked. All processing steps use native ImageJ plugins and functions such that additional downloads or plugins are not necessary. Furthermore, this tool is written in the ImageJ macro language, which is more accessible to those with limited programming knowledge. We have used AggreCount to analyze and quantify a variety of cellular aggregates including aggresomes, stress granules, aggresome-like induced structures (ALIS) and Htt polyQ inclusions. AggreCount will permit researchers to carry out unbiased quantification of diverse protein aggregates and is particularly suited to high-content image-based screening approaches.

## Results

### Defining settings in AggreCount for the multi-parametric quantification of aggregates

The general workflow for analysis with the AggreCount macro involves three phases: assembling images, macro setup, and batch analysis. Images that will be analyzed as part of one experiment must be gathered into a single folder (the macro code and detailed instructions to run AggreCount are provided in Supplementary information 2 and 3). AggreCount is compatible with both common (e.g TIFF, JPEG etc) and proprietary file formats commonly used in microscopy that are recognized by ImageJ. Each image must consist of one channel for nuclei and one for aggregate quantification. An optional third channel may be used for cell body identification. After images are compiled, the macro may be run to determine analysis settings (Figure 1 and Supplementary information 1). The setup will guide the user through selecting appropriate channels for each fluorophore, adjusting the threshold, determining appropriate size cut offs for aggregates, nuclei and cells as well as the best processing strictness for cell bodies. The main settings window (Supplementary information 1) allows further refinement of the macro options. Once the variables are set, the user may proceed to batch analysis for all images in the folder. The collected data is saved as tab-delimited text files that can be imported into excel or another data analysis software.

**Figure 1.**
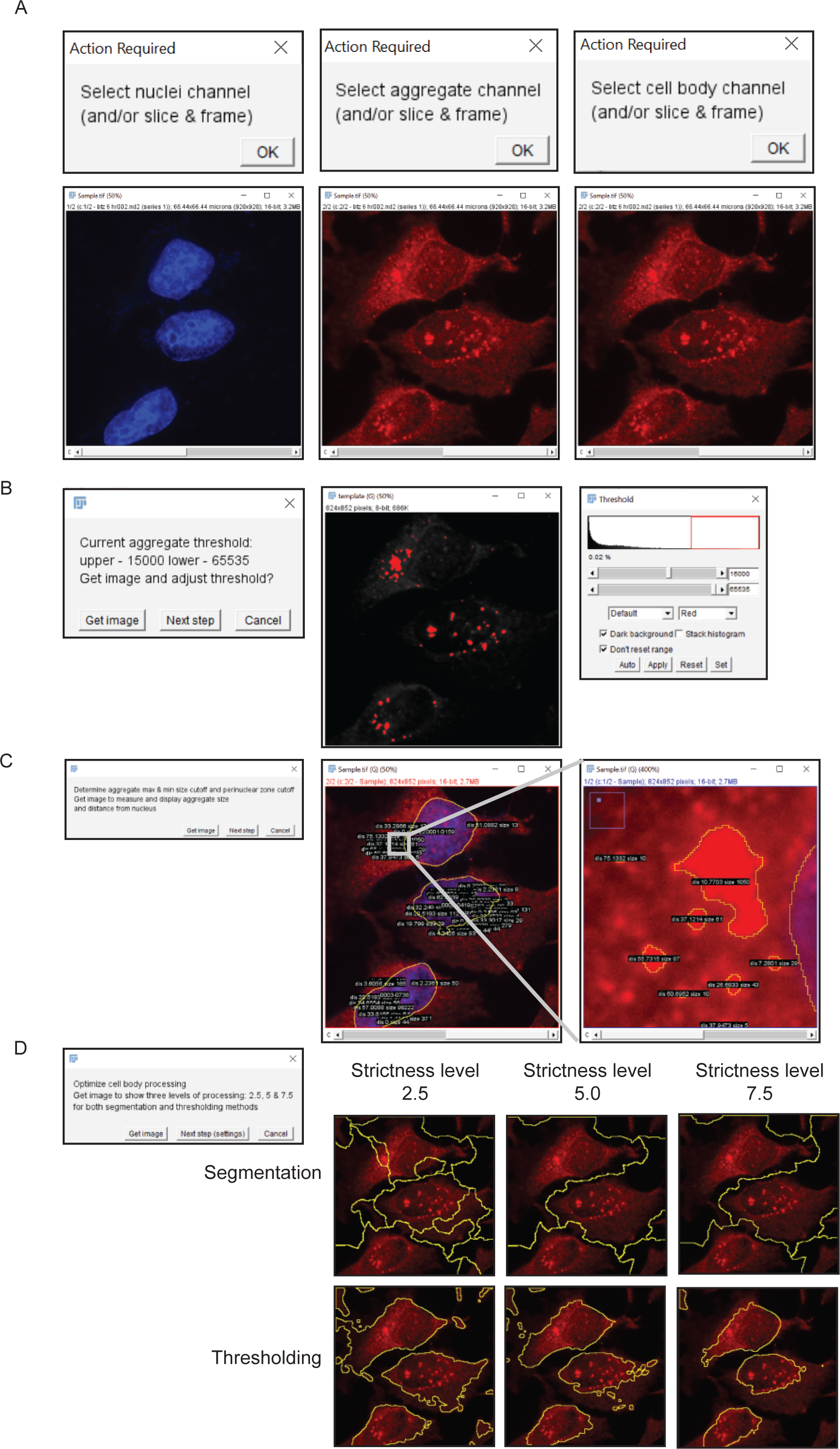
Overview of AggreCount workflow. **A.** User selects fluorescent channels corresponding to nuclei, aggregates, and cell bodies. **B.** Threshold for aggregates is adjusted by the user through the native ImageJ threshold tool. **C.** AggreCount displays a sample image analysis with aggregate size and distance from nucleus. **D.** Multiple levels of processing strictness are displayed for segmentation and thresholding of cell bodies.

After selecting the folder of image files to be analyzed and selecting the ‘setup’ option, the first image in the analysis folder will be opened. The user will be prompted to select the channel for the nuclei, aggregates and cell bodies (Figure 1A). For cell body segmentation, we recommend a dedicated channel using commercially available dyes such as CellMask; however, an antibody that detects a cytosolic protein (here we have used ubiquitin) works equally well. The user may also select the stack and frame from z-stacks or movies. AggreCount will use the selected slice and frame for that channel for every image processed. For z-stacks or movies, it may be preferable to collapse the image (e.g. average Z projection) before analysis for more consistent results, although in such instances the nuclear localization of aggregates should be carefully assessed. The user will then be prompted (‘Get image’) to select an appropriate threshold in the aggregate channel using the ImageJ threshold tool (Figure 1B). AggreCount will continue to loop through the prompt to allow the user to open multiple images using the ‘Get image’ option and continue refining the threshold. It is suggested that for each analysis, at least one image with aggregates and one image without aggregates is used as positive and negative controls respectively to set the threshold. The final threshold parameter defined by the user will be applied to all images. Once the threshold is established, the user is prompted (‘Next step’) to open an image that will be processed for nuclei and aggregates using the previously established threshold. The size and distance of each aggregate from the nucleus will be displayed on the composite image (Figure 1C). This allows the user to determine appropriate size cut-offs for aggregates and distance limits to the perinuclear region.

The main AggreCount settings window allows for manual changes to each option (Supplementary Figure 1). The values defined by the user during the setup phase will be auto-populated in the settings window. The remaining values are default values and may be changed. Perinuclear distance is the pixel distance on either side of the nuclear ROI that defines the perinuclear zone. This setting may be useful for quantification of structures such as aggresomes that reside adjacent to the nucleus. User-defined variables are available to set the aggresome, aggregate, nucleus and cell body size minimums in microns. Additionally, there is an option to set a maximum size for aggregates. The selected channel, slice and frame for each structure to be analyzed will be displayed and may be changed manually. If there is no channel for cell bodies or if the analysis does not require the identification of cell bodies, uncheck “Find cell bodies”. The macro will calculate distances but not assign aggregates to cells.

Unchecking the save results option allows the macro to be run without any files being saved and may be useful when optimizing settings. Otherwise, the macro will create a new folder to save the result files. These results consist of summary text files: (1) summary analysis for all images (AC_analysis\dataset_summary.txt), (2) all cells (AC_analysis\dataset_cells.txt), and (3) all aggregates (AC_analysis\dataset_aggregates.txt). Additionally, a detailed results file (‘FileName’_analysis.txt) and a .zip file containing cell, nuclei and aggregate ROIs identified for each image will be saved. In batch mode AggreCount will proceed to analyze each image in the analysis folder. Images will open in the background and will not be visible to the user. The status of the analysis may be monitored in the summary table.

### Identification and classification of aggregates

Nuclei can be visualized using standard dyes (DAPI, Hoechst or DRAQ5 for example), and processing nuclei images is minimal (Figure 2A). The macro uses the “Enhance contrast” function before subtracting background as a function of the mean pixel intensity. The proportion of the mean that is subtracted can be modulated by changing the “strictness” setting in the settings window. Then a median filter is applied to smooth the image while maintaining borders. Properly defined borders are critical for this analysis as the nucleus ROIs define sub-compartments within the cell. The image is then thresholded using the “Make Binary” function followed by the “Dilate” and “Fill Holes” functions. “Dilate” is used to better define the border of the nucleus as the intensity of the fluorescent stain is often reduced at the edges of nuclei causing the outermost portion to be excluded during the “Make Binary” function.

**Figure 2.**
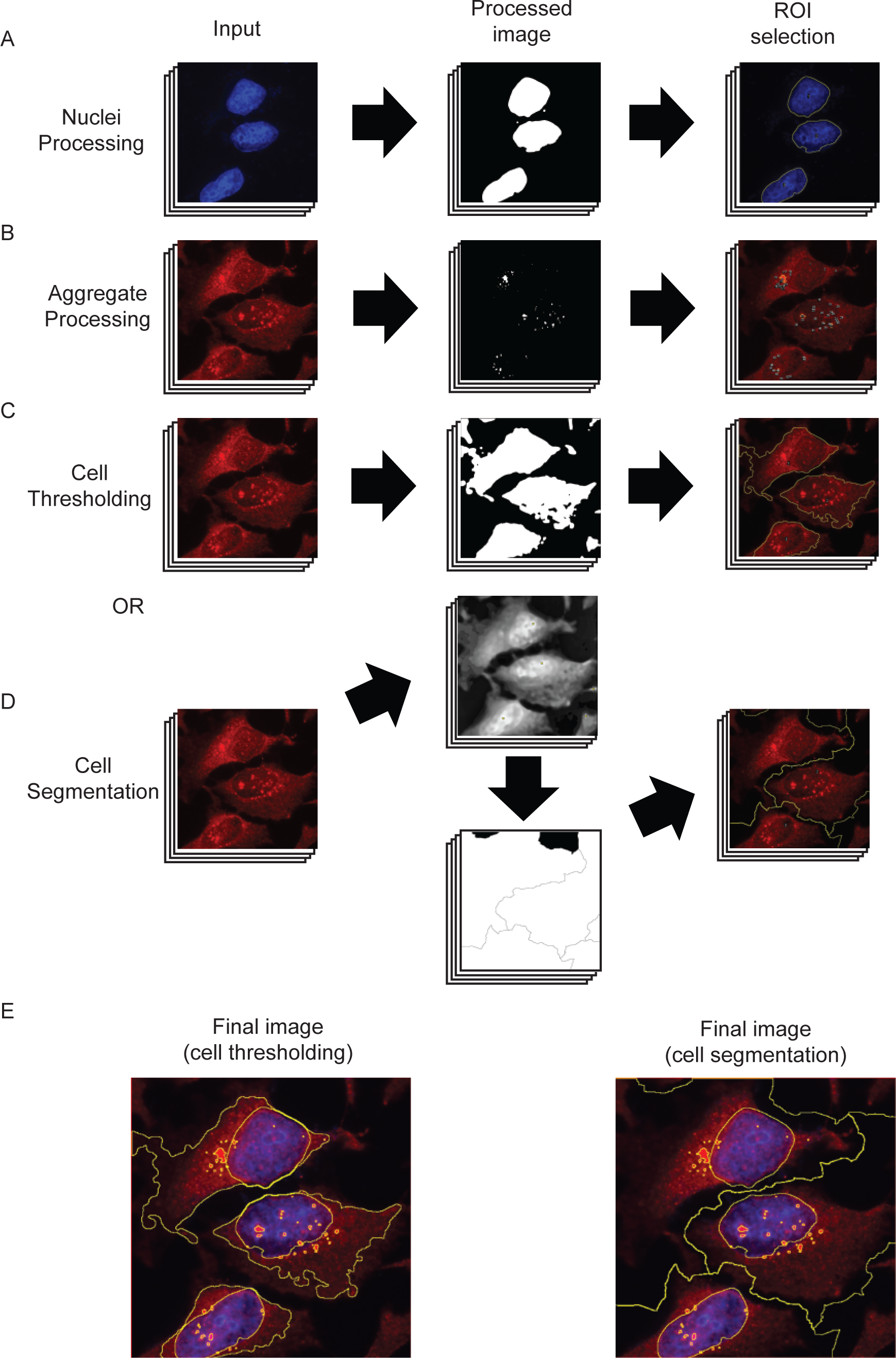
AggreCount image processing steps. **A.** Nuclei are processed via fluorescent signal enhancement, conversion to a binary image, and capture of ROIs. **B.** Aggregates are processed via the default method or user-inserted processing steps before being converted to a binary image for ROI identification. **C.** Cell thresholding via enhancement of fluorescent signal, conversion to binary image, and capture of ROIs similar to that of nuclei processing. **D.** Cell segmentation process that identifies fluorescent maxima (top middle panel), segments cell via a watershed algorithm into a Voronoi diagram (bottom middle panel) which is used to capture ROIs. **E.** Final merged image with nucleus, aggregate and cell body ROIs.

The default method for processing aggregate images aims to enhance relatively small, bright puncta (Figure 2B). The first step uses a crude background subtraction based on the mean pixel intensity of all areas with fluorescent signal to remove diffuse fluorescent noise. This is followed by isolating bright puncta using the difference of Gaussians approach (27). The image is then converted to 16-bit and the threshold that was set previously is applied. This binary image is used with the “Analyze particles” function to capture ROIs which are filtered based on the previously set size criteria.

To analyze aggregates at single cell resolution, AggreCount uses two methods for cell body processing: segmentation and thresholding (Figure 2C-E). The main difference between these methods is that segmentation defines a cell by the portion of the image that contains it while thresholding aims to capture the cell as a ROI without any surrounding area. Segmentation (Figure 2D) is often more accurate in differentiating cells that are in close proximity while thresholding (Figure 2C) provides information on the cell itself such as size and pixel intensity but may omit portions of cells or erroneously merge adjacent cells. Cell area may be an important metric if treatments collapse the cytoplasm which may spatially restrict aggregates to the perinuclear zone. AggreCount provides both methods so the user may choose the appropriate one for their images. Images of differing cell density may require optimization of segmentation parameters so testing multiple images is good practice.

Segmentation is achieved using the “Find Maxima” function that finds points of maximal fluorescent intensity in local areas in a merged image of the cell body and nucleus. This function utilizes a prominence setting that determines the trough required between maxima which can be changed from the settings window using the cell body strictness option. After the maxima are determined, a pixel intensity-based watershed algorithm segments the image into a Voronoi diagram which is used for ROI capture (Figure 2D and E). The cell thresholding method uses a process similar to that of the nuclei processing that enhances contrast, subtracts background as a proportion of the mean (that may be changed with the strictness option), and applies a median filter (Figure 2C). If cell bodies are not stained, the user may deselect the “Find cell bodies” option in the main settings window. The macro will proceed with localization analysis but will not be able to provide analysis on a cell-by-cell basis.

### Distance calculation and aggregate localization

Once nuclei and aggregate ROIs have been captured, AggreCount will proceed to calculate the distance between aggregates and nuclei (Figure 3A i and ii). These distances are used to spatially classify aggregates within the cell and are calculated based on all the pixel coordinates encompassing the perimeters of the aggregate and nucleus (Figure 3A iii). The distance of each coordinate of the aggregate is calculated for each coordinate of the nucleus perimeter using the distance formula (Figure 3A iv). If the “Find cell bodies” option is unchecked, aggregate distance will be calculated from the nearest nuclei. This value is used to localize the aggregate within one of three subcellular compartments: cytosol, perinuclear zone, or nucleus (Figure 3B). The user defines the distance by which the perinuclear zone is determined using the perinuclear distance option in the settings window. This value is calculated on either side of the nucleus. Any aggregate that is at least partially contained in this zone is categorized as perinuclear. Thus, aggregates classified as perinuclear can reside on either side of the nuclear envelope. We have specified the default value for perinuclear distance cutoff as 10 pixels for 60x magnification images, higher values may result in nuclear aggregates being classified as perinuclear. The cutoff value should be empirically adjusted for images of different magnification. Aggregates that are farther inside the nucleus are considered nuclear and aggregates that are excluded from both the perinuclear zone and the nucleus are considered cytosolic.

**Figure 3.**
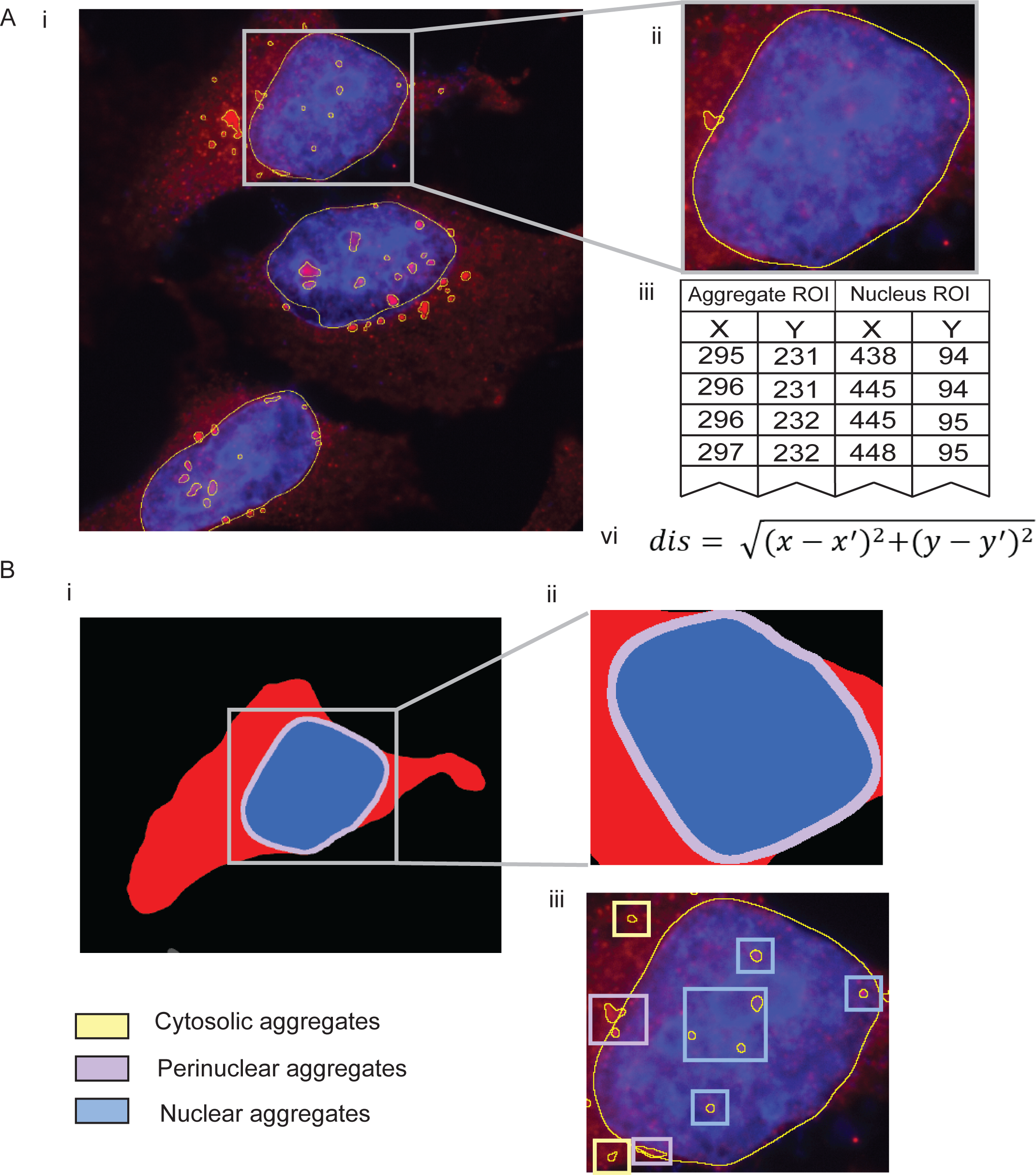
Delineating subcellular compartments. **A.** After ROIs have been captured (i and ii), each coordinate for aggregate ROIs (iii) is compared to each coordinate of nucleus ROIs to determine distance using the distance formula (iv). The smallest distance value is recorded as the aggregate distance. **B.** Subcellular compartments within the cell are determined by the nucleus ROI and the user-defined distance parameter (i and ii) and labelled as cytosolic, perinuclear and nuclear (iii).

### Deploying AggreCount to quantify cellular aggregates

We present several examples of different cellular aggregates to demonstrate the versatility of AggreCount for their quantification. First, we quantified ubiquitin-positive aggresomes and aggregates that arise in cells upon proteasome inhibition. Aggresomes are peri-nuclear, membrane delimited structures that form via retrograde trafficking of smaller cytosolic aggregates via the dynein motor (28). HeLa Flp-in TRex cells were treated with the reversible proteasome inhibitor bortezomib for 8 or 18 hours before being fixed and stained for ubiquitin and nuclei (Hoescht) (Figure 4A). Images were analyzed using AggreCount. As shown in Figure 4B, both timepoints have an increased number and size of aggregates compared with the untreated controls. The AggreCount analysis allows for the stratification of aggregates by subcellular location at single cell resolution. This deeper analysis reveals that the 18-hour treatment reduces the number of cytosolic aggregates while the number of perinuclear and nuclear aggregates remain stable (Figure 4C). However, the size of the perinuclear aggregates greatly increases (Figure 4D and E). For this analysis, AggreCount analyzed 221 cells in less than five minutes. In contrast, manual analysis of all these parameters for this number of cells would have taken on average several hours.

**Figure 4.**
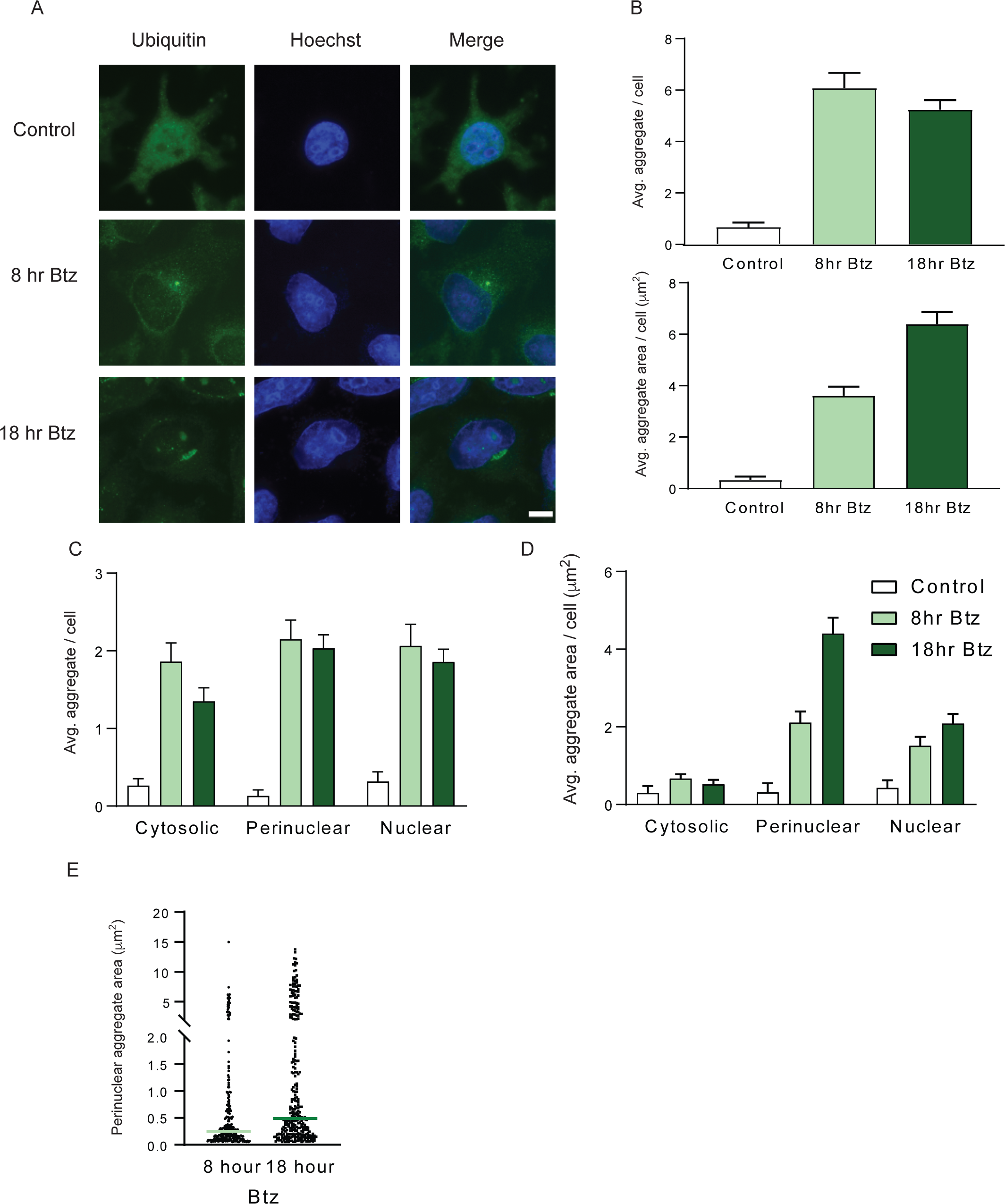
Analysis of Bortezomib induced aggregates by AggreCount. **A.** HeLa Flp-in TRex cells were treated with 1 μM Bortezomib (Btz) for 8 or 18 hours, fixed and stained with an anti-ubiquitin antibody to visualize ubiquitin-positive aggregates and aggresomes. Nuclei were stained with Hoechst. **B.** AggreCount was used to determine the average number of aggregates per cell (upper panel) and average aggregate area (lower panel). **C.** The number of aggregates in different cellular compartments was quantified. **D.** The area of aggregates in different cellular compartments was determined. **E.** Prolonged Btz treatment (18 hr) leads to an increase in perinuclear aggresomes. Scale bar is 10 μm.

Puromycin treatment causes premature chain termination during translation and the accumulation of defective ribosomal products (DRiPs) (29,30). DRiPs are frequently sequestered into punctate, ubiquitin-positive structures in the cytosol referred to as aggregate-like induced structures (ALIS) (31,22). AggreCount was able to effectively identify ALIS in puromycin treated cells (Figure 5A). We have previously shown that ubiquitin X domain containing 1 (UBXN1) is required for the clearance of ALIS (33). The increase in ALIS in UBXN1 knockout (KO) cells was effectively captured by AggreCount (Figure 5A-C). Notably, ALIS in UBXN1 KO cells were increased in the peri-nuclear area relative to wildtype cells (Figure 5C), illustrating capability of AggreCount to spatially localize aggregates when perturbations alter the subcellular localization of aggregates.

**Figure 5.**
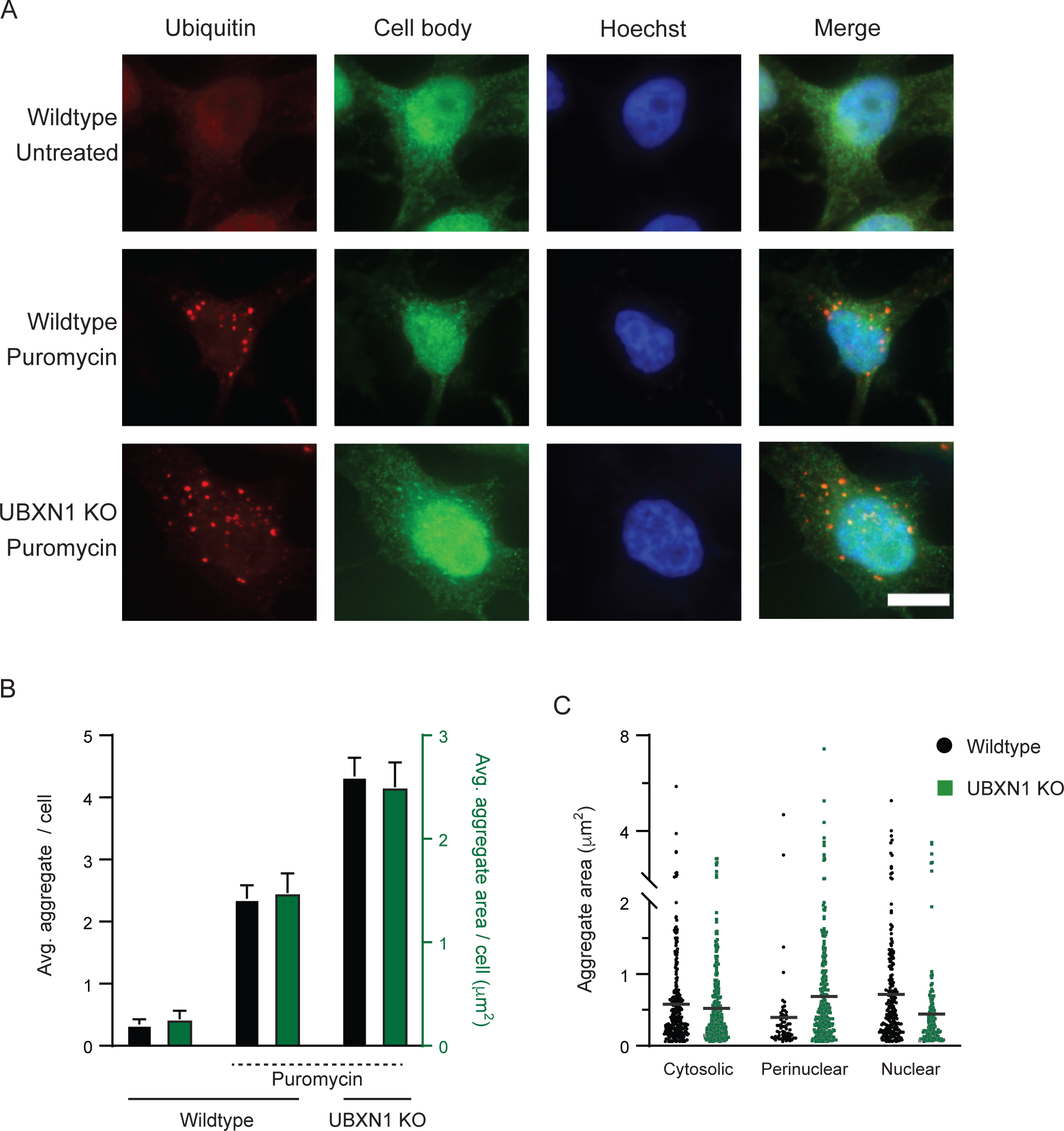
Aggregate like Induced Structures (ALIS) quantification by AggreCount. **A.** HeLa Flp-in TRex cells (wildtype and UBXN1 knockout, KO) were treated with 5 μg/ml of puromycin for 2 hours, fixed and stained with an anti-ubiquitin antibody to visualize ubiquitin-positive ALIS. Nuclei were stained with Hoechst. **B.** AggreCount was used to determine the average number of ALIS per cell (black bars) and average ALIS area (green bar). UBXN1 KO cells have a greater number of ALIS with increased areas **C.** The area of ALIS in different cellular compartments was determined. Scale bar is 10 μm.

We next addressed RNA-based cellular aggregates such as stress granules. Stress granules form as a result of a variety of cellular stressors that cause stalling of translation and disassembly of ribosomes (34). Stress granules contain 40S subunits of ribosomes, ribonucleoproteins and RNAs and their inappropriate persistence in neurons is linked to neurodegeneration (35). We induced stress granule formation in HeLa cells stably expressing GFP-tagged G3BP1 (a bona fide stress granule component), using sodium arsenite, a well-established chemical that robustly induces stress granule formation (Figure 6A). Release of cells post-sodium arsenite treatment into drug-free media leads to the dissolution of stress granules (Figure 6A and B). Previous studies have demonstrated a role for p97, an essential, ubiquitin-selective, AAA-ATPase that is critical for multiple PQC processes in the cell, in the clearance of stress granules (36,37,38). Treatment of cells with an ATP-competitive p97 small molecule inhibitor (CB-5083) during the formation or release periods produced distinct outcomes (39). Cotreatment of cells with both sodium arsenite and CB-5083 results in the formation of fewer and smaller stress granules compared to sodium arsenite treatment alone (Figure 6A and B). Conversely, p97 inhibition during the release period prevents the clearance of stress granules predominantly in the perinuclear region (Figure 6A-D). These results suggest that p97 may also have an unappreciated role in stress granule formation that warrants further investigation.

**Figure 6.**
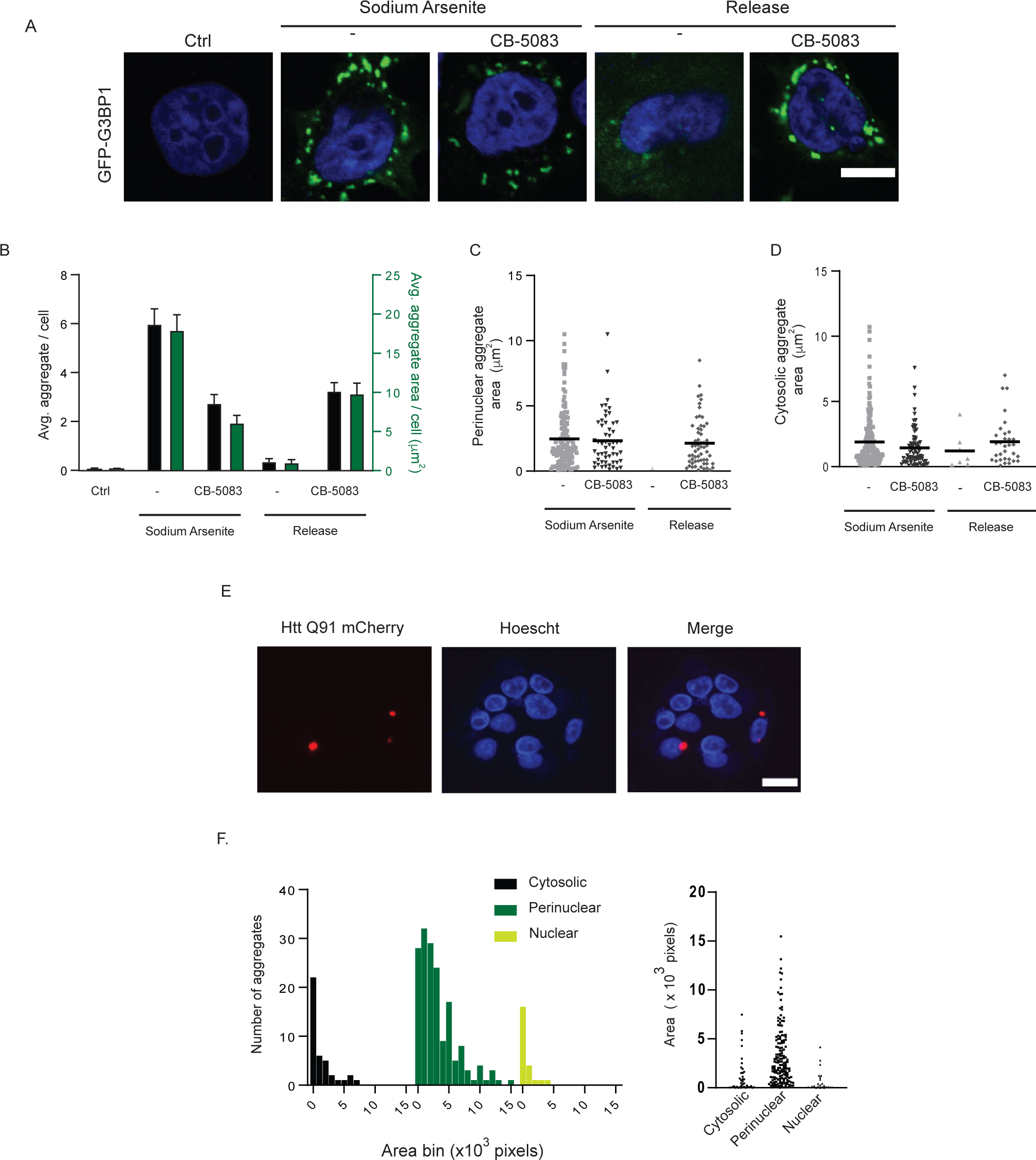
Analysis of stress granules and inclusion bodies by AggreCount. **A.** HeLa Flp-in TRex cells stably expressing the stress granule marker GFP-G3BP1 were treated with 0.5 mM sodium arsenite for 4 hours. Cells were pretreated with 5 μM CB-5083 (p97 inhibitor) for 1 hour prior to addition of 0.5 mM sodium arsenite for 4 hours. Cells were released into drug free media or in media containing CB-5083 fixed and imaged. Nuclei were stained with Hoechst. **B.** AggreCount was used to determine the average number of stress granules per cell (black bars) and average stress granule area (green bars). **C and D.** The area of perinuclear (C), cytosolic stress granules (D) was quantified. **E.** U2OS Htt91-mCherry cells were induced with 1 μg/ml of doxycycline for 48 hours. Cells were fixed and imaged for inclusion bodes. Nuclei were stained with Hoechst. **F.** The area of individual inclusion bodies based on their cellular location was quantified and binned based on the indicated size bins. Scale bar is 10 μm.

Expansion of a CAG tract in the first exon of the *Huntingtin* (*Htt*) gene beyond a threshold of approximately 35-40 repeats causes Huntington’s disease and results in a mutant Htt protein containing an expanded polyglutamine (polyQ) segment (40). Expression of exon 1 Htt fragments longer than 40 residues leads to the formation of insoluble amyloid-like aggregates of Htt known as inclusion bodies and causes neurodegeneration (40). We measured inclusion body formation using a doxycycline inducible Htt polyQ91 construct tagged with mCherry (41). AggreCount was able to identify polyQ aggregates in cells and classify their localization as predominantly perinuclear as previously reported (Figure 6E and F) (42). Previous studies have suggested that large inclusion bodies formed by polyQ are largely cyto-protective and smaller soluble aggregates are cytotoxic (43). Because AggreCount can effectively identify and quantify the area of cytosolic and perinuclear aggregates (Figure 6F); conditions that shift the dynamic between cytosolic aggregates and inclusion bodies can be rapidly assessed using AggreCount.

## Discussion

ImageJ and CellProfiler are two of the most commonly used software for immunofluorescent image analysis. Both provide a robust set of tools for image processing, thresholding, and capture of data (44). Our survey of literature pertaining to image analysis of aggregates found no uniform method of image analysis. In many cases, aggregates were quantified manually which introduces bias and human error. CellProfiler allows for the batch processing and analysis of images which helps alleviate these problems However, the software is optimized for high-throughput image analysis and can be difficult to adjust for medium or low throughput experiments. Additionally, a certain level of image processing background is required for initial development of CellProfiler pipelines (24). Here, we present the AggreCount macro that is based on the widely used FIJI distribution of ImageJ (45). This macro provides an unbiased automated platform for aggregate quantification on a cell-by-cell basis. By utilizing the built-in ImageJ macro language, this tool is easily modifiable for different fluorophores. We have deposited AggreCount to GitHub to be downloaded as a macro for the reproducible and efficient analysis of aggregates by the research community. While we believe our method for processing images with aggregates works well for a wide variety of applications, we understand that these processing steps may not be applicable for all experiments. Thus, the portions of the macro which contain the processing steps for aggregates, nuclei and cells are highlighted and may be altered by the user (Supplementary Information 1B). The ImageJ macro language was designed such that users with limited programming knowledge may utilize it. The “Record” function in ImageJ may be used to further customize the script in a user-specific manner.

Here we demonstrate that AggreCount can capture and quantify a variety of distinct cellular aggregates that arise due to specific perturbations, including aggresomes, ALIS, stress granules and Htt inclusion bodies. While the AggreCount macro was designed for the quantification of aggregates, it may be used to quantify and localize organelles such as mitochondria or lysosomes, and disease markers in neurodegeneration such as TDP-43 or FUS that relocalize from the nucleus to the cytoplasm and aggregate. Additionally, as this tool analyzes ROIs on a cell-by-cell basis, it may be used to parse out different cell types in a heterogenous population provided individual cell populations can be uniquely labelled. The AggreCount macro provides a powerful platform for unbiased, automated quantification and localization of cellular aggregates.

## Experimental Procedures

### Antibodies and Chemicals

The mouse anti-ubiquitin (FK2) used for immunofluorescence was from EMD Millipore. Hoechst dye was from Sigma and Alexa Fluor-conjugated secondary antibodies were from Molecular Probes. Bortezomib was obtained from Selleckchem, Sodium arsenite and puromycin were from Sigma.

### Cell Culture

HeLa Flp-in TRex (kind gift from Brian Raught University of Toronto), HeLa Flp-in TRex GFP-G3BP1 and U2OS HttQ91-mCherry (kind gift from Ron Kopito, Stanford University) were cultured in Dulbecco’s modified Eagle’s medium (DMEM) supplemented with 10% fetal bovine serum (FBS) and 100 U/ml penicillin. Cells were maintained in a humidified, 5% CO2 atmosphere at 37°C. Cells were treated with 1 μM bortezomib, 0.5 mM sodium arsenite, 5 μM CB-5083 and 5 μg/ml of puromycin for the indicated times. HttQ91-mCherry expression was induced by treating cells with 1 μg/ml of doxycycline for 48 hours. UBXN1 knockout cells were previously described (33). GFP-G3BP1 stable HeLa Flp-in TREX cell lines were generated by lentiviral transduction as previously reported (46). Cells expressing low levels of GFP-G3BP1 were isolated by flow cytometry. Care should be taken to determine whether treatment of cells with specific agents causes the cytosol to collapse or shrink. In such cases, cytosolic aggregates may be overwhelmingly identified as perinuclear. In such cases, it is useful to use CellMask and the threshold tool to measure the area of cells between treatment conditions.

### Immunofluorescence and microscopy

HeLa Flp-in T-REX and U2OS cells were grown on coverslips (no. 1.5) in a 12-well plate. For optimal image analysis, we recommend imaging low to moderate cell densities as images with cells in close proximity to one another negatively impact accurate cell counts and segmentation. Cells were washed briefly in phosphate-buffered saline (PBS) and fixed with either 4% paraformaldehyde at room temperature (puromycin treatment) for 15 minutes or ice cold methanol at 4°C for 10 minutes. Cells were washed three times in PBS, and then blocked in 1% bovine serum albumin (BSA) 0.3% Triton X-100 in PBS for 1 hour. For the puromycin treatment, cells were blocked in 3% chicken serum plus 0.1% Triton X-100 in PBS for 1 hour. The coverslips were incubated with the indicated antibodies overnight at 4°C in a humidified chamber, washed, and incubated for an hour with the appropriate Alexa Fluor-conjugated secondary antibodies for 1 hour in the dark at room temperature. Cells were washed with PBS, and nuclei were stained with Hoechst dye and mounted onto slides. Images were collected by using a Nikon A1R scan head with a spectral detector and resonant scanners on a Ti-E motorized inverted microscope equipped with a 60x Plan Apo 1.4-numerical-aperture (NA) objective lens. The indicated fluorophores were excited with either a 405-nm, 488-nm, or 594-nm laser line.

## Data availability

The AggreCount macro is available in Supporting Information associated with this manuscript and will be uploaded to GitHub.

## Acknowledgements

We are grateful to Dr. Ron Kopito for the U2OS polyQ91-mCherry cell line. We thank members of the Raman lab, Dr. Peter Juo and Dr. Shireen Sarraf for suggestions and feedback.

**Supplementary Information 1.** Settings and aggregate processing may be customized by the user. **A.** The main dialogue window may be used to change analysis parameters in a user-friendly manner. **B.** Processing steps have been highlighted within the macro code for easy editing.

**Supplementary Information 2.** Code for AggreCount

**Supplementary Information 3.** Step by step instruction for running AggreCount. Sample images to can be found at the following link: https://tufts.box.com/s/ys3kktb5ujdnilyqu8mnup3mrtiej6b6

## Supplementary Information 1

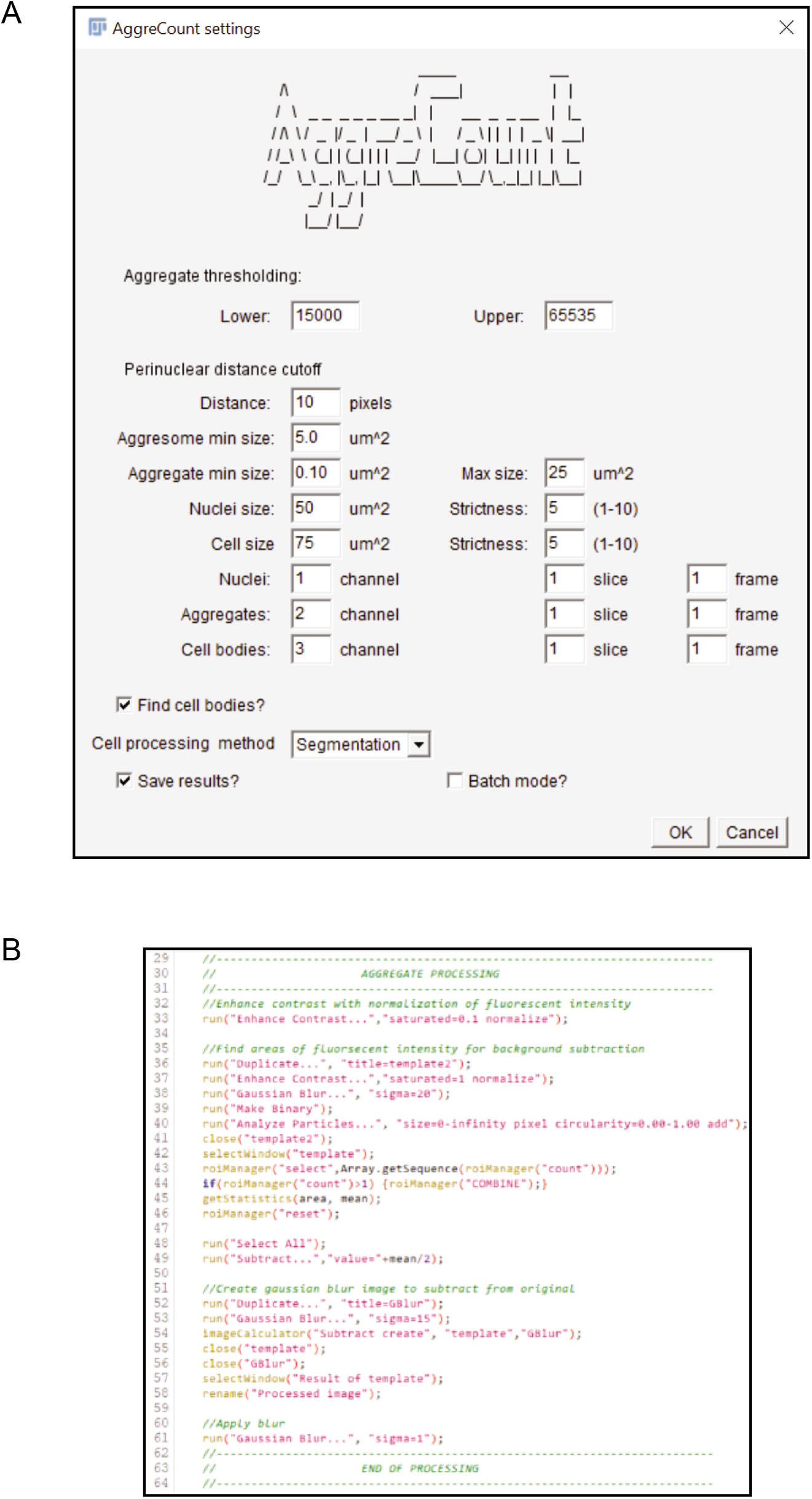

## Supplementary Information 2

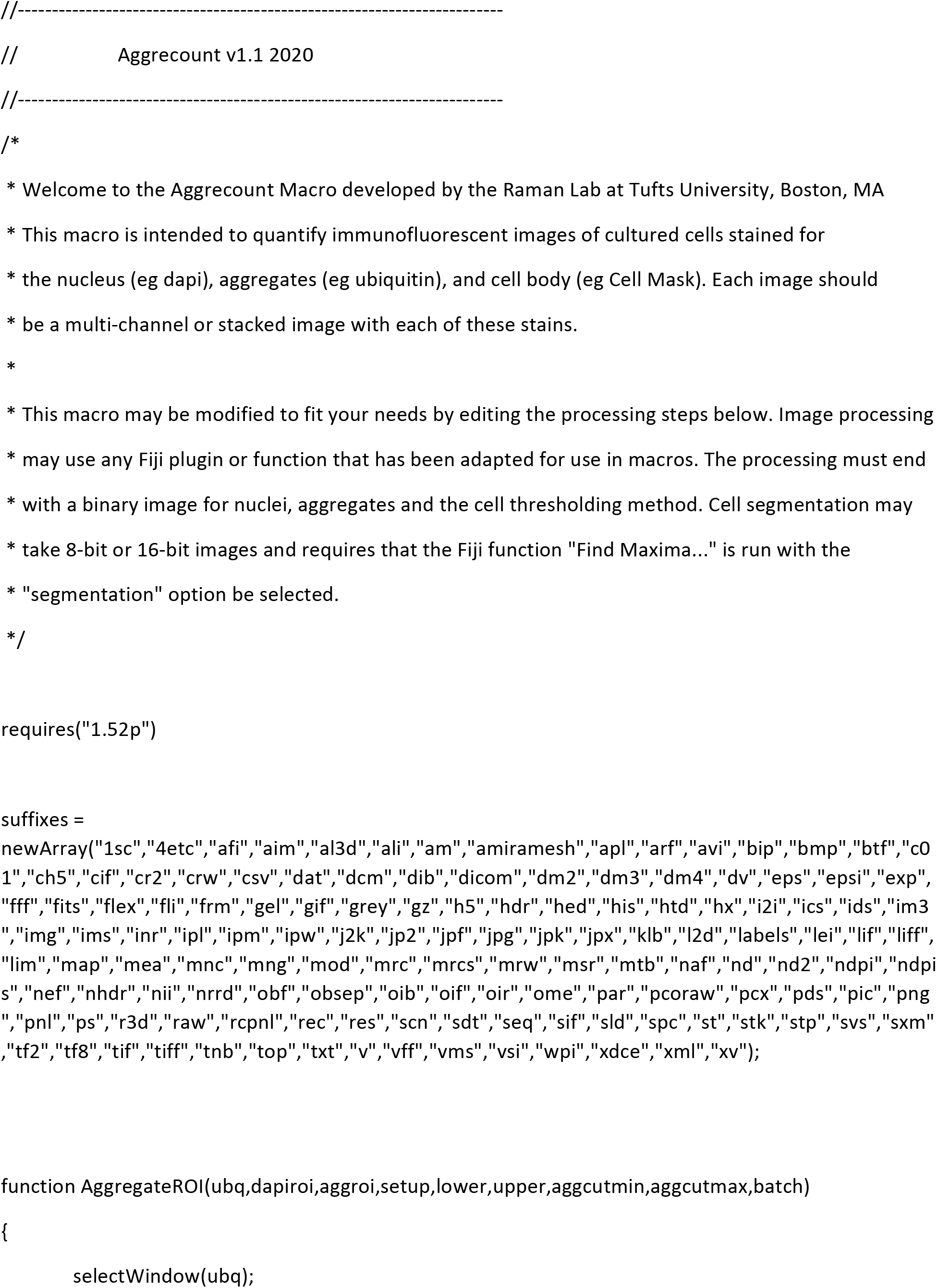

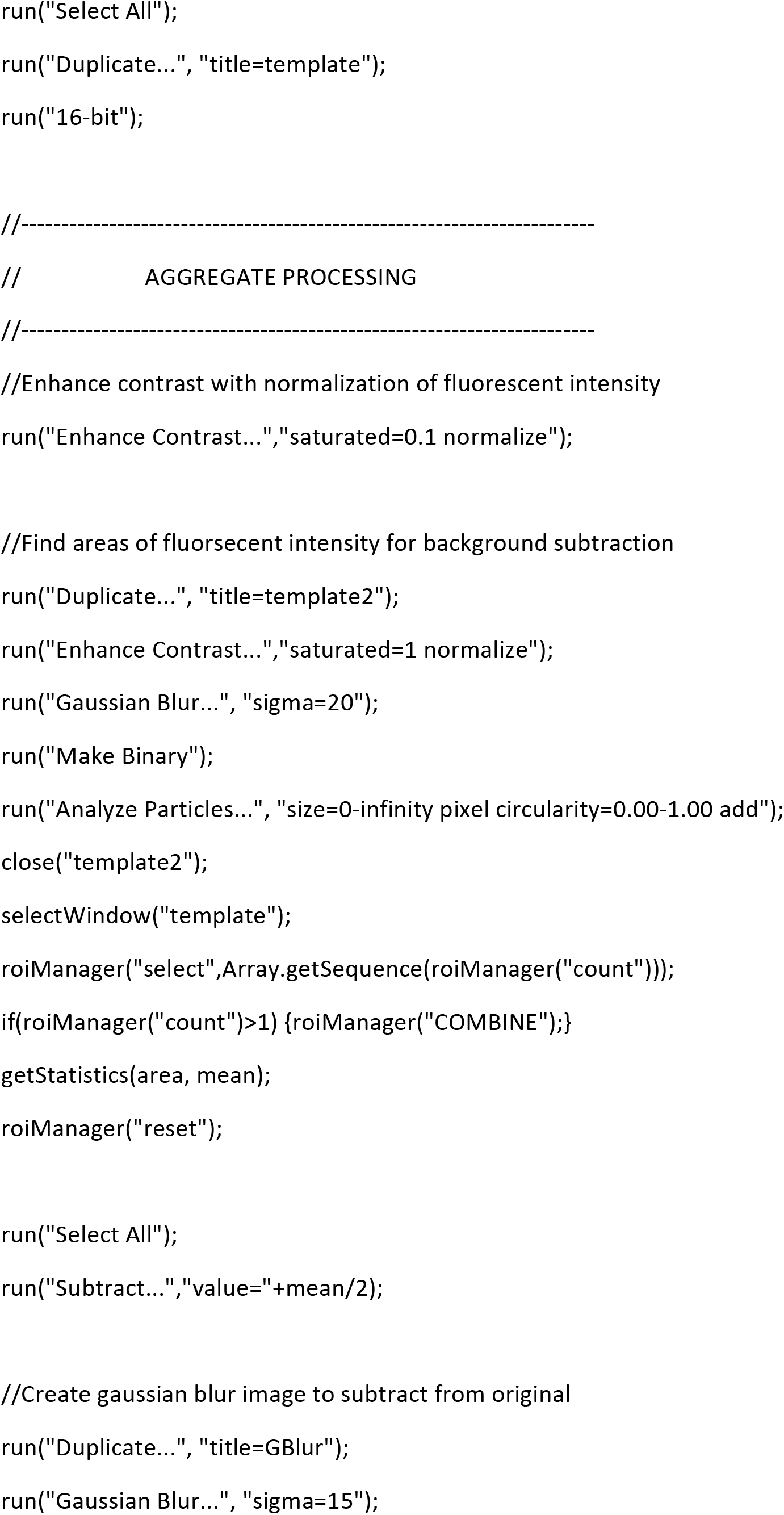

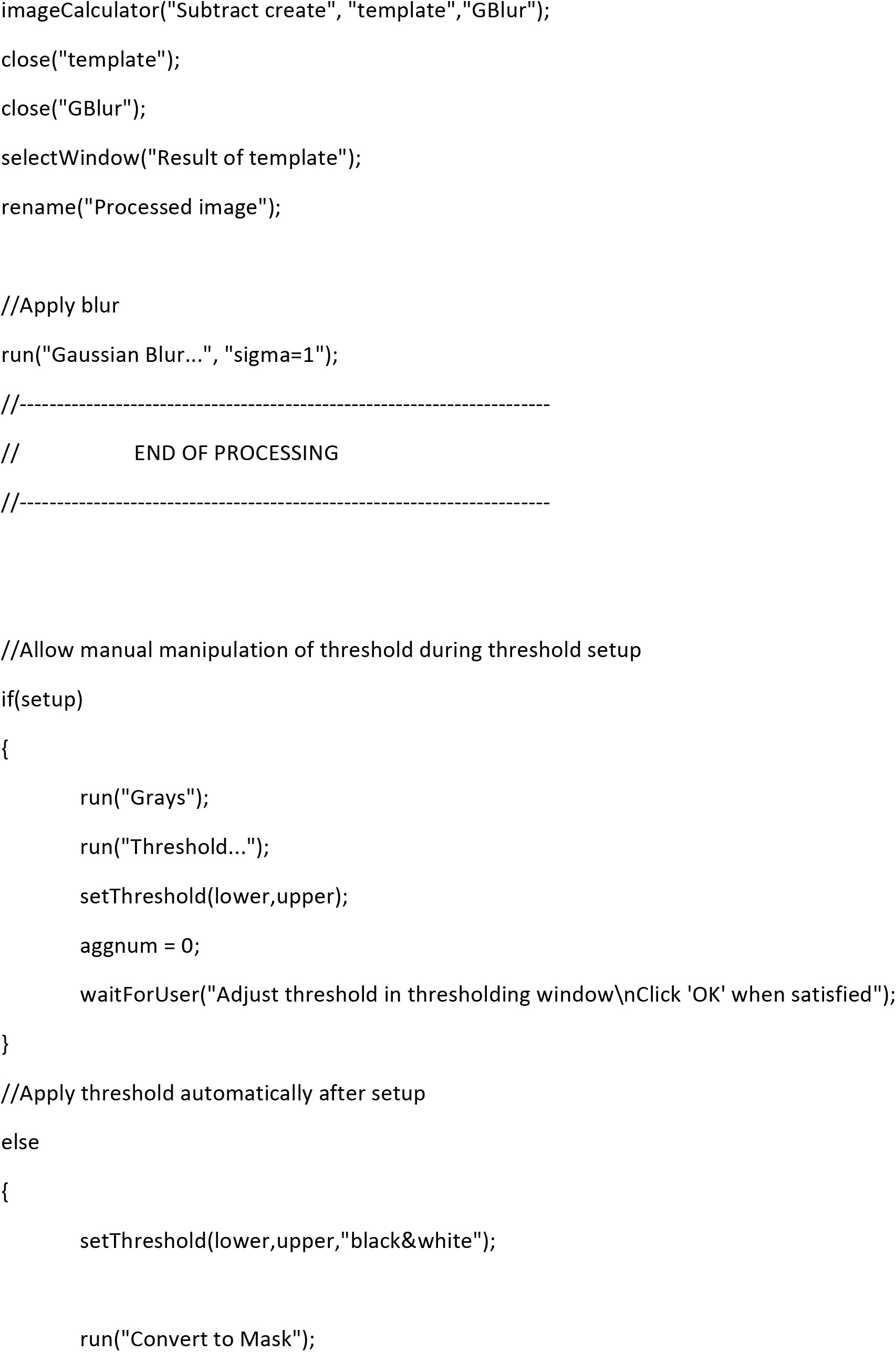

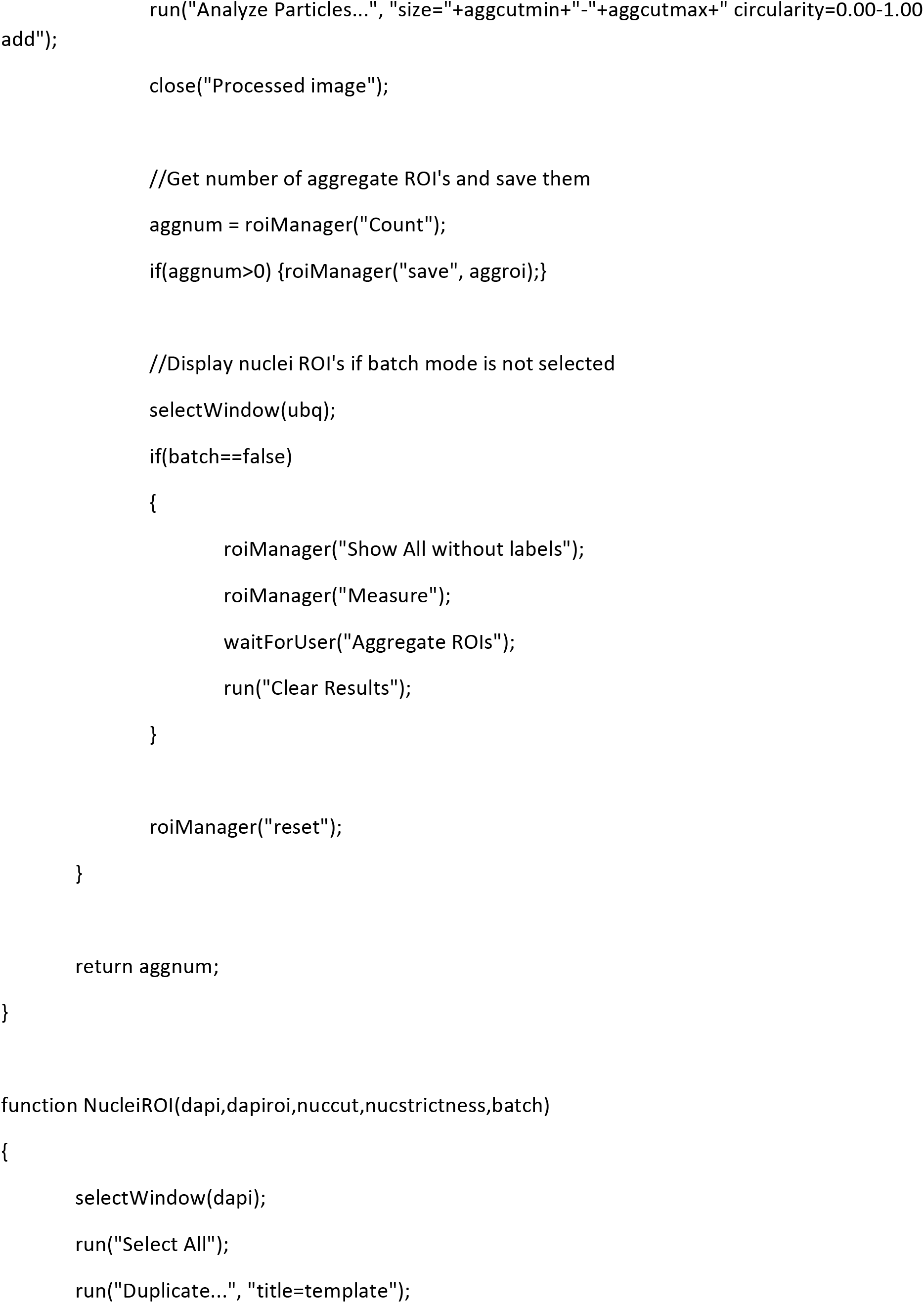

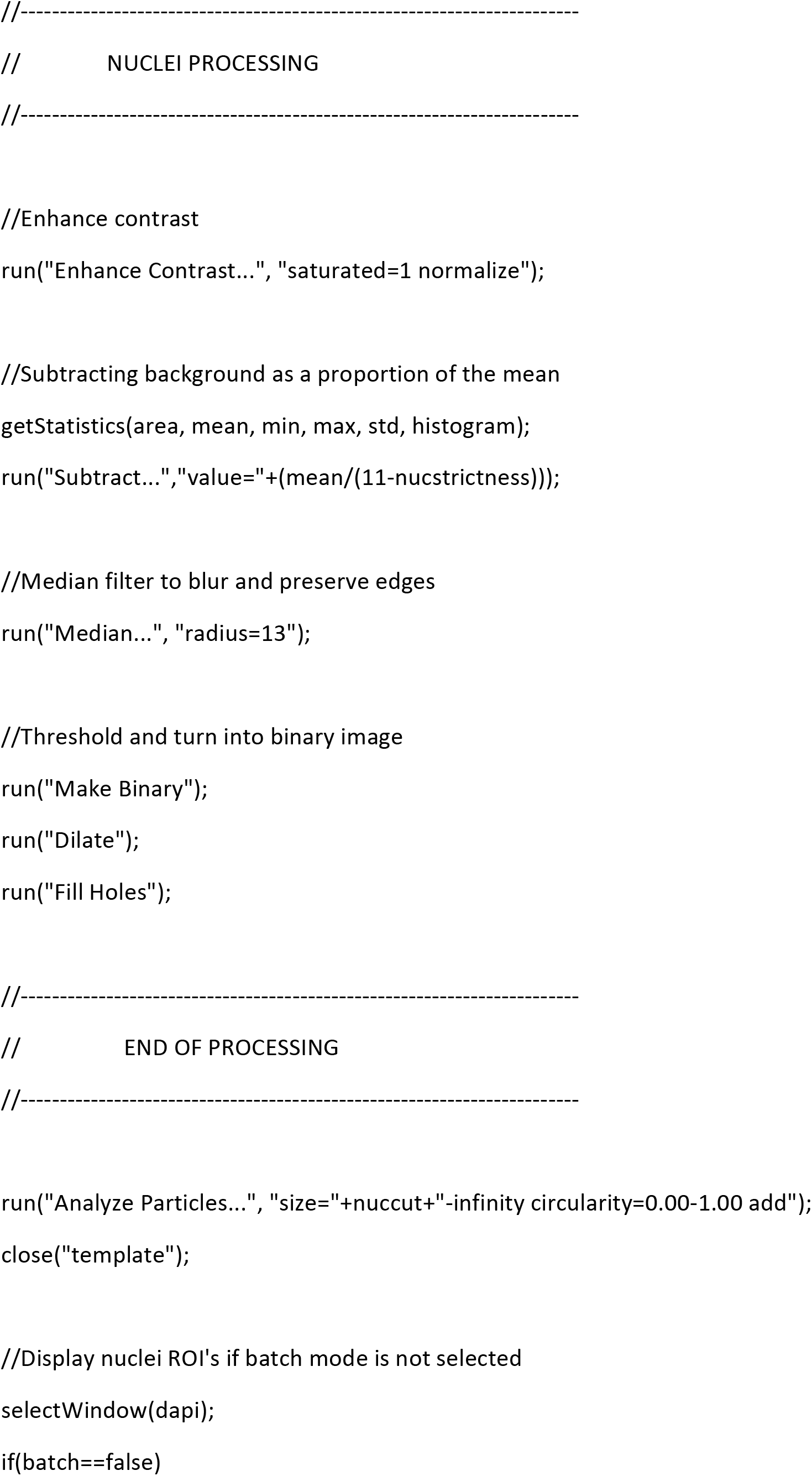

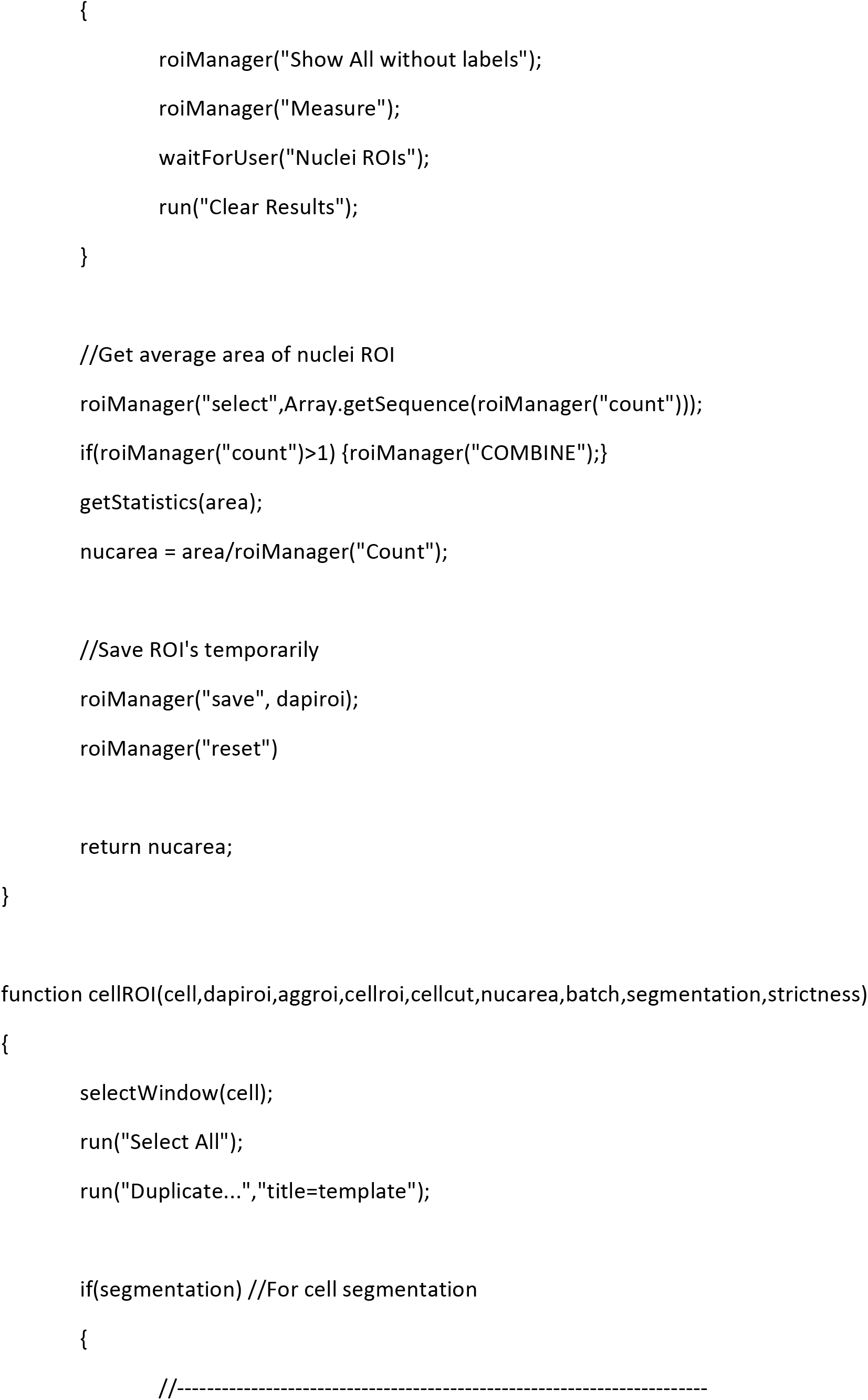

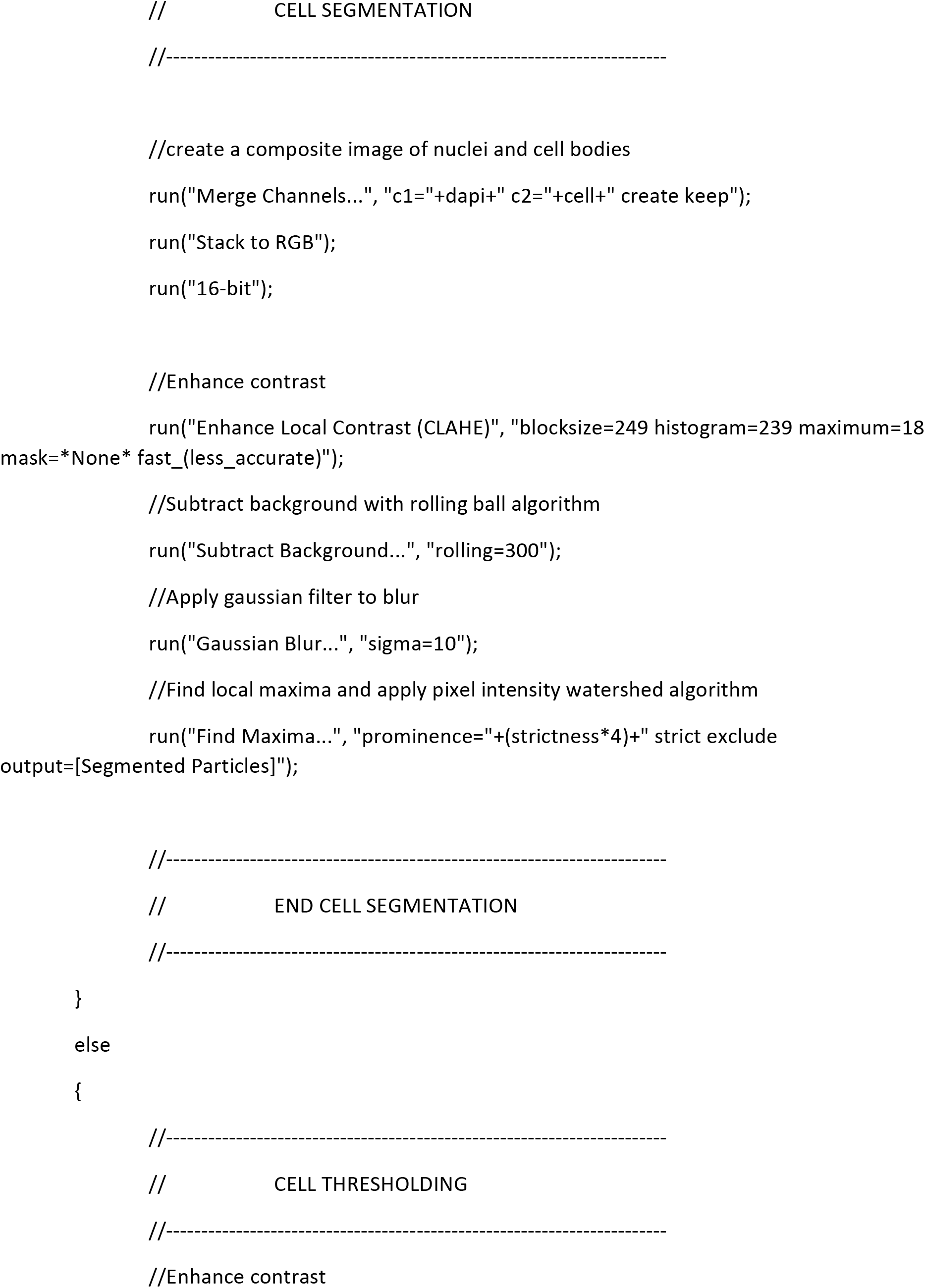

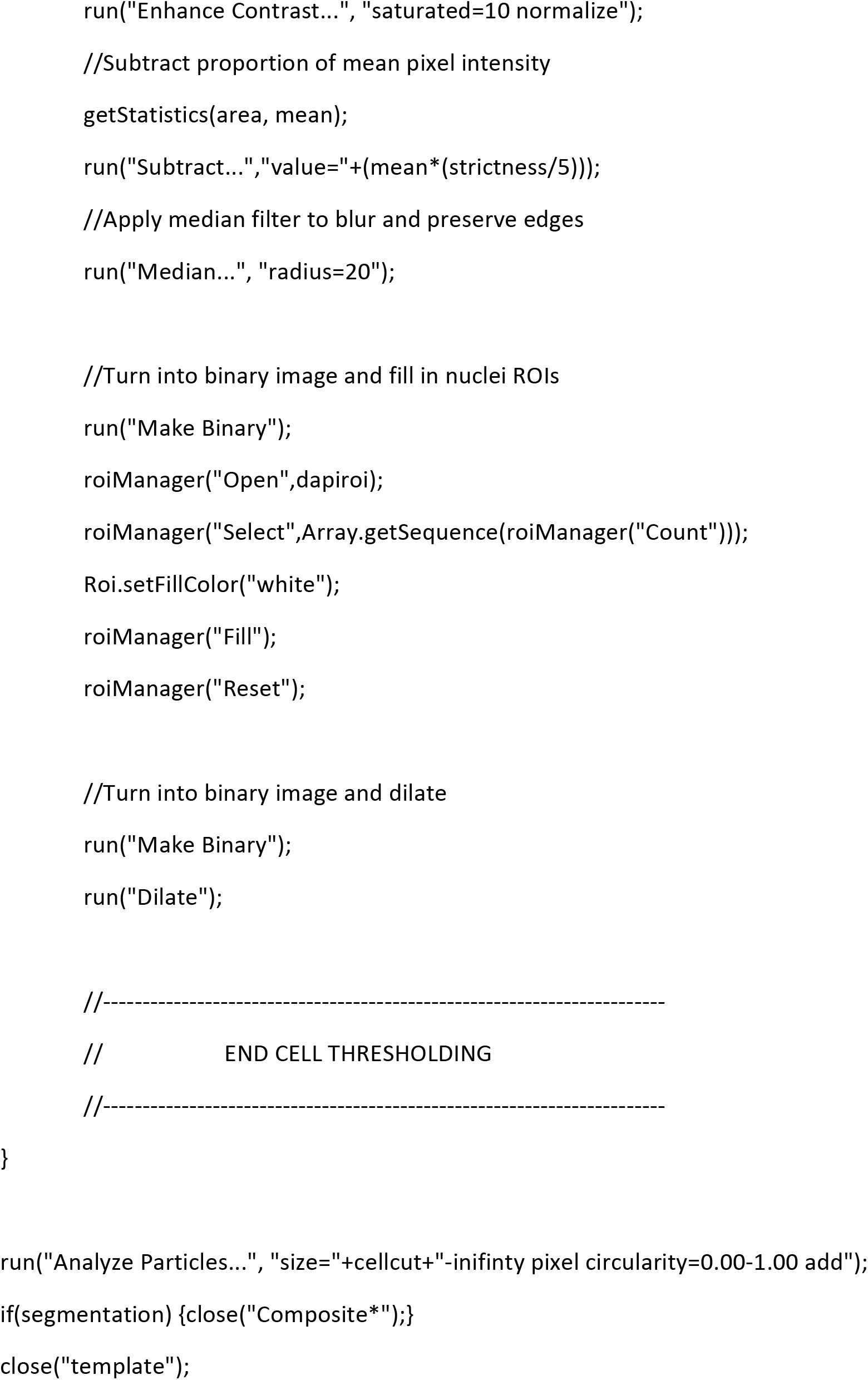

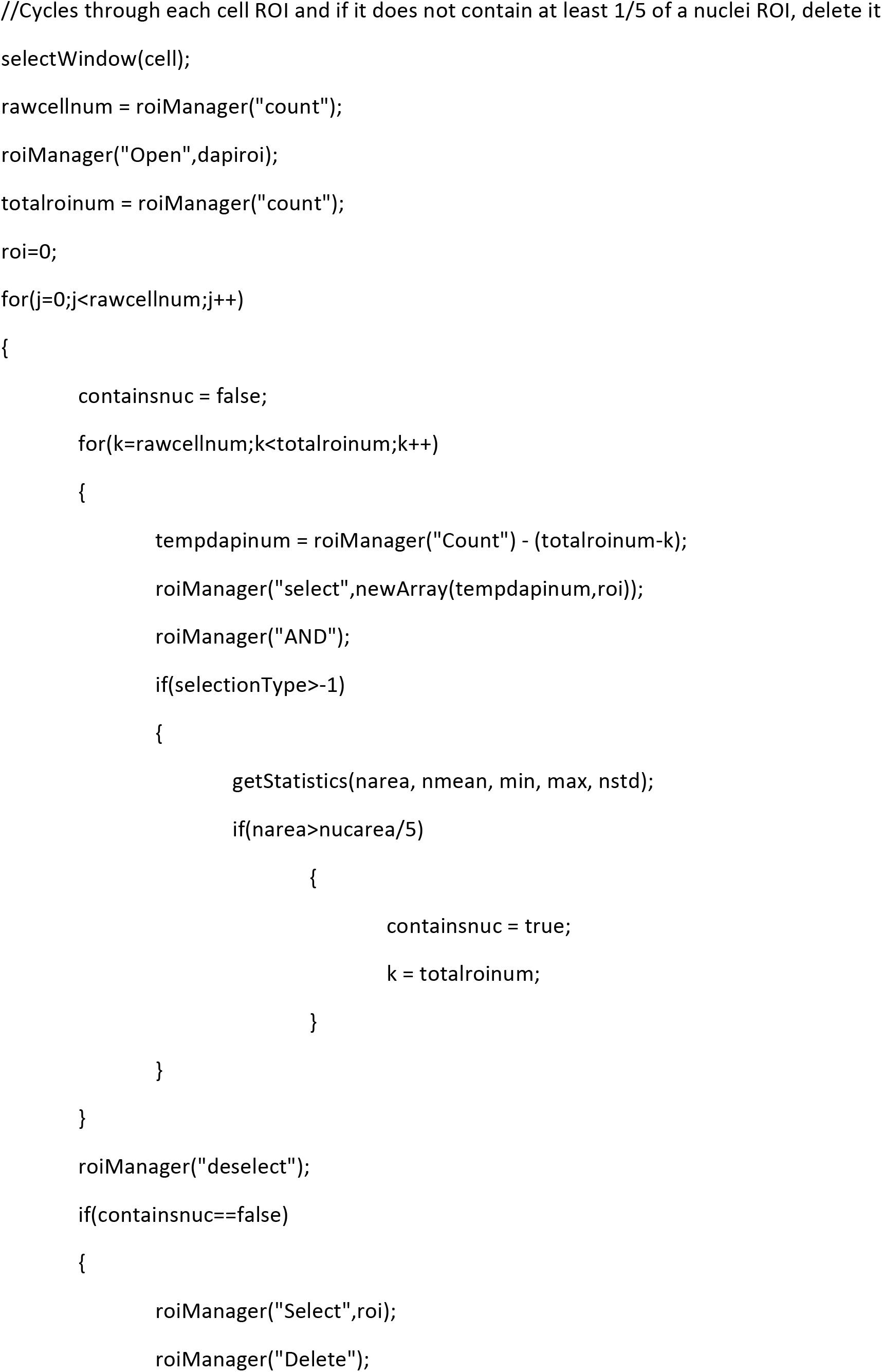

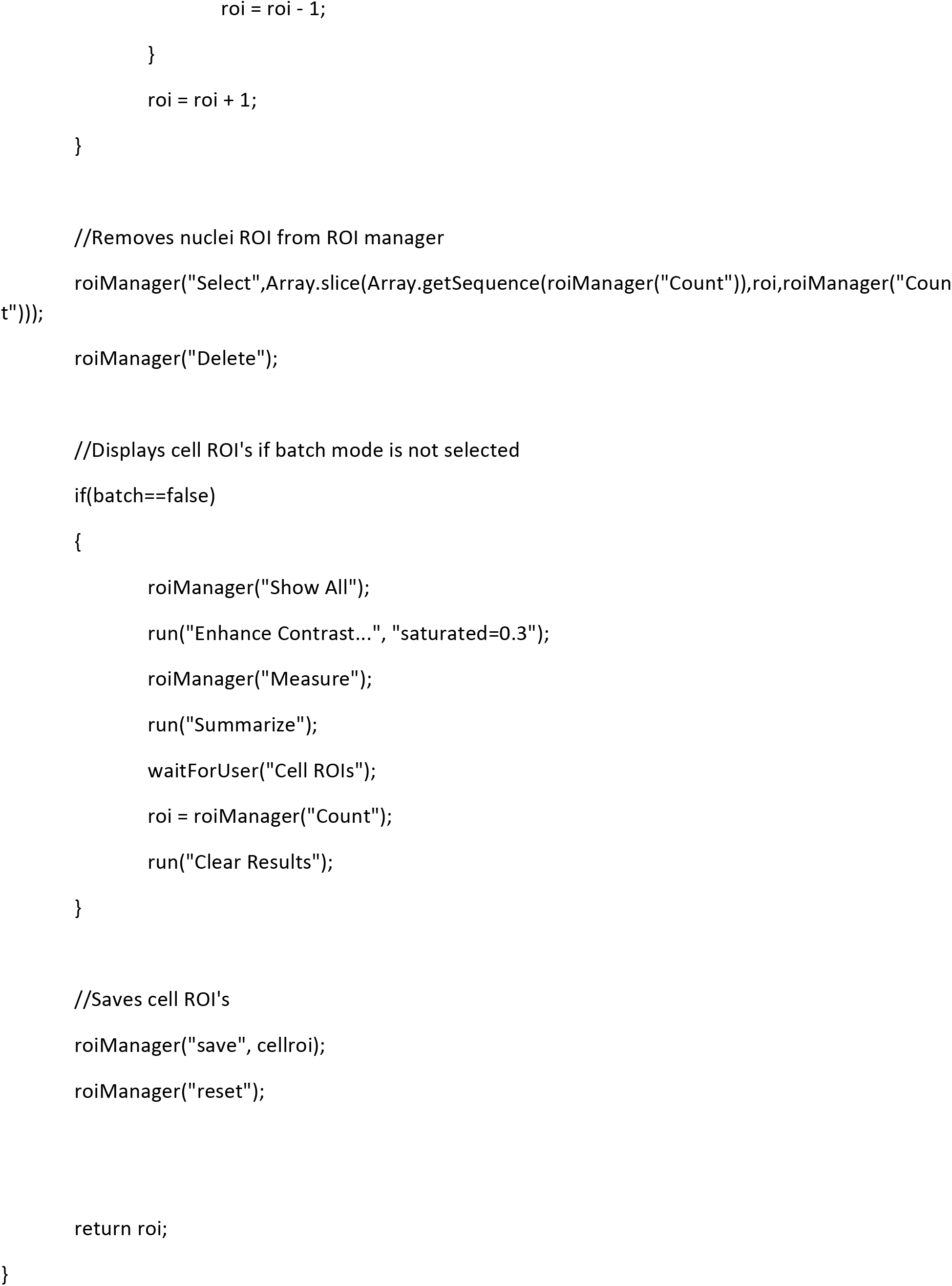

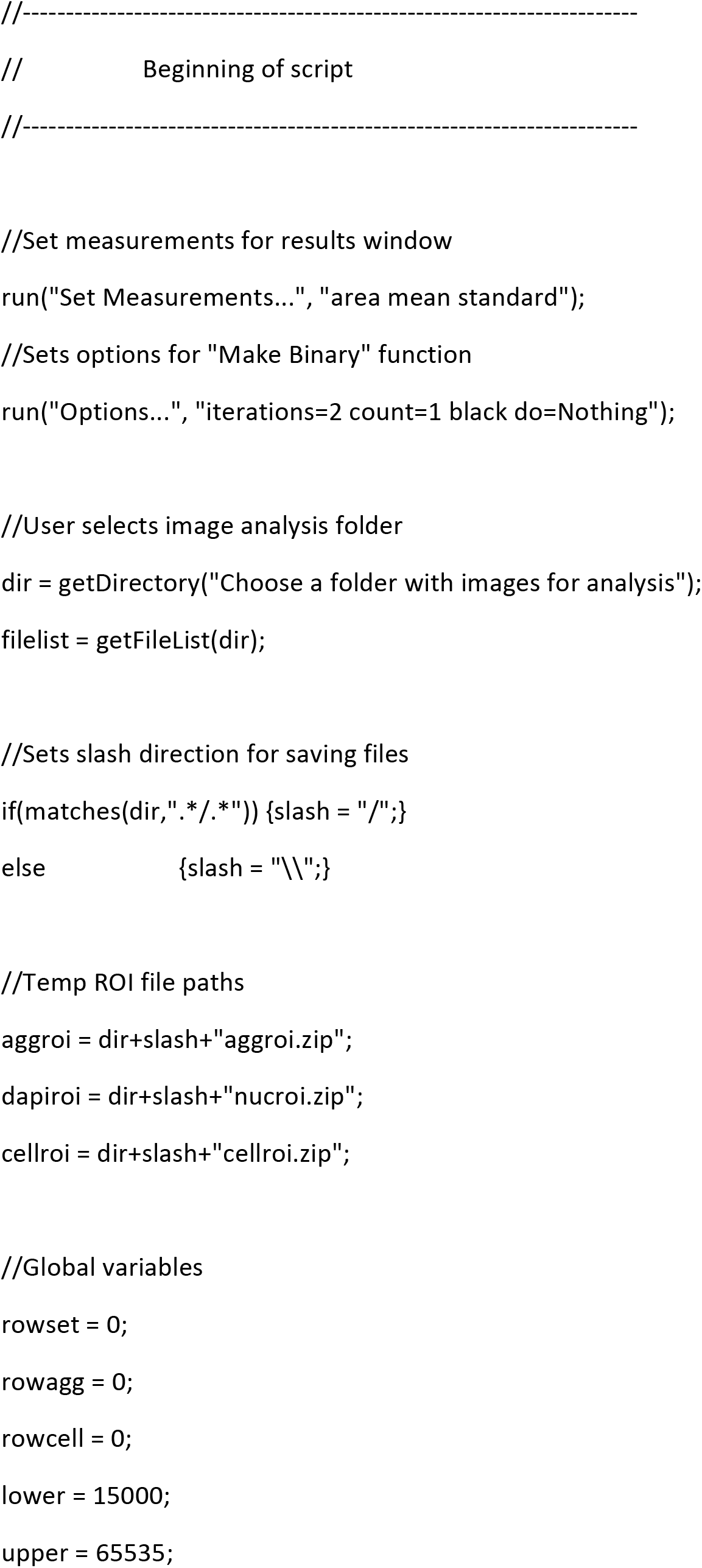

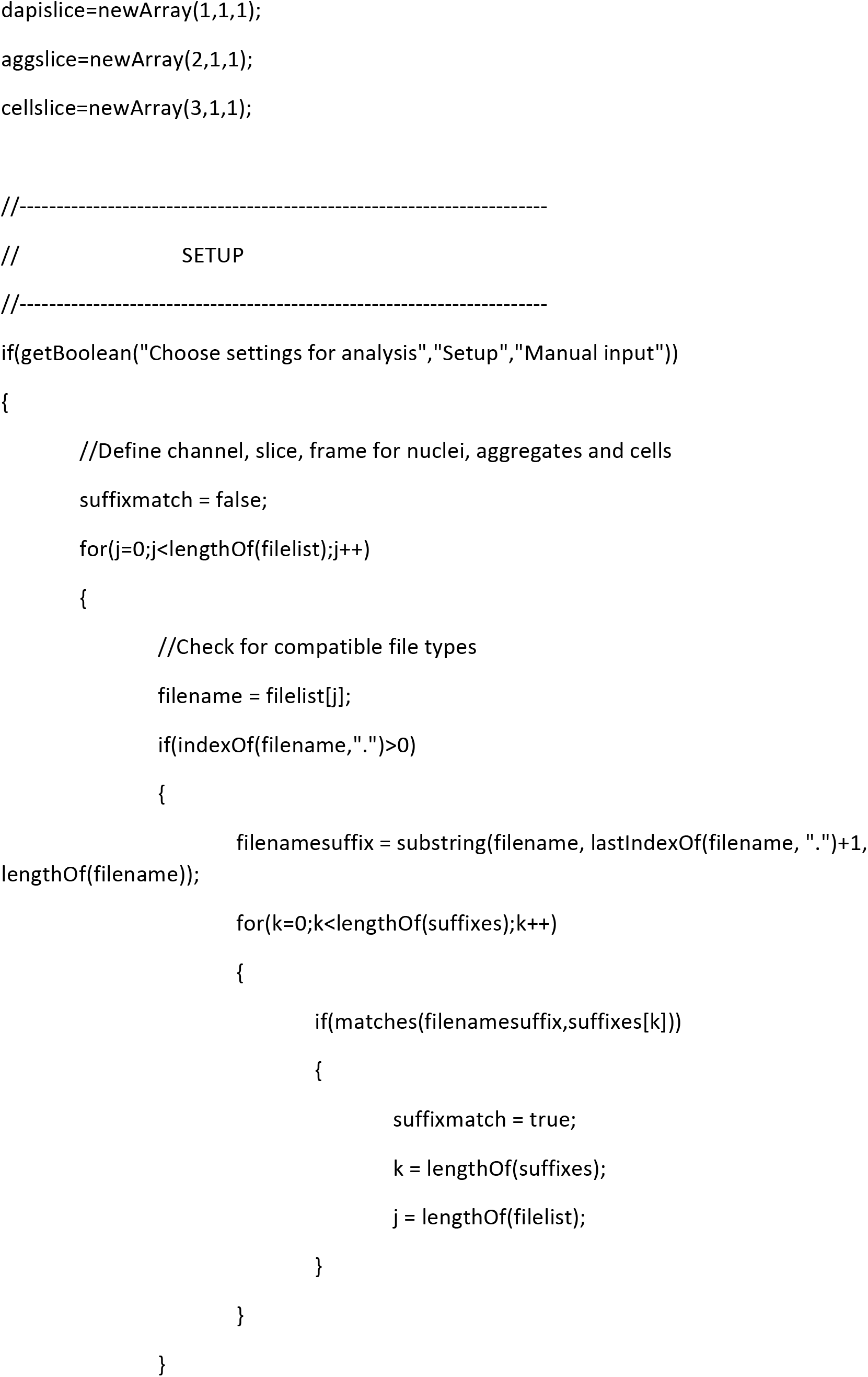

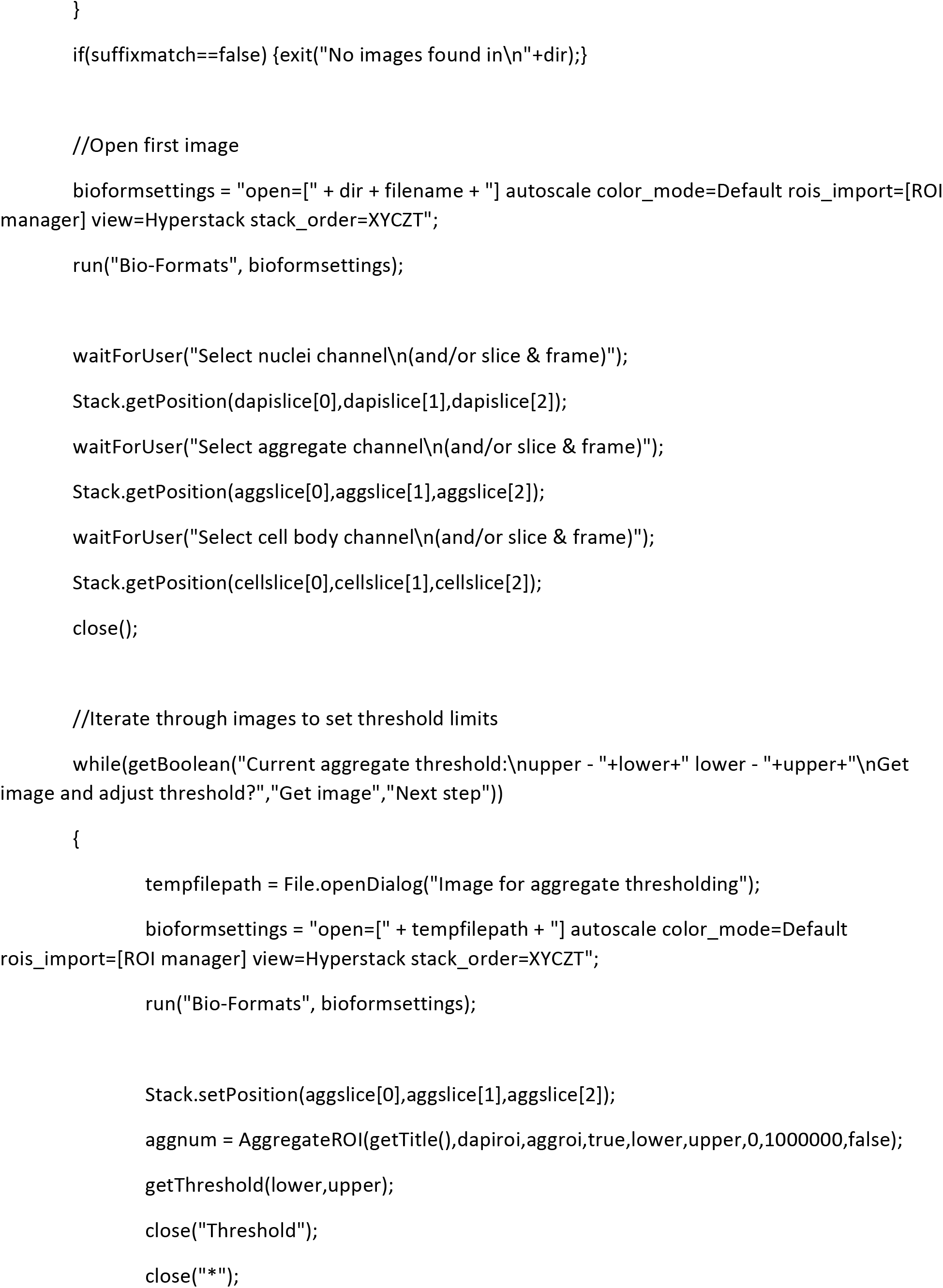

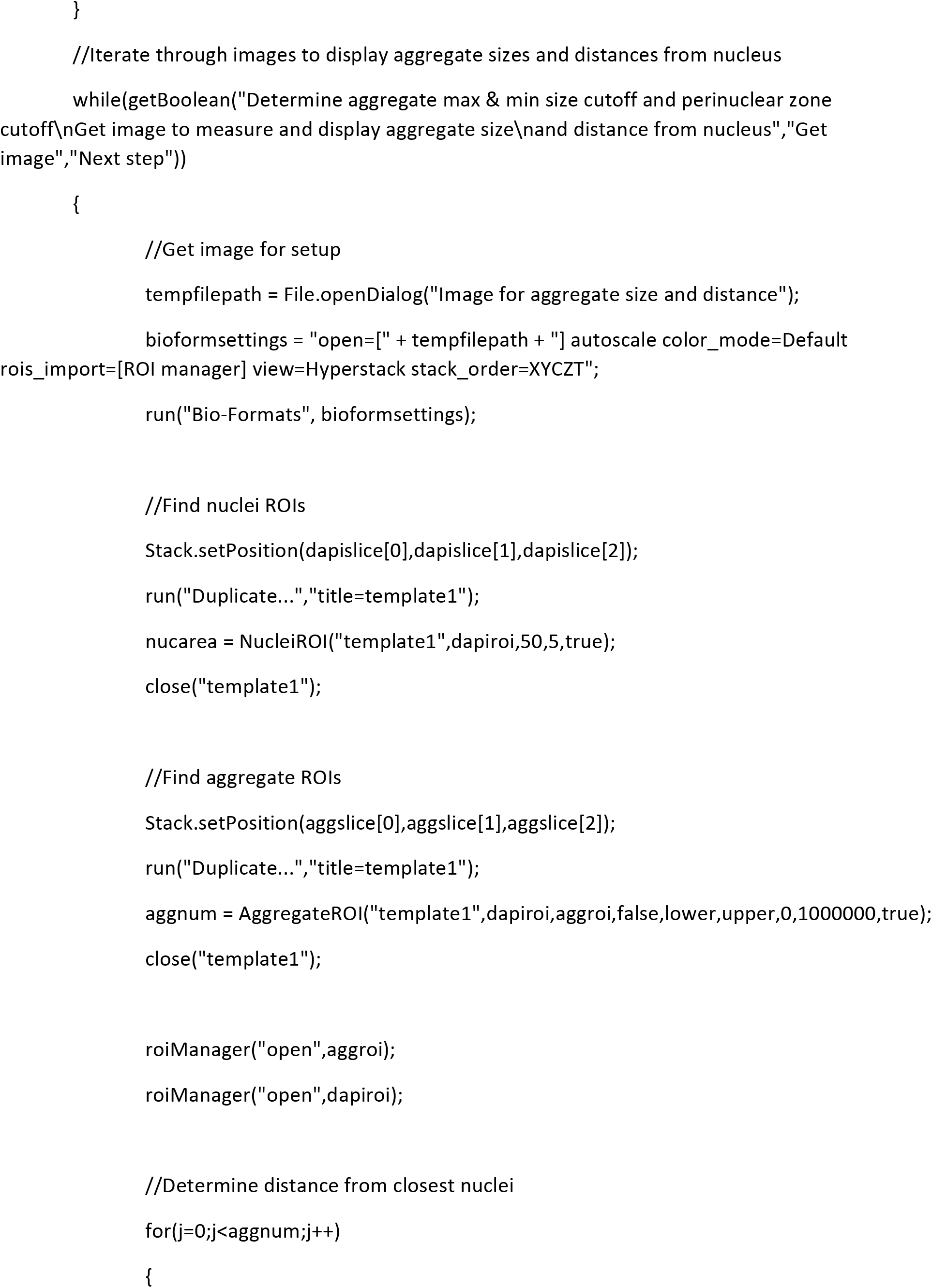

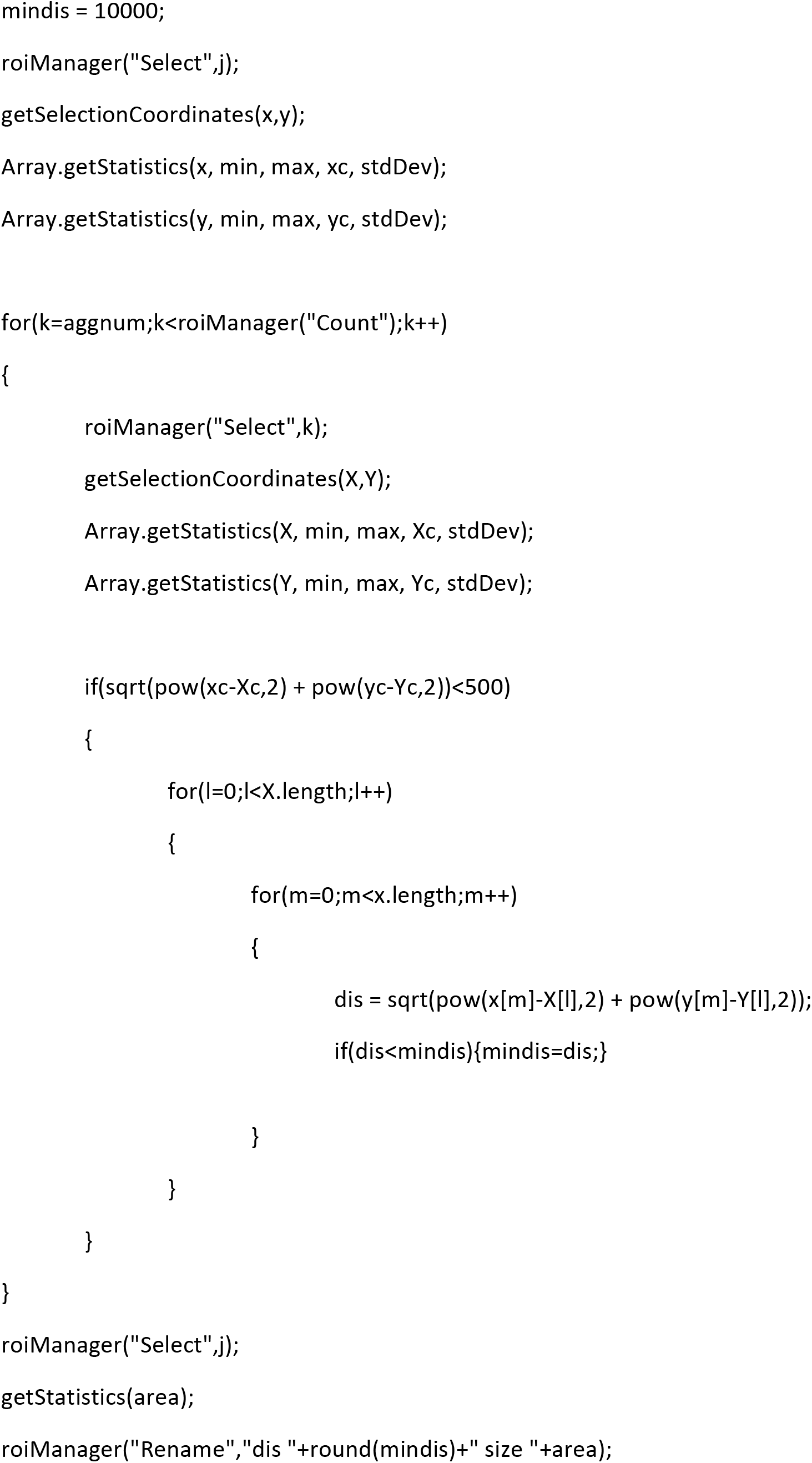

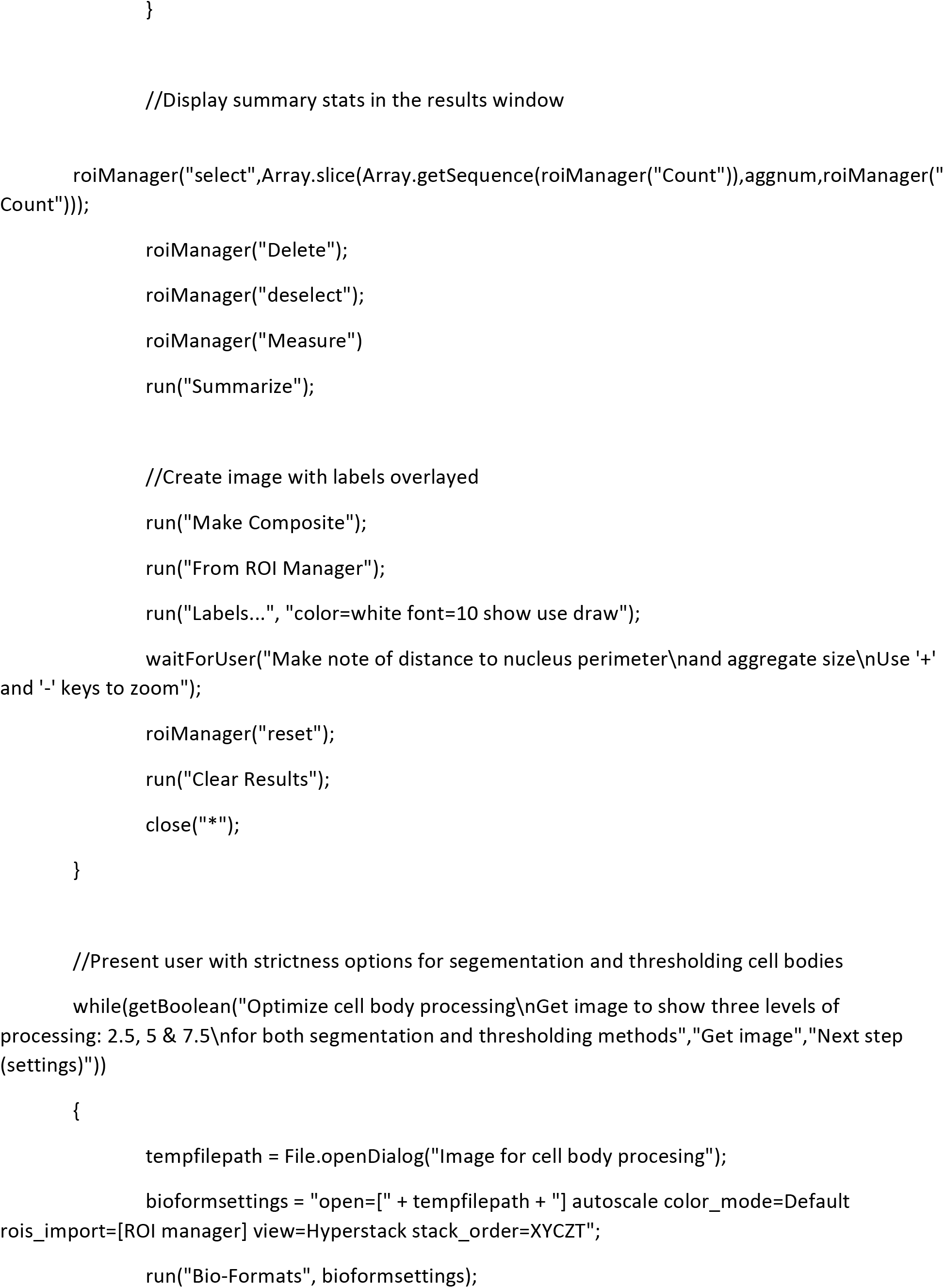

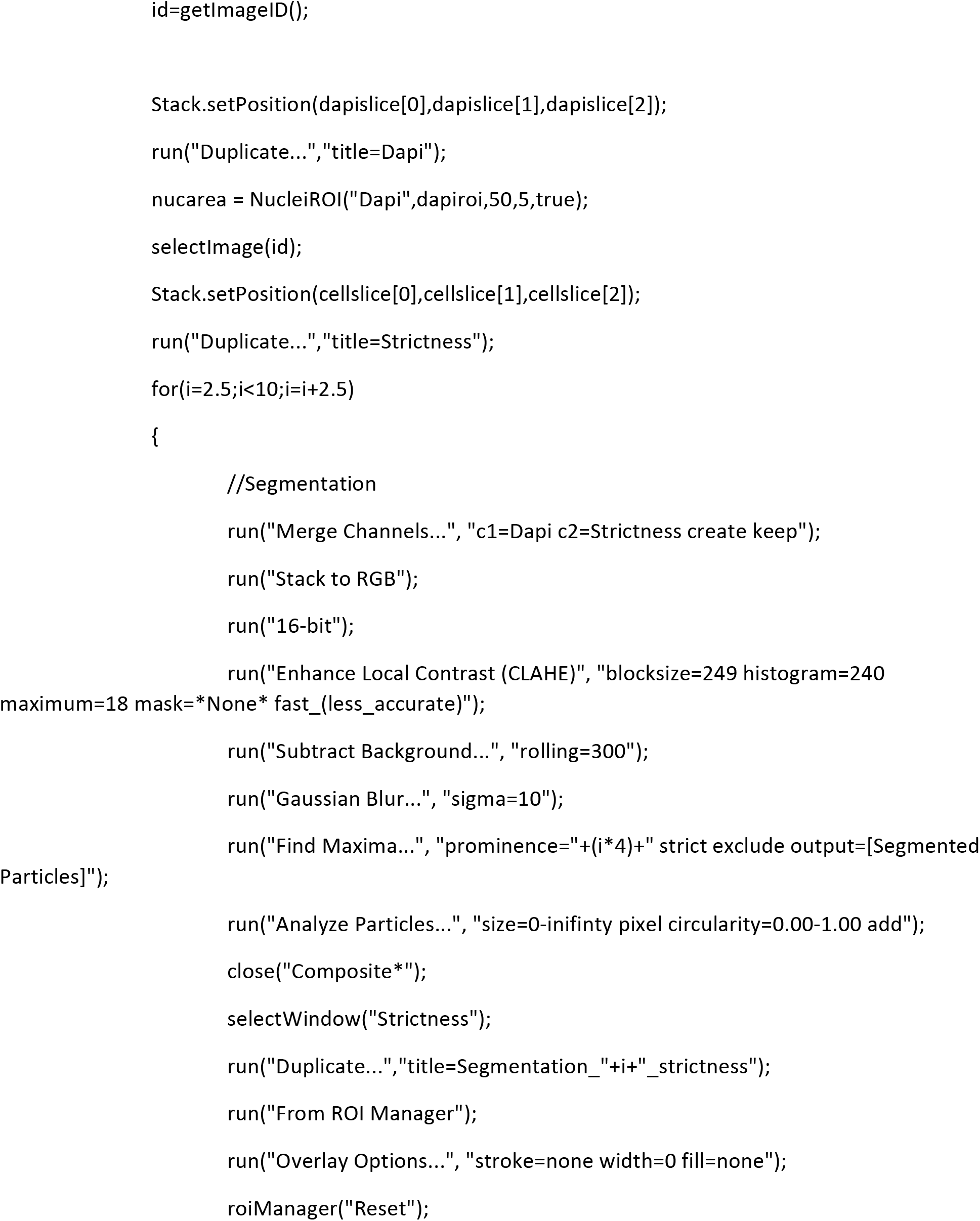

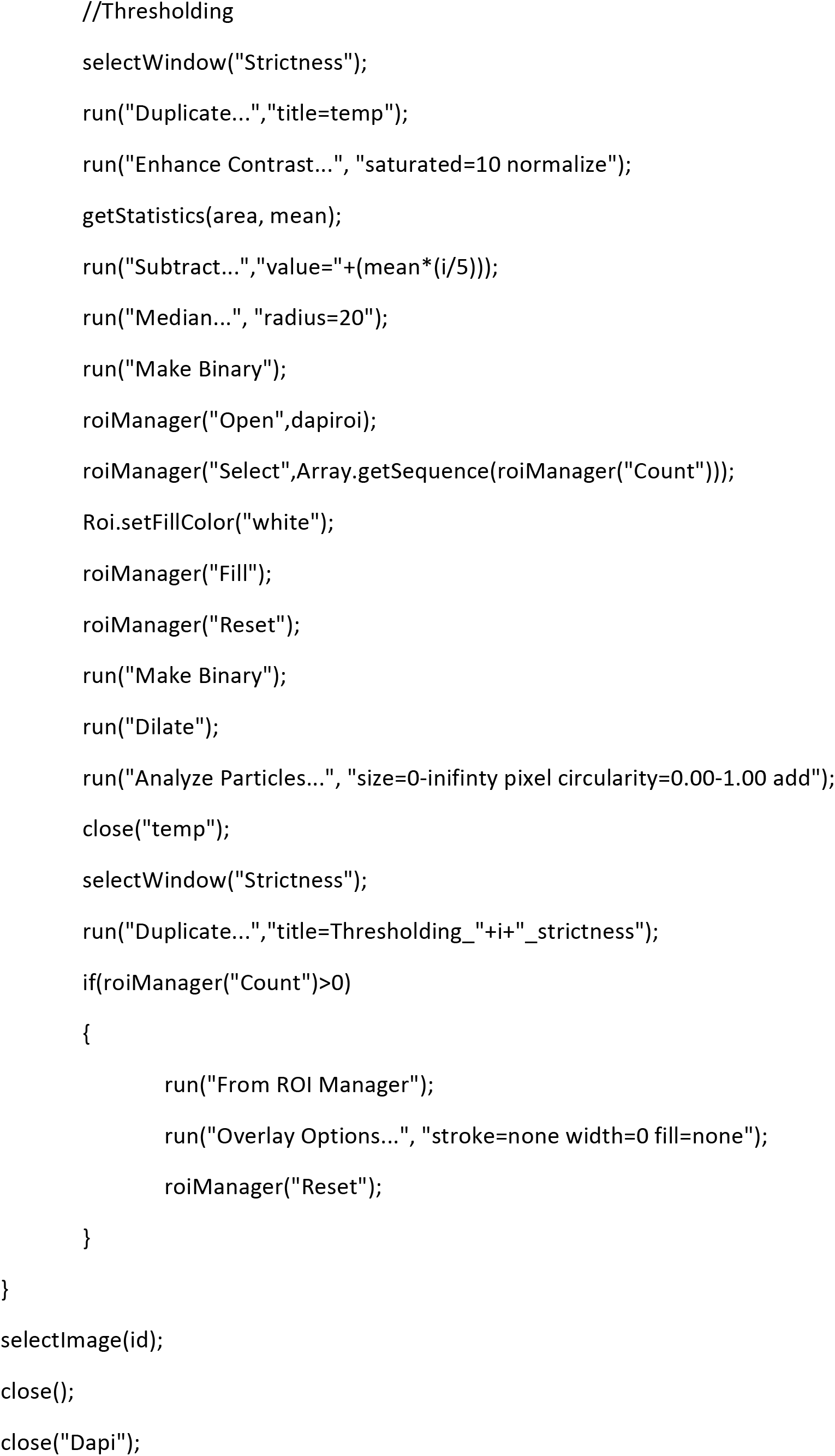

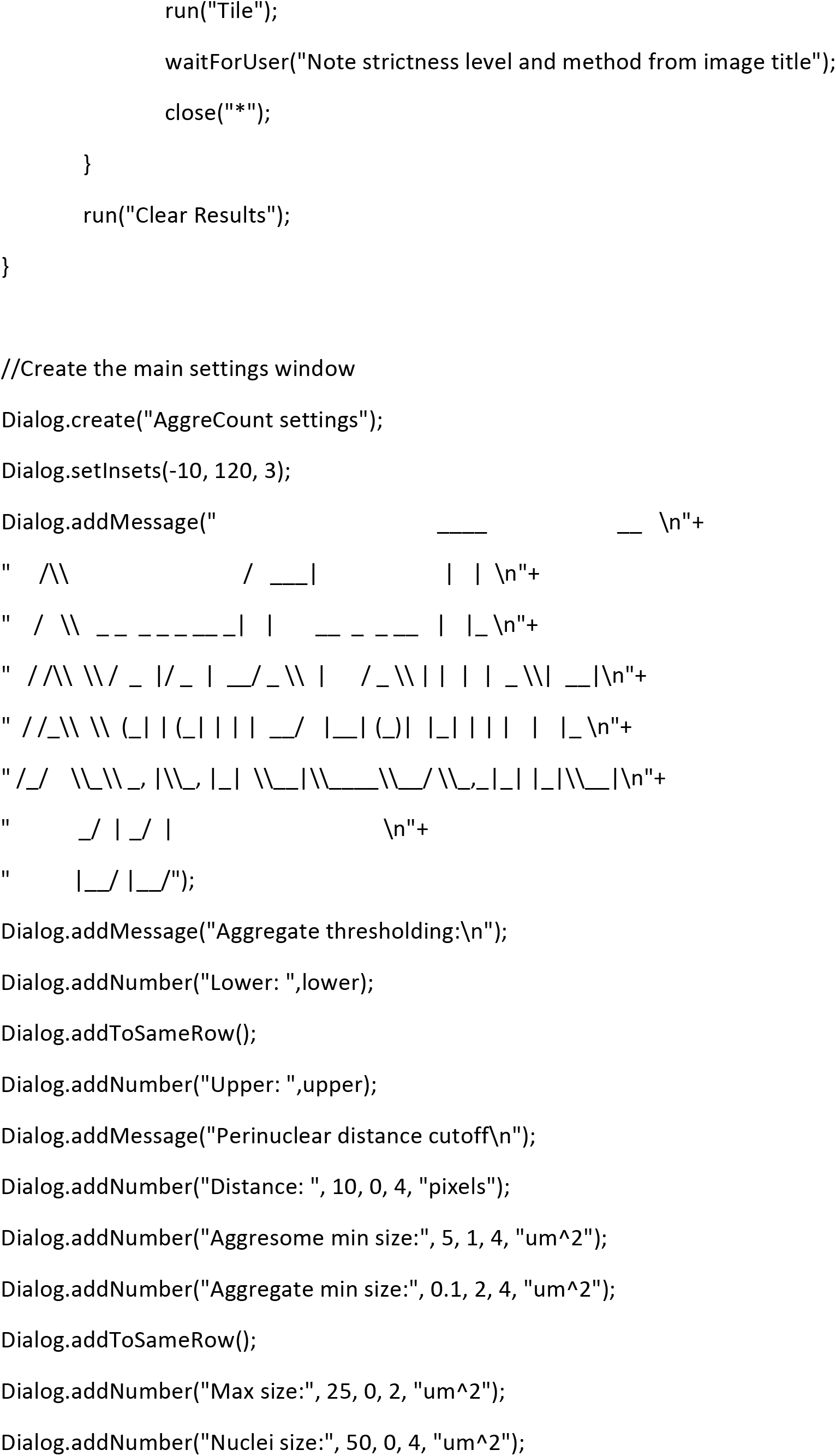

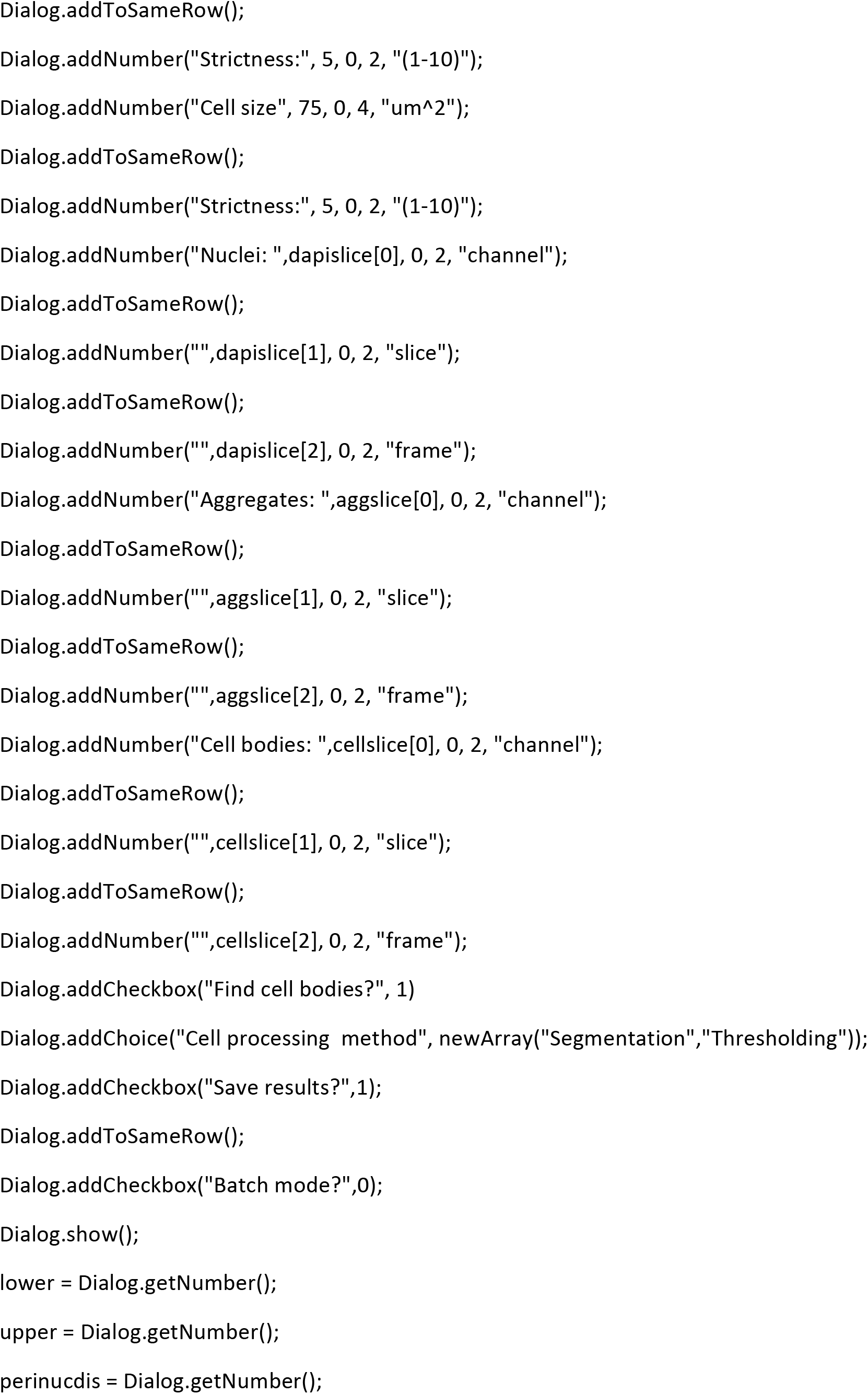

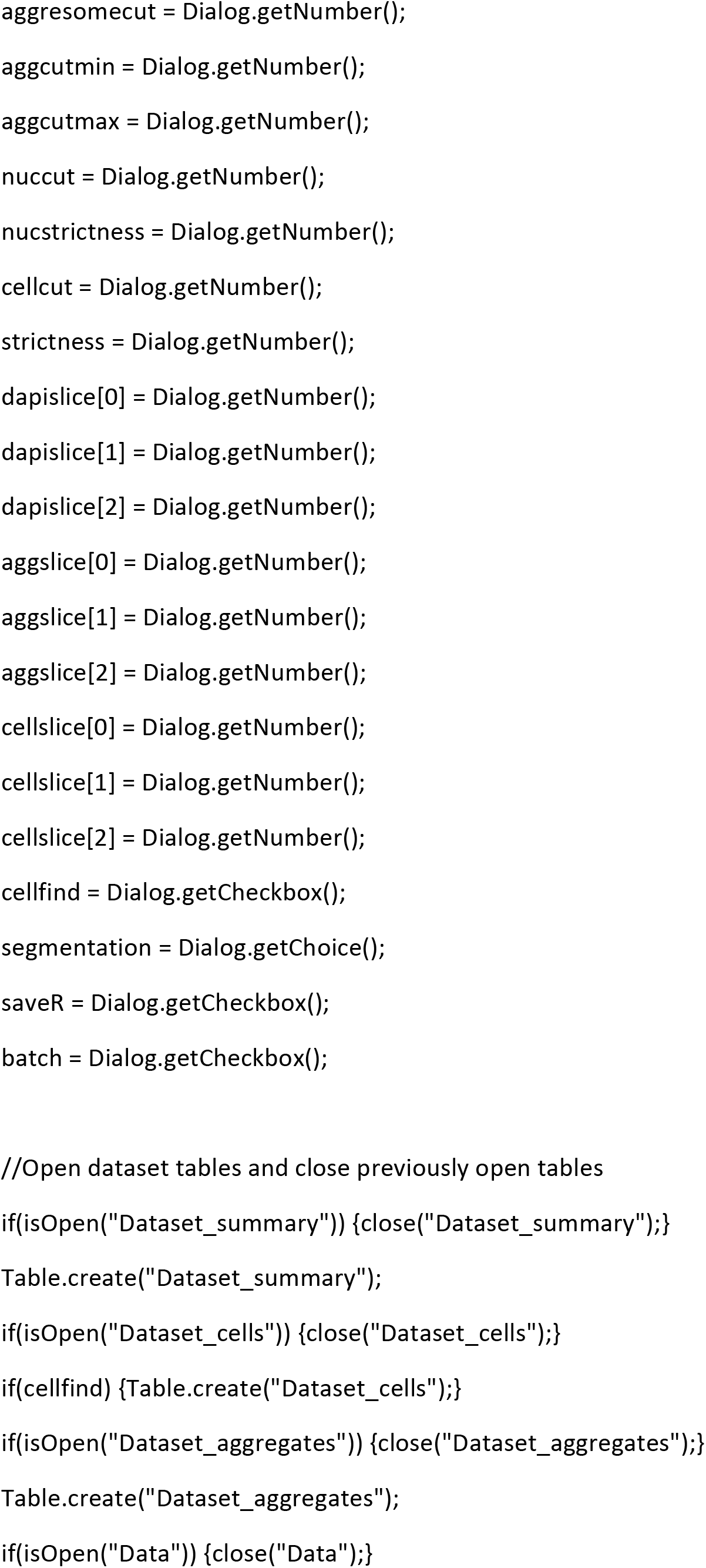

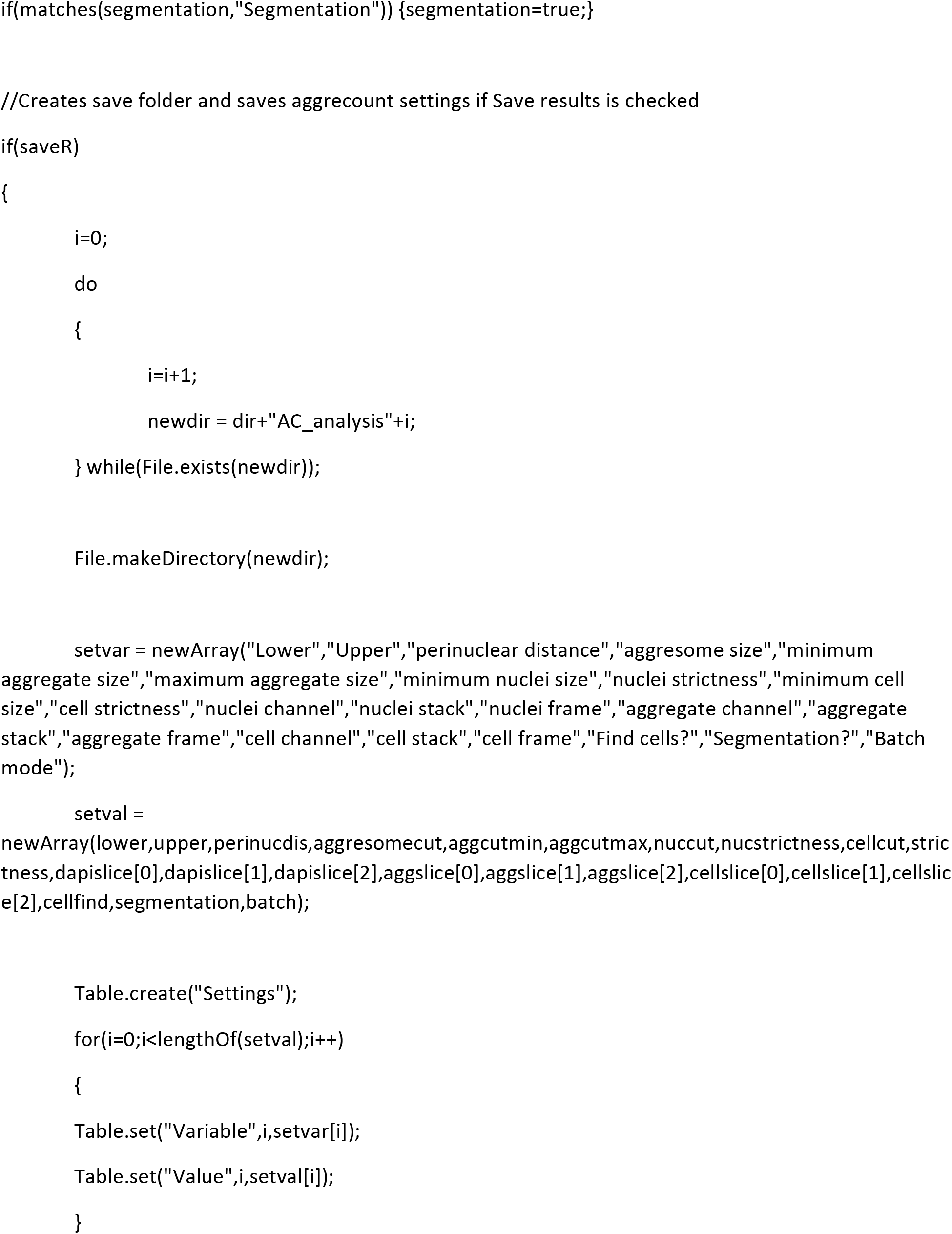

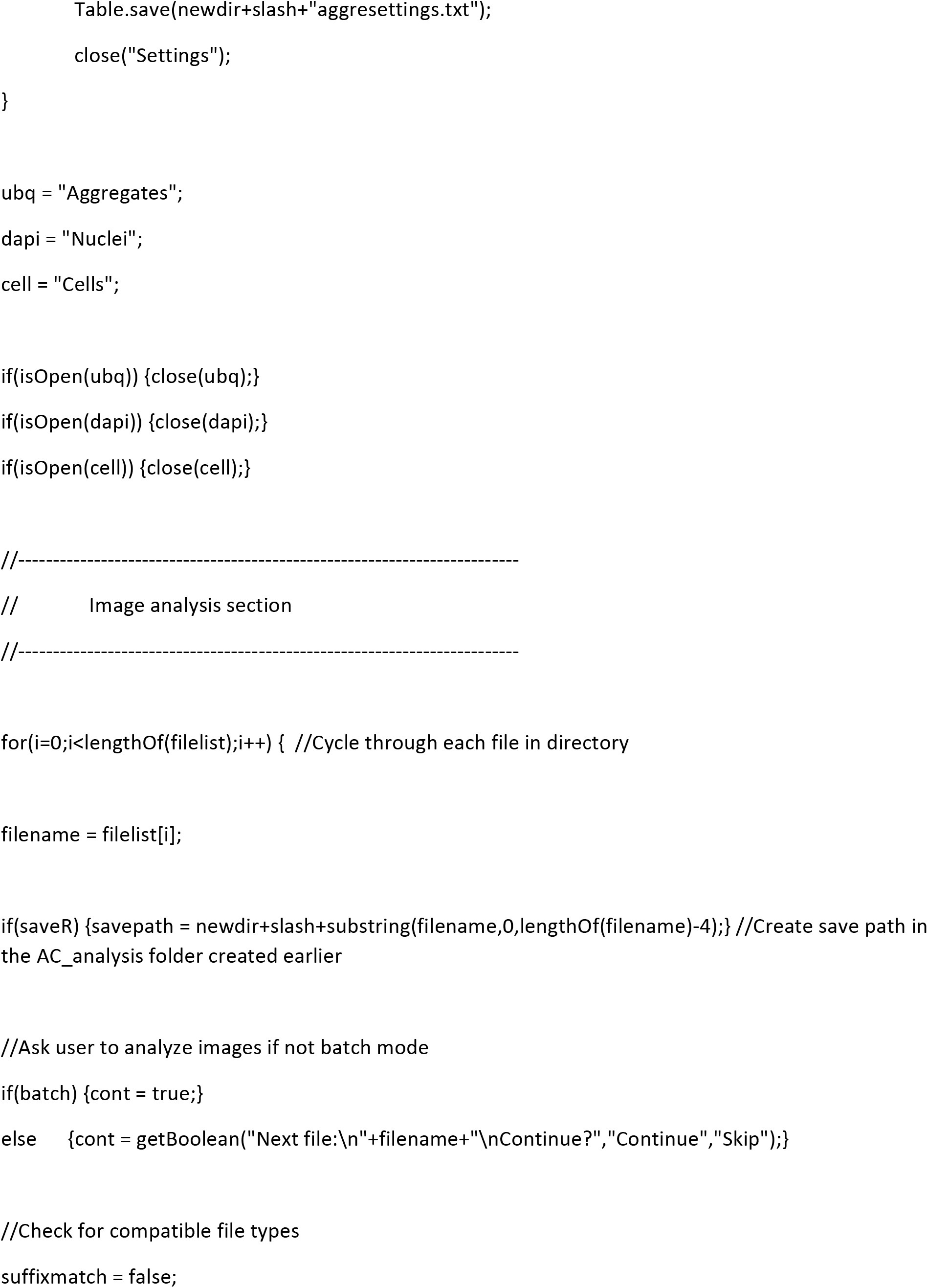

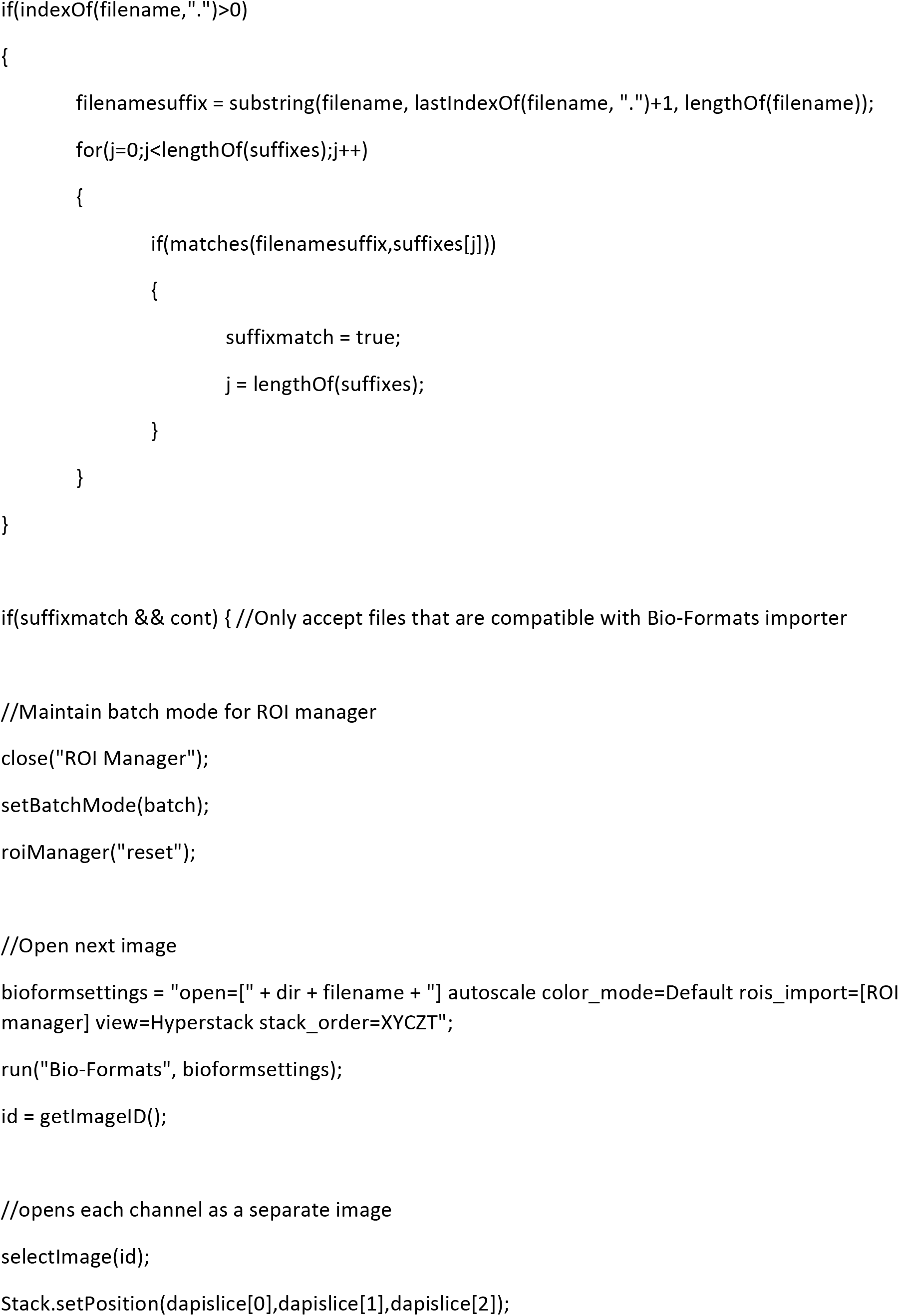

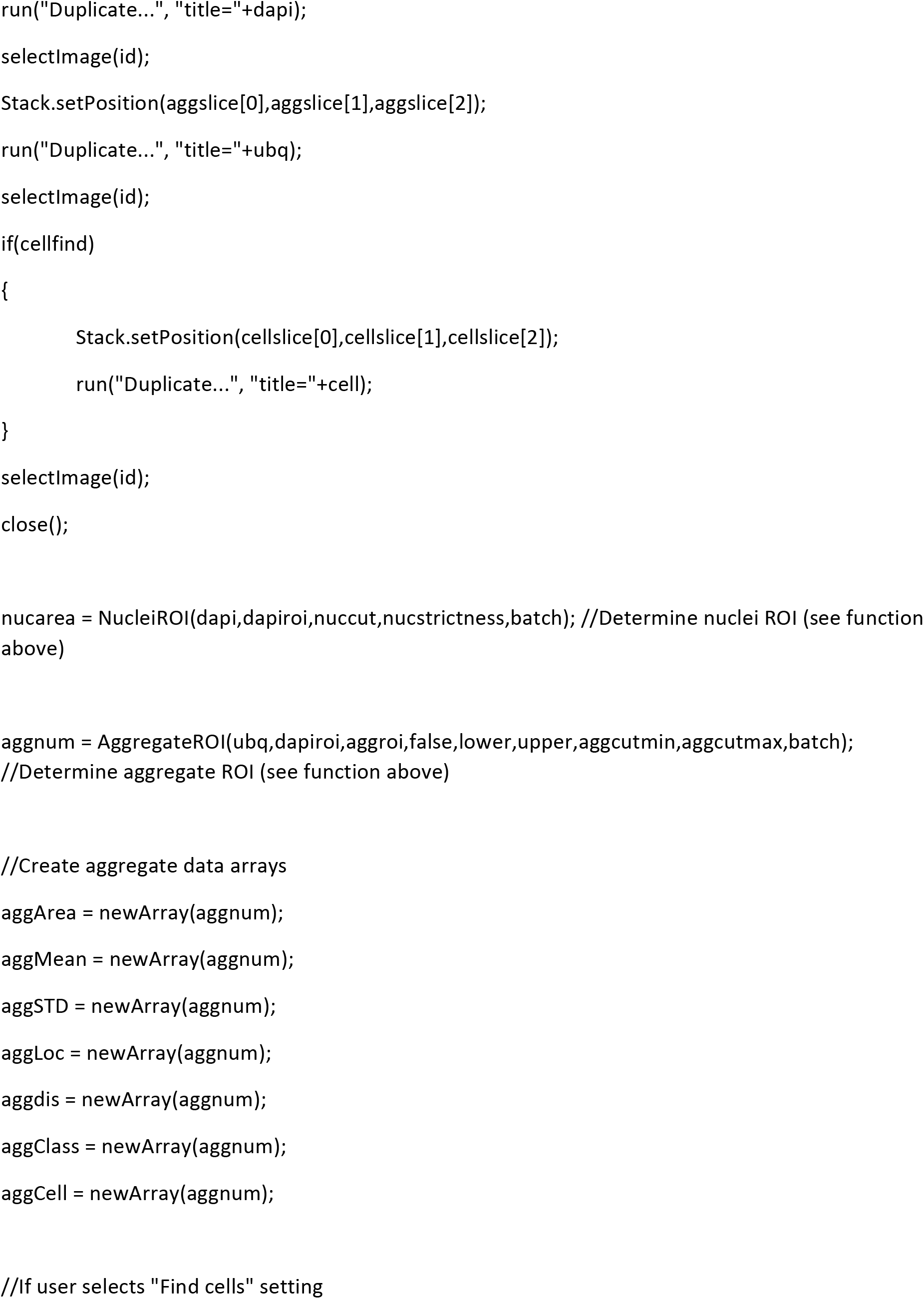

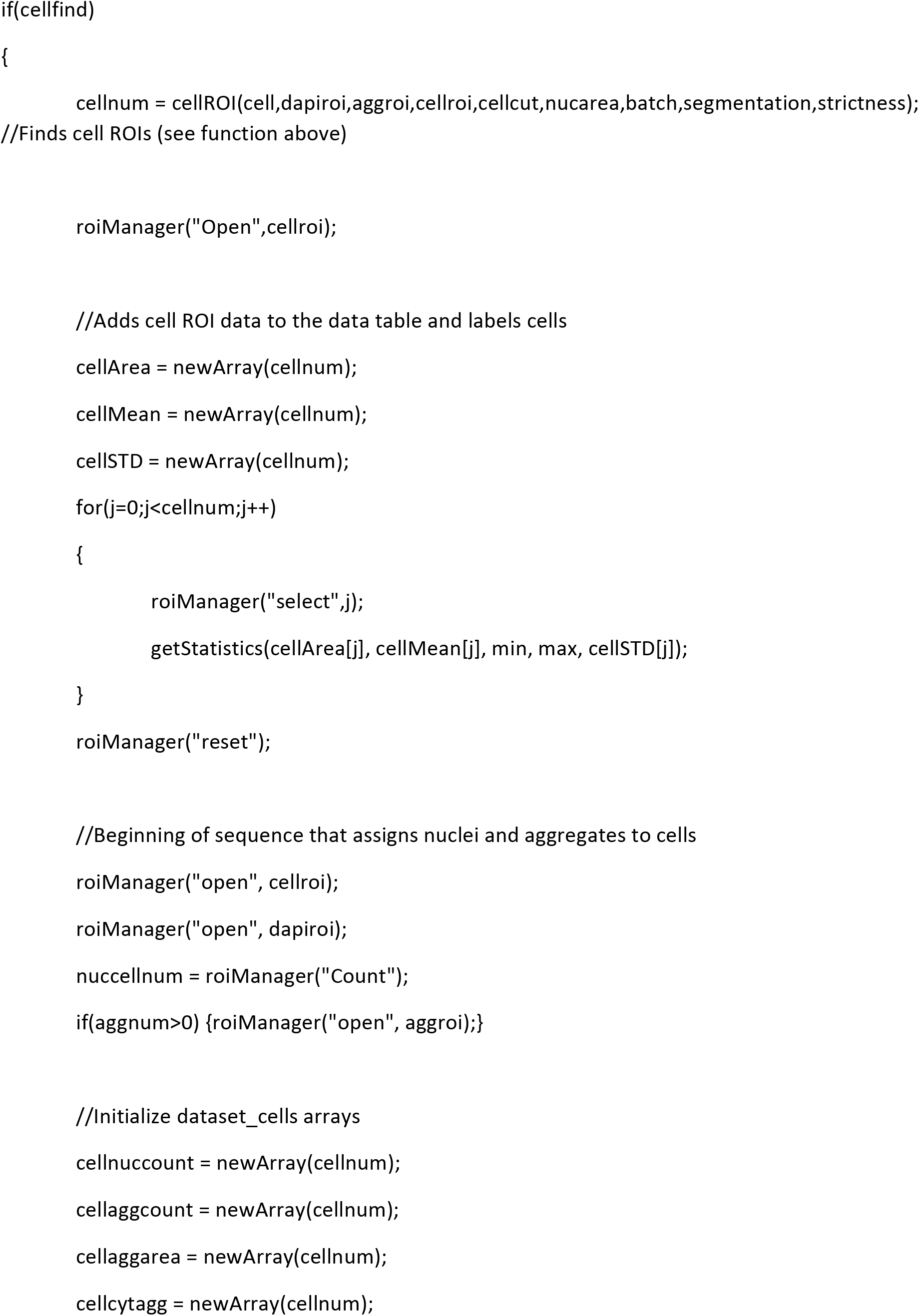

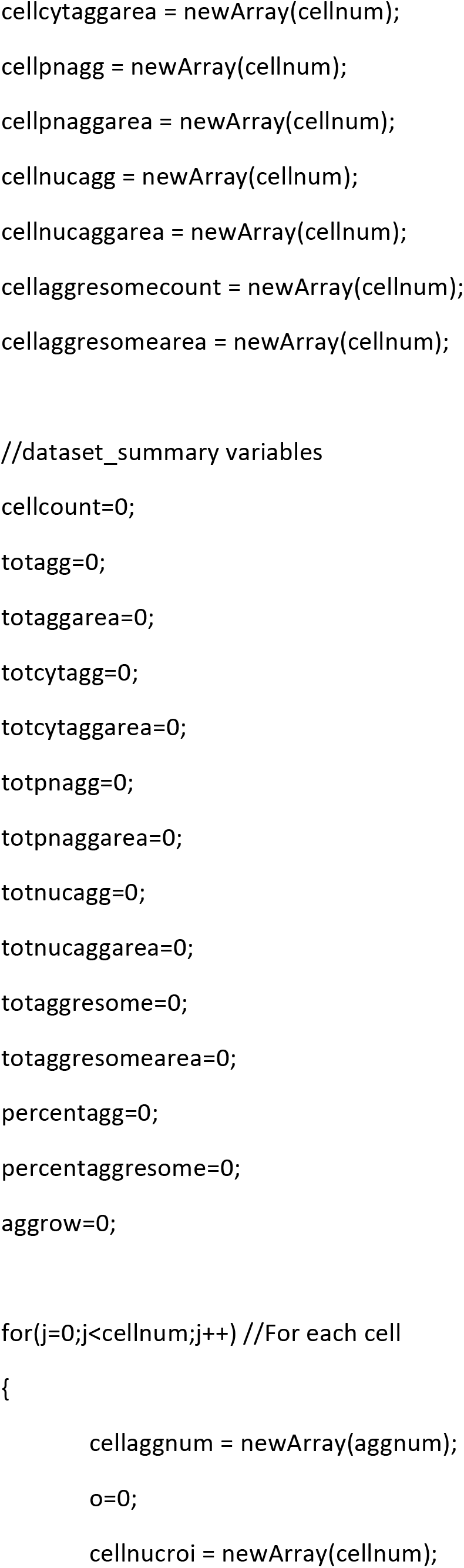

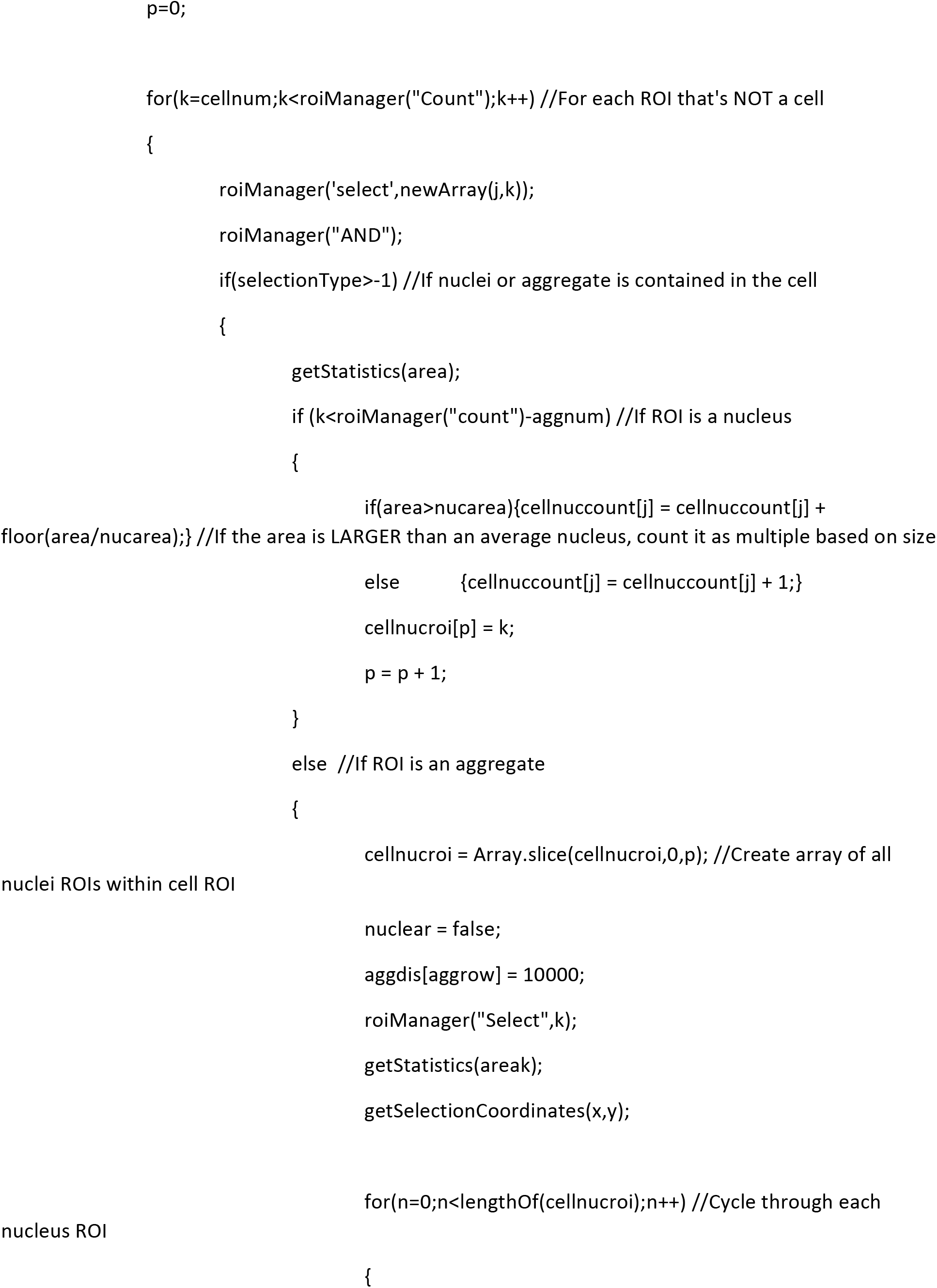

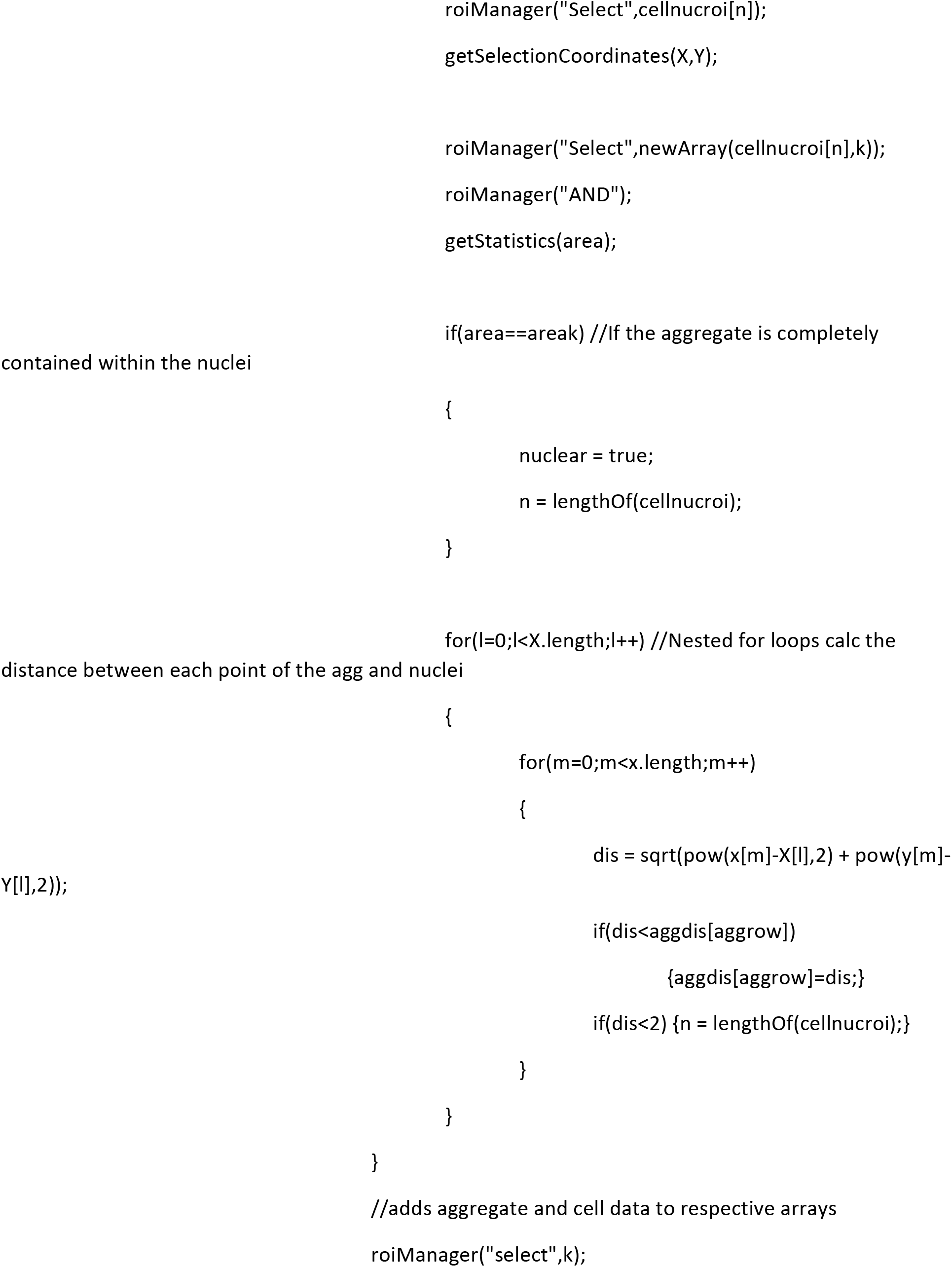

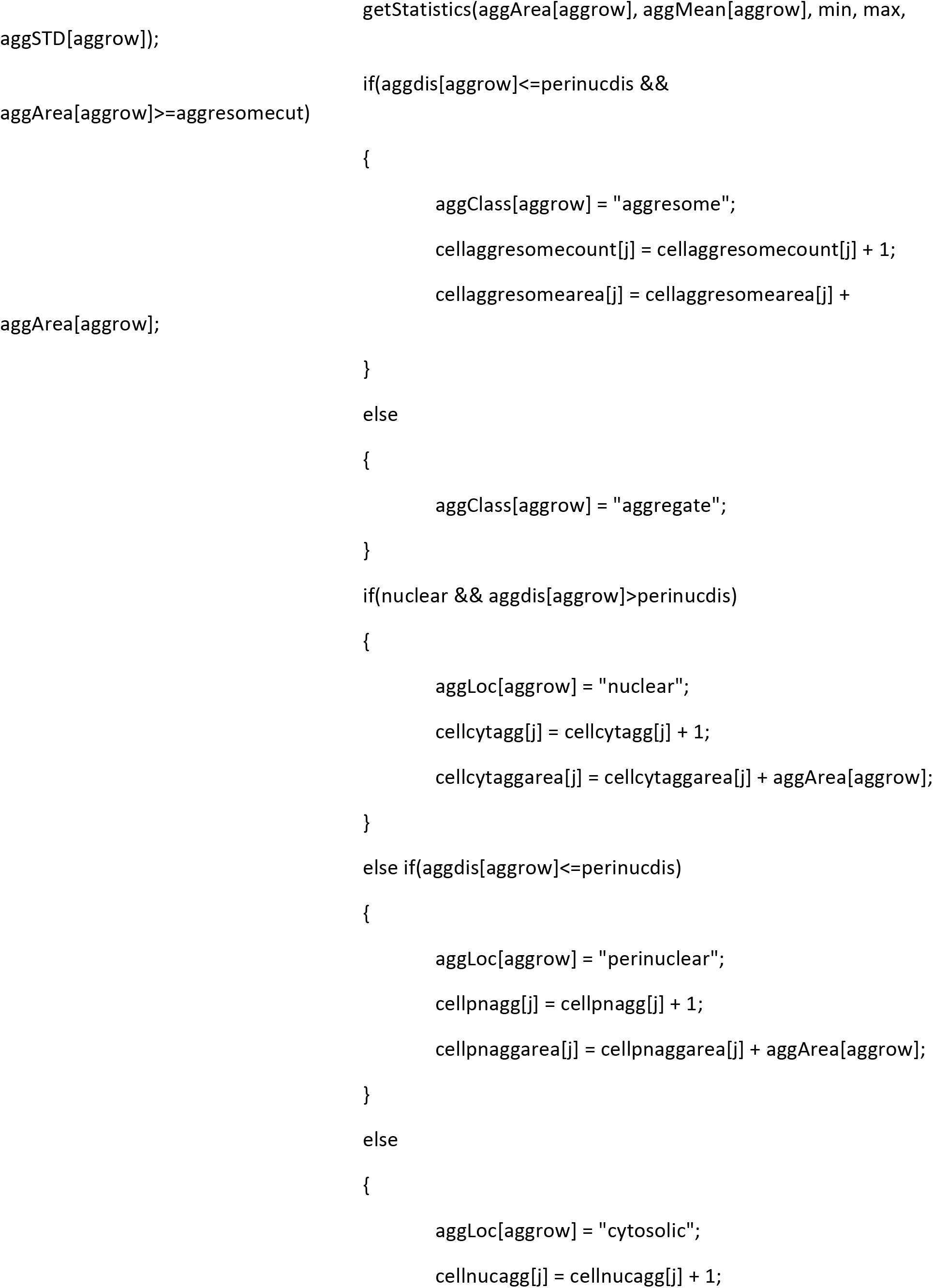

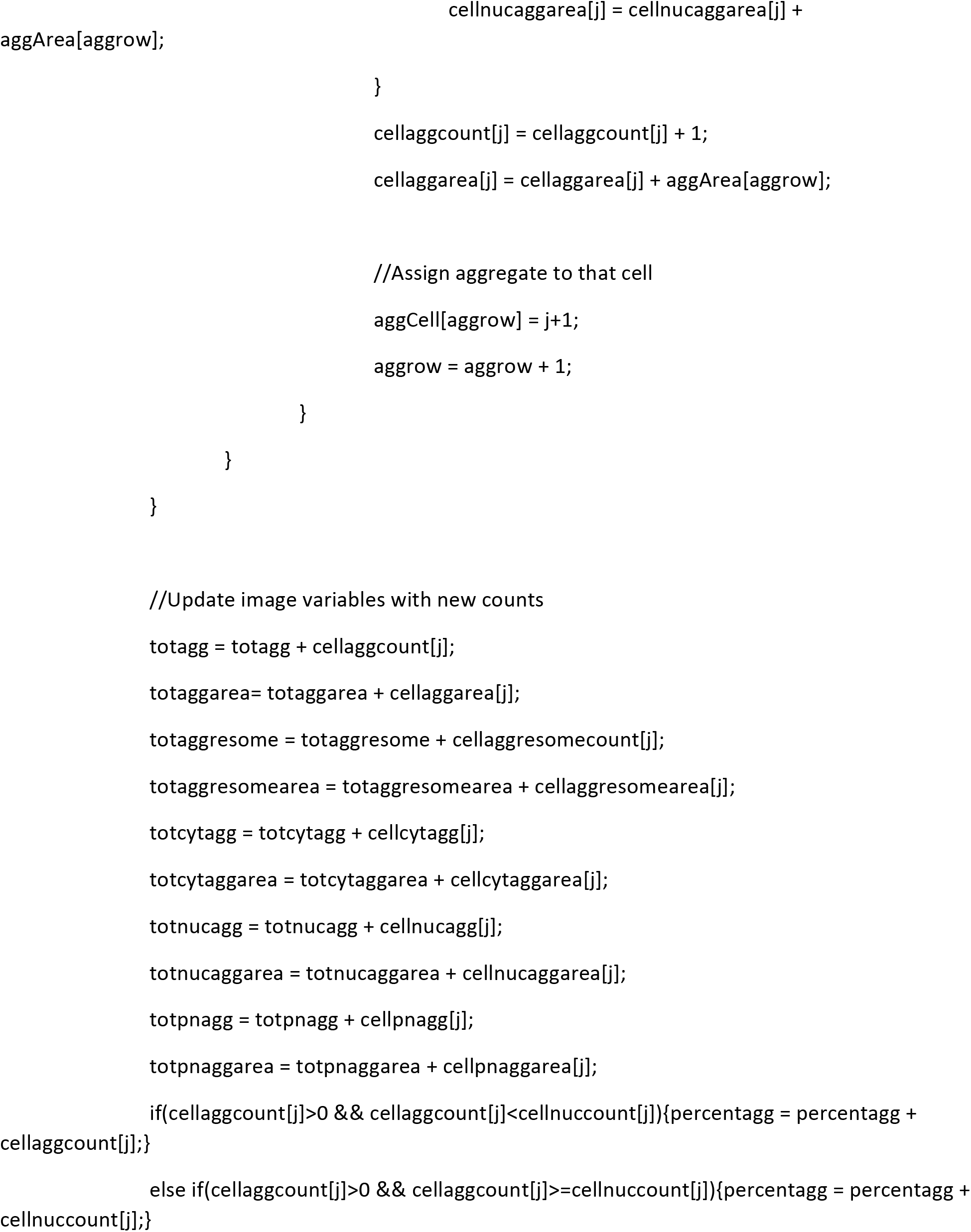

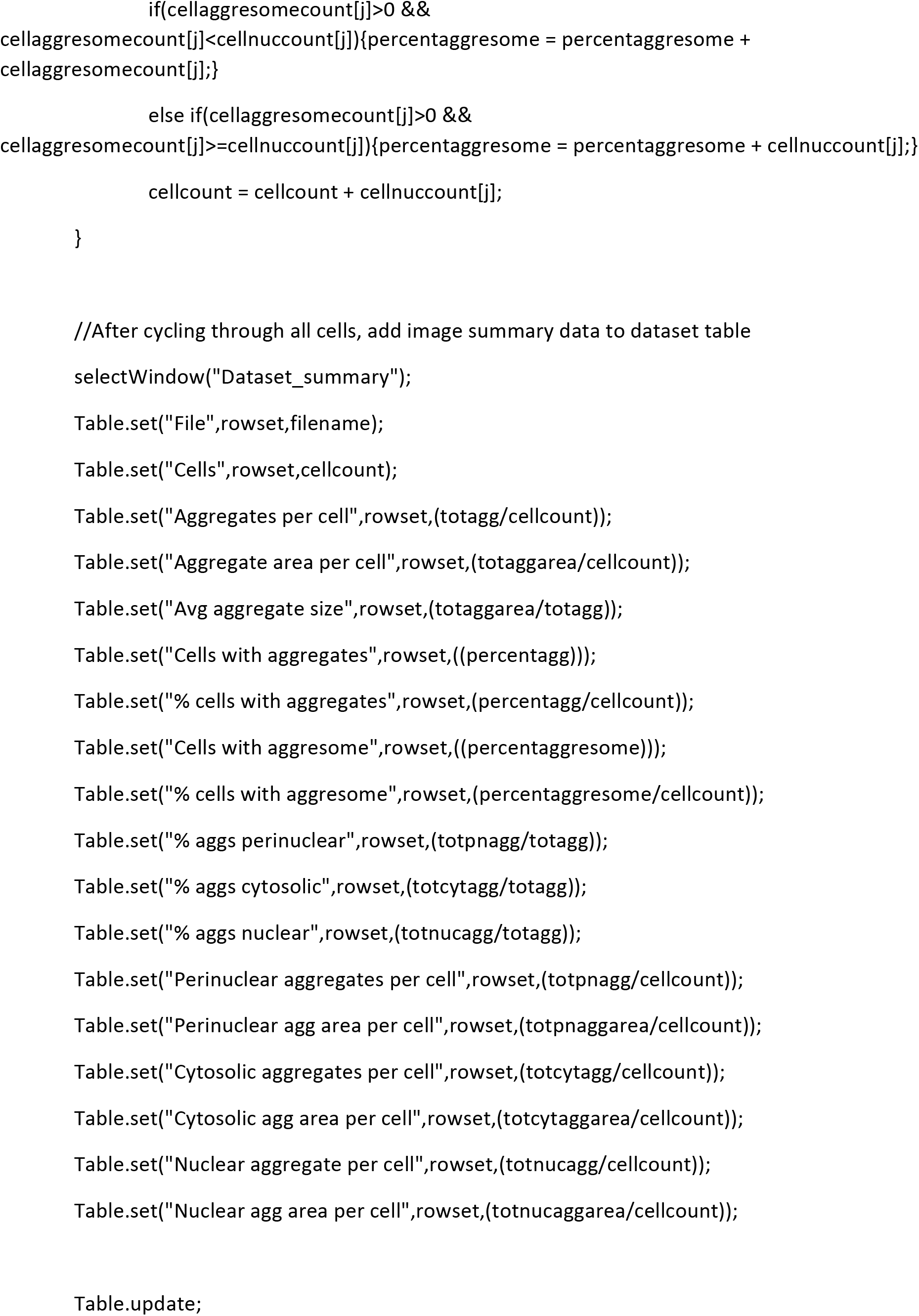

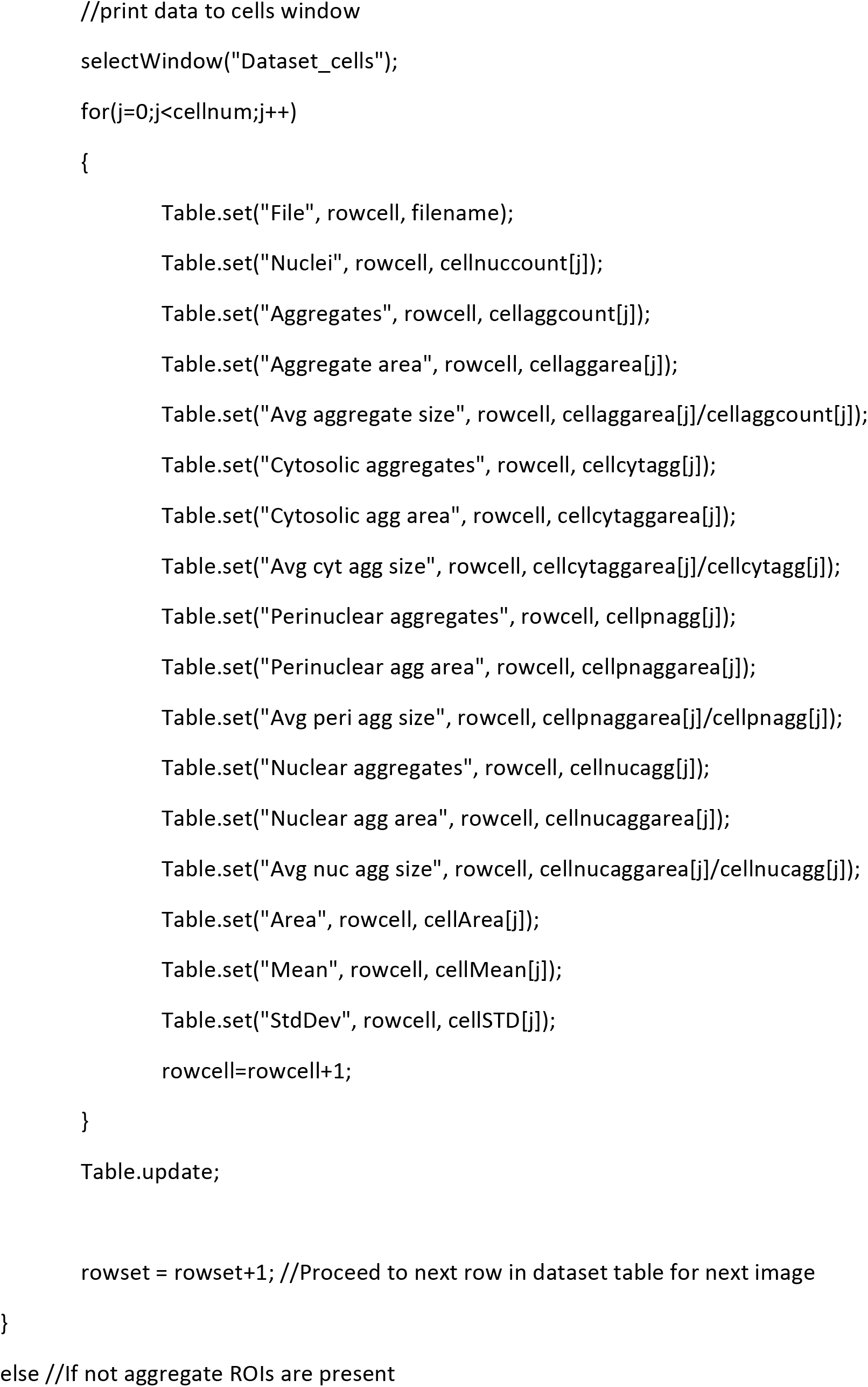

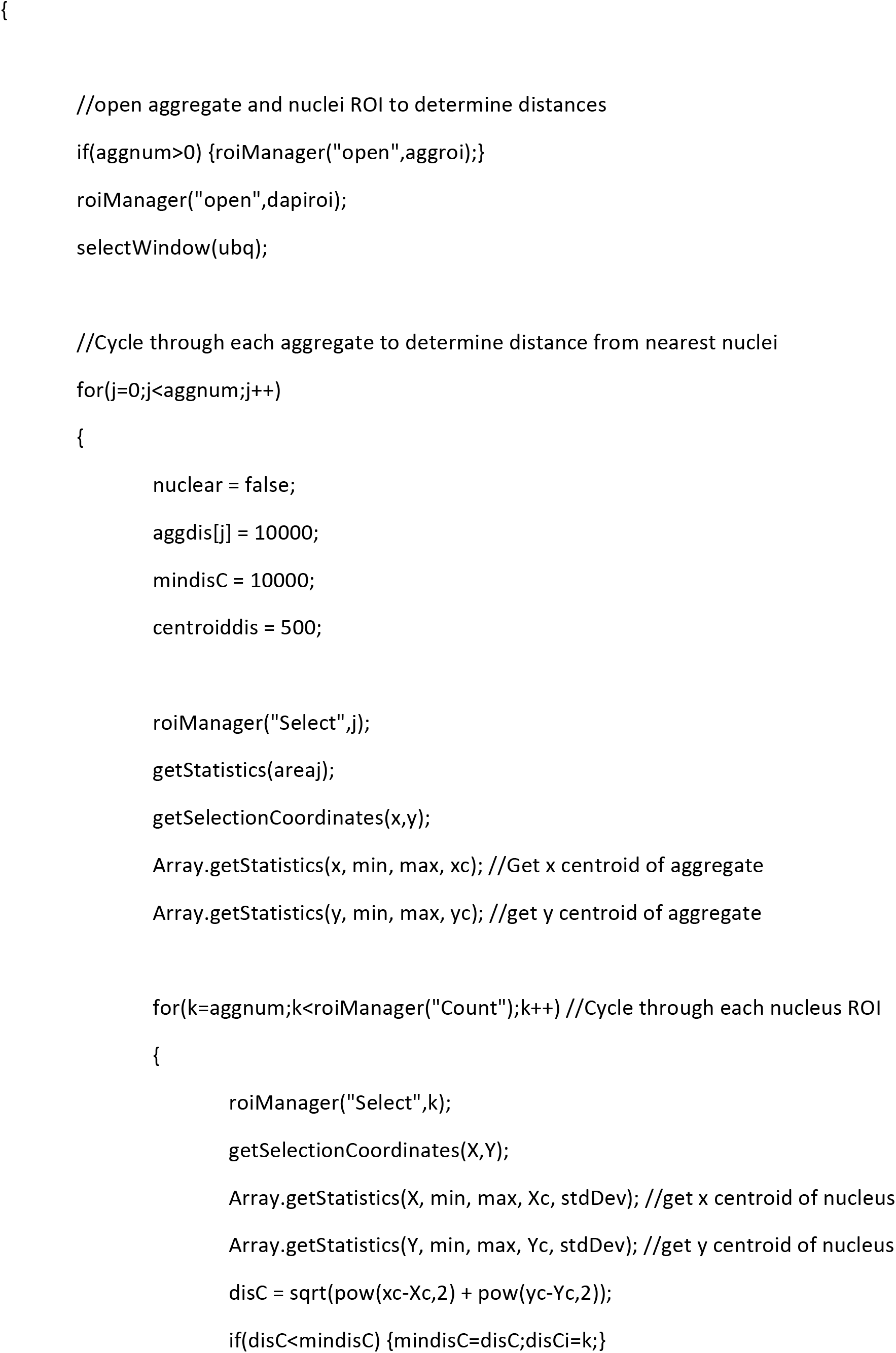

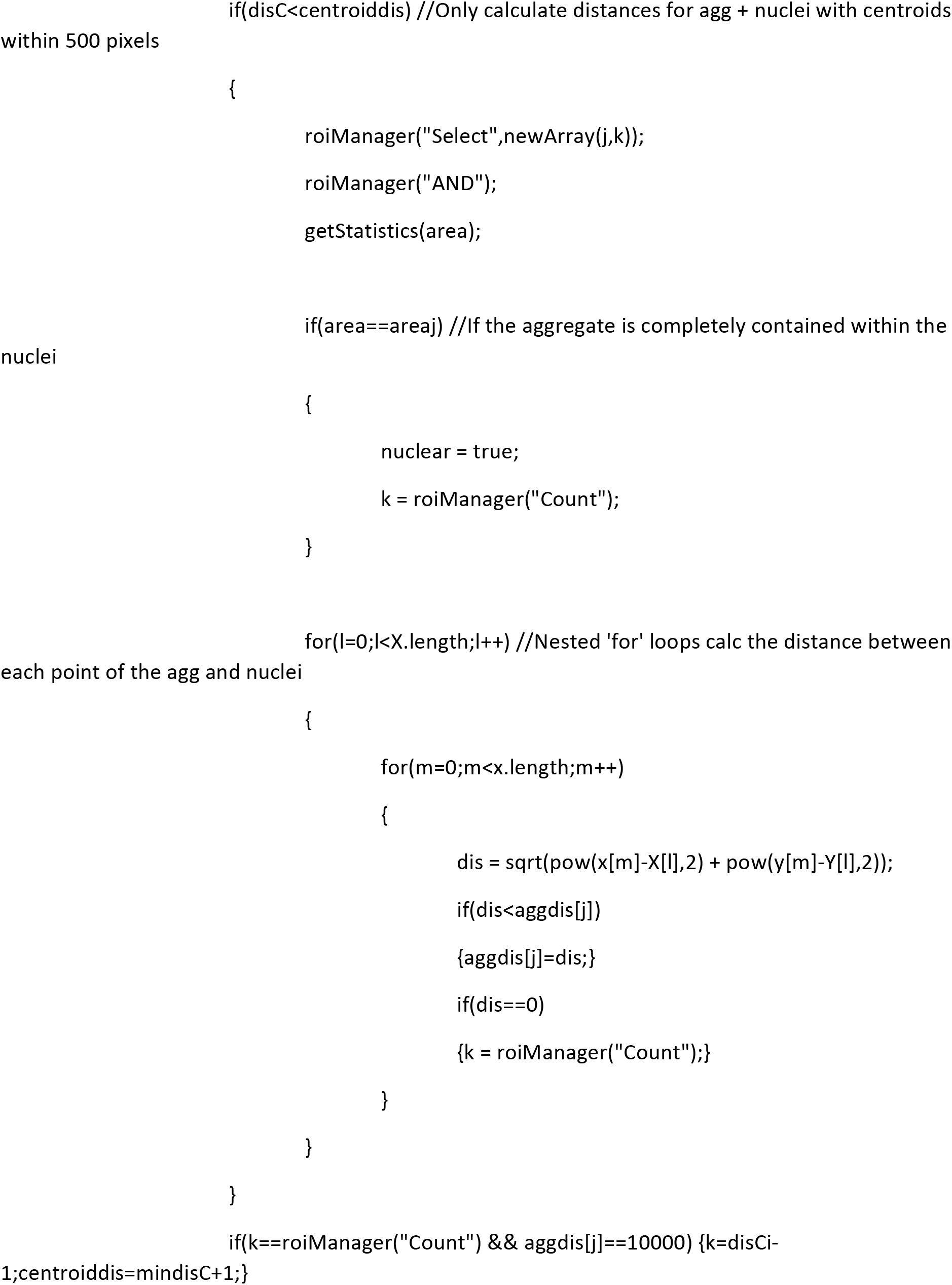

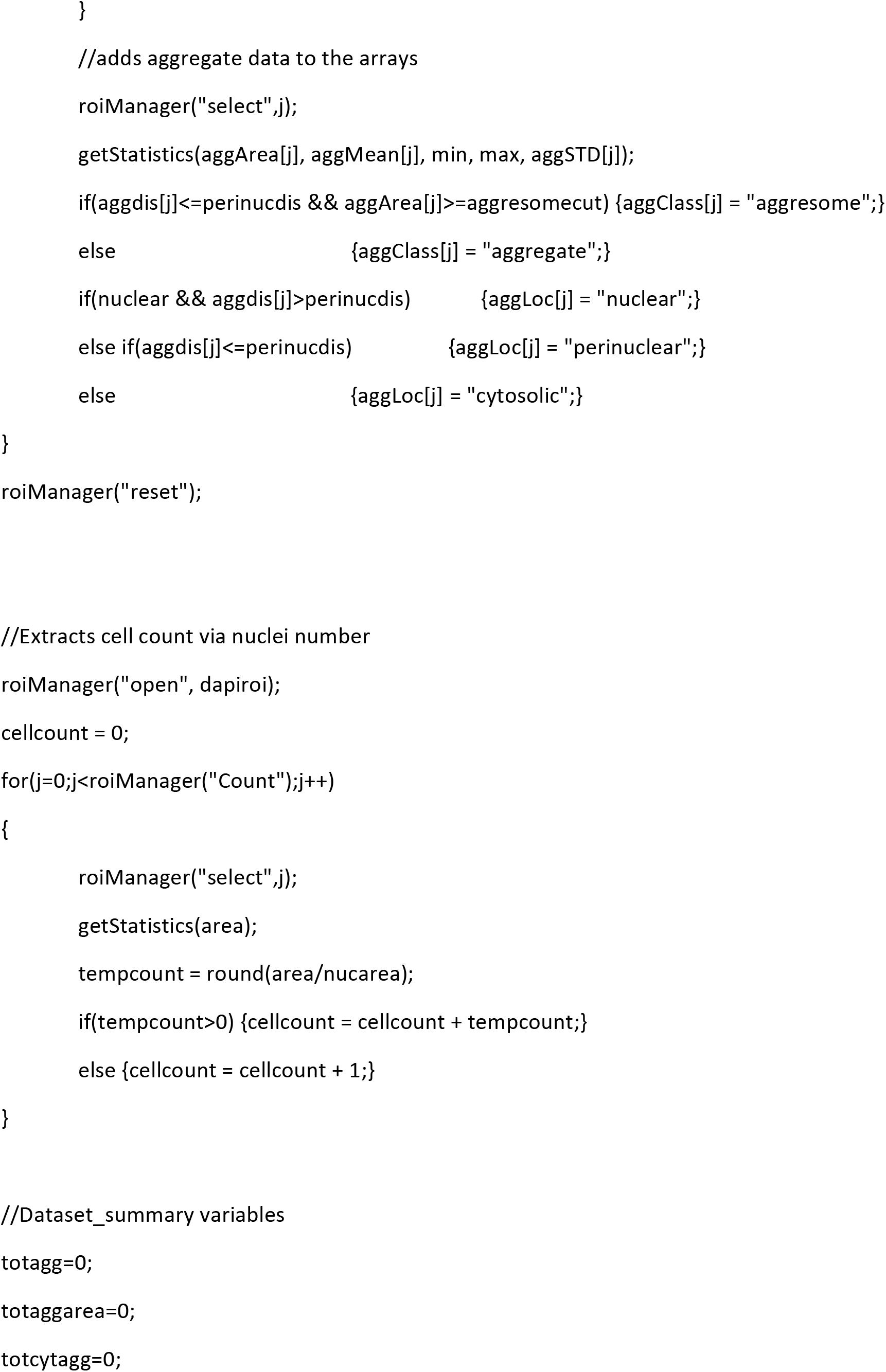

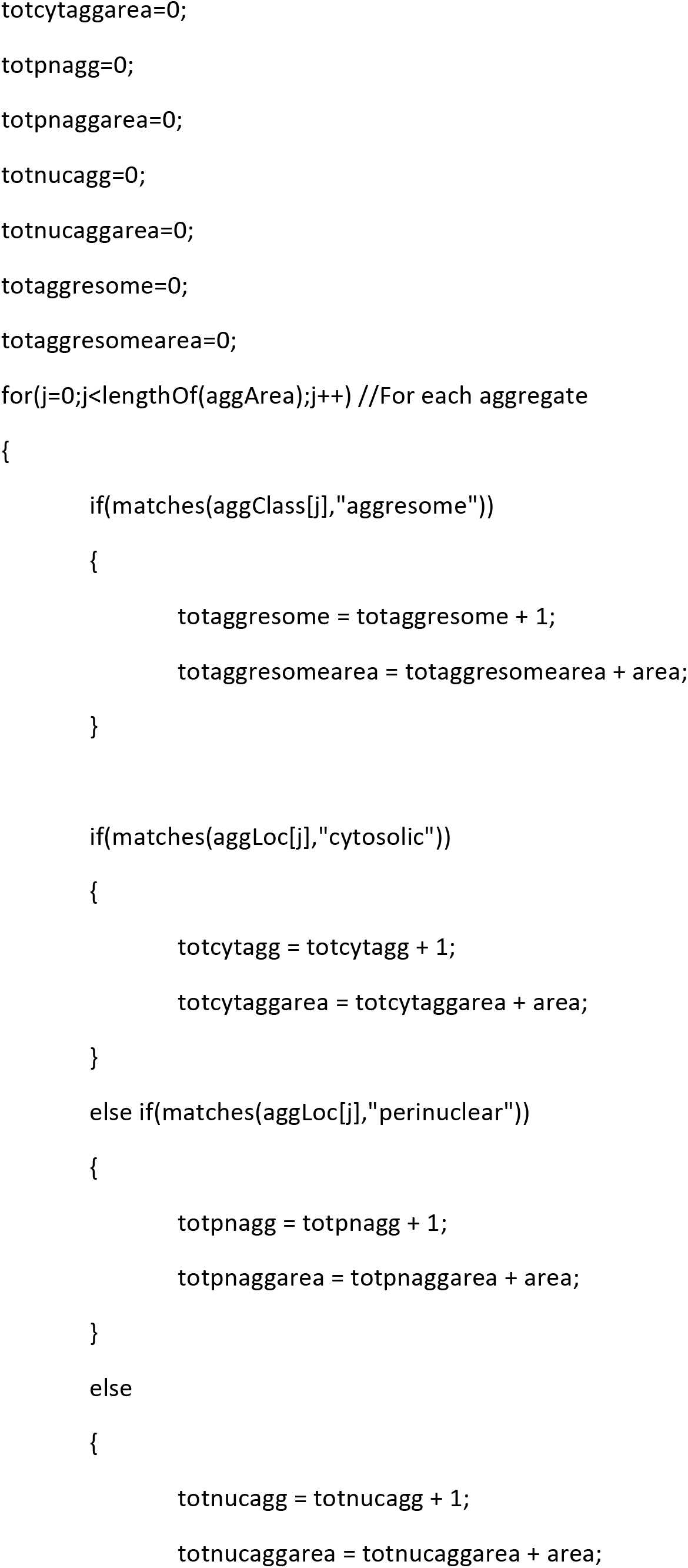

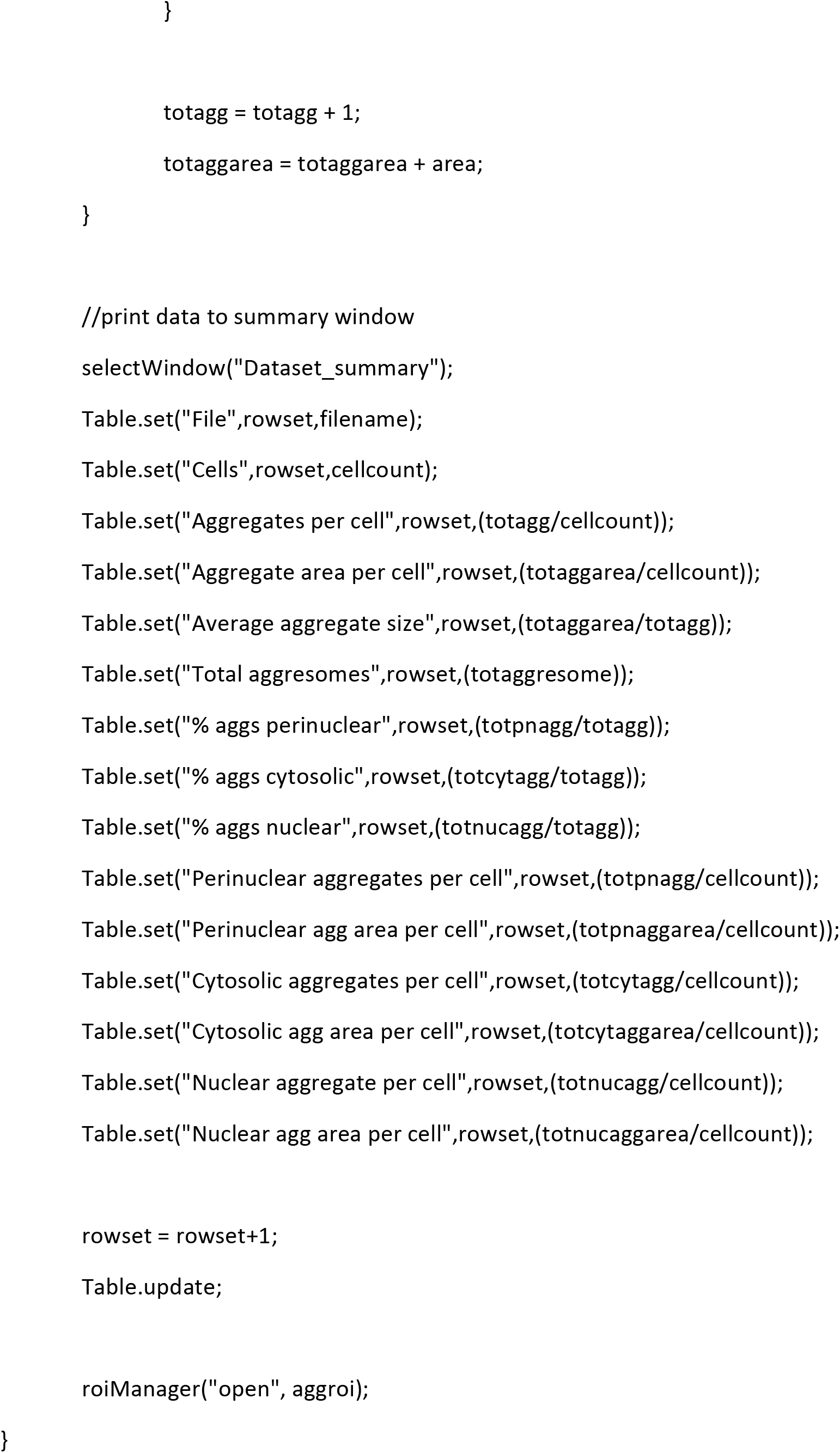

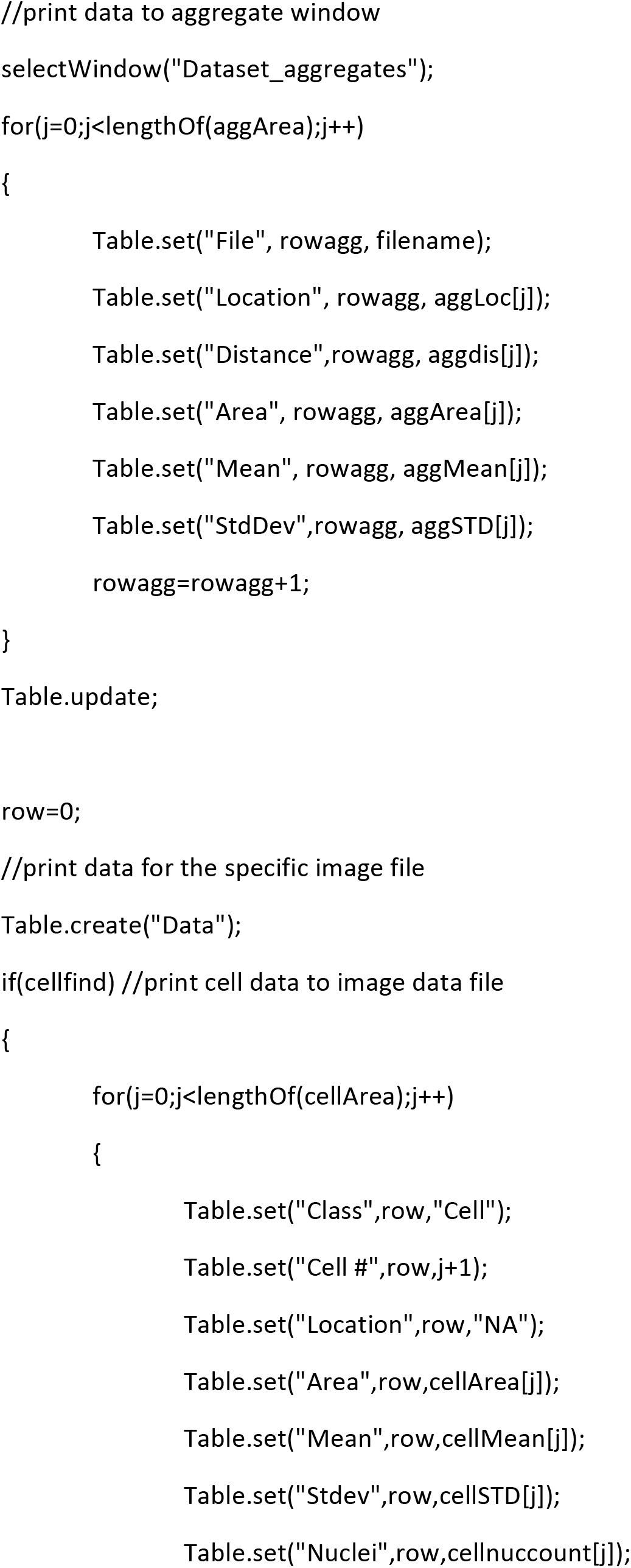

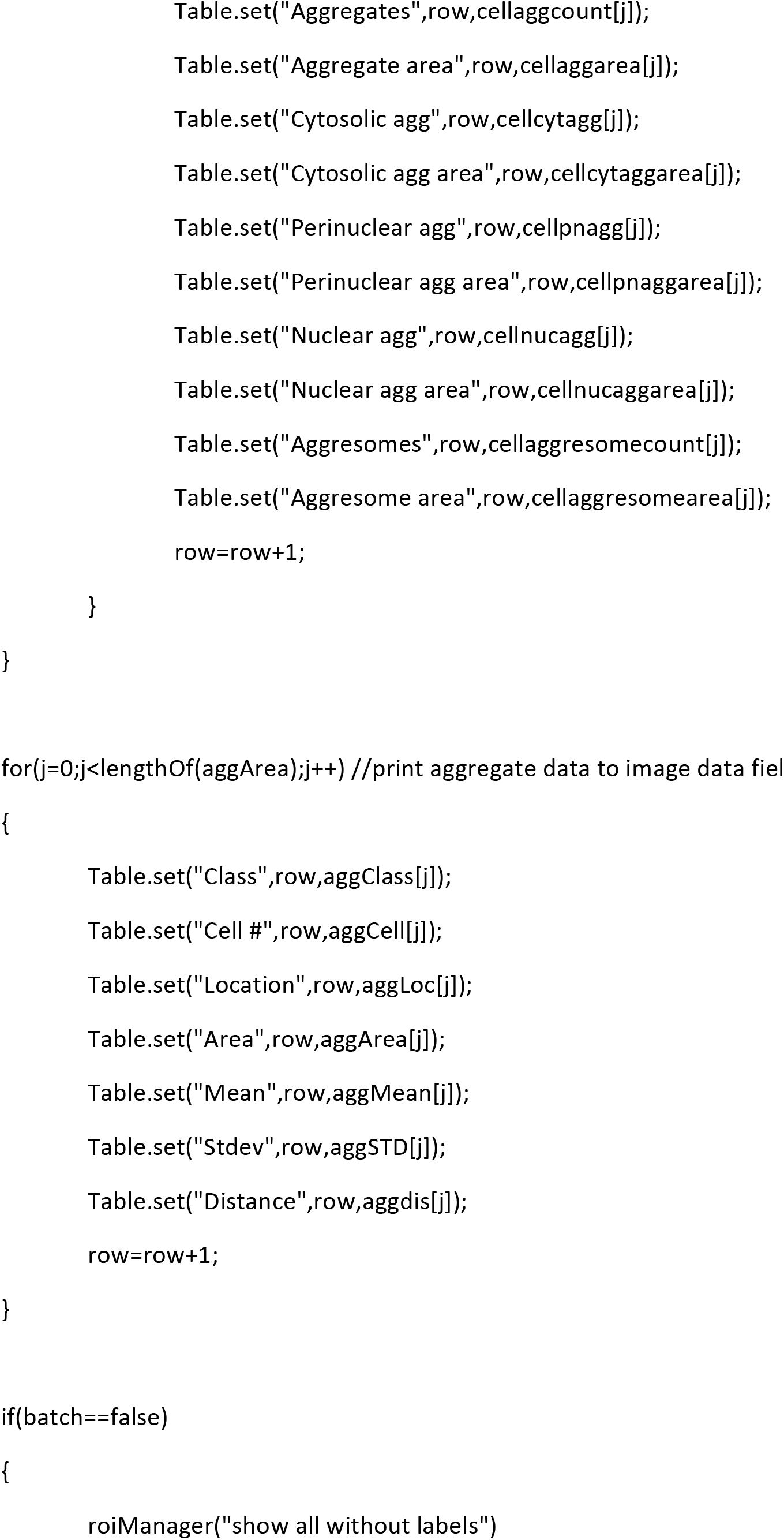

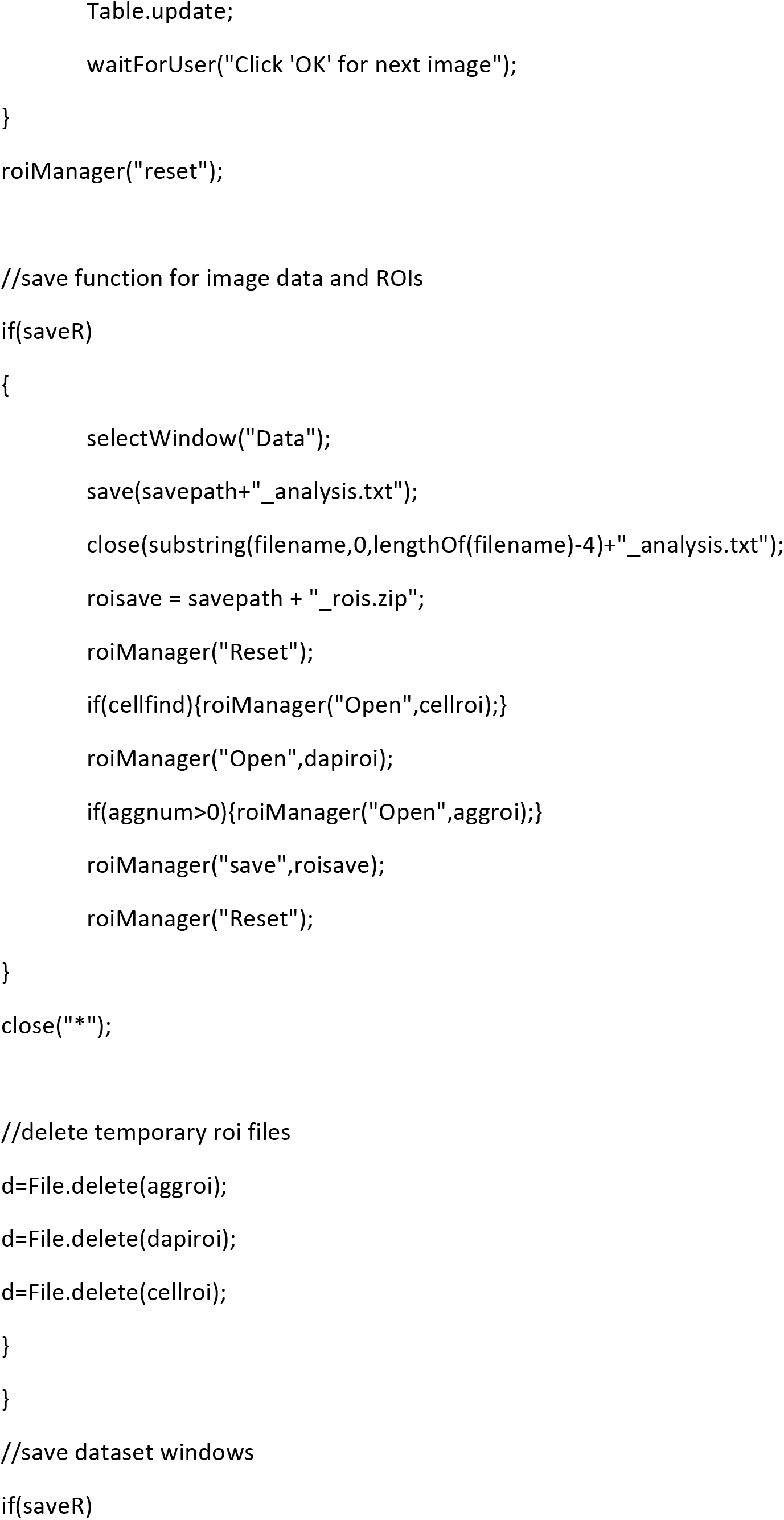

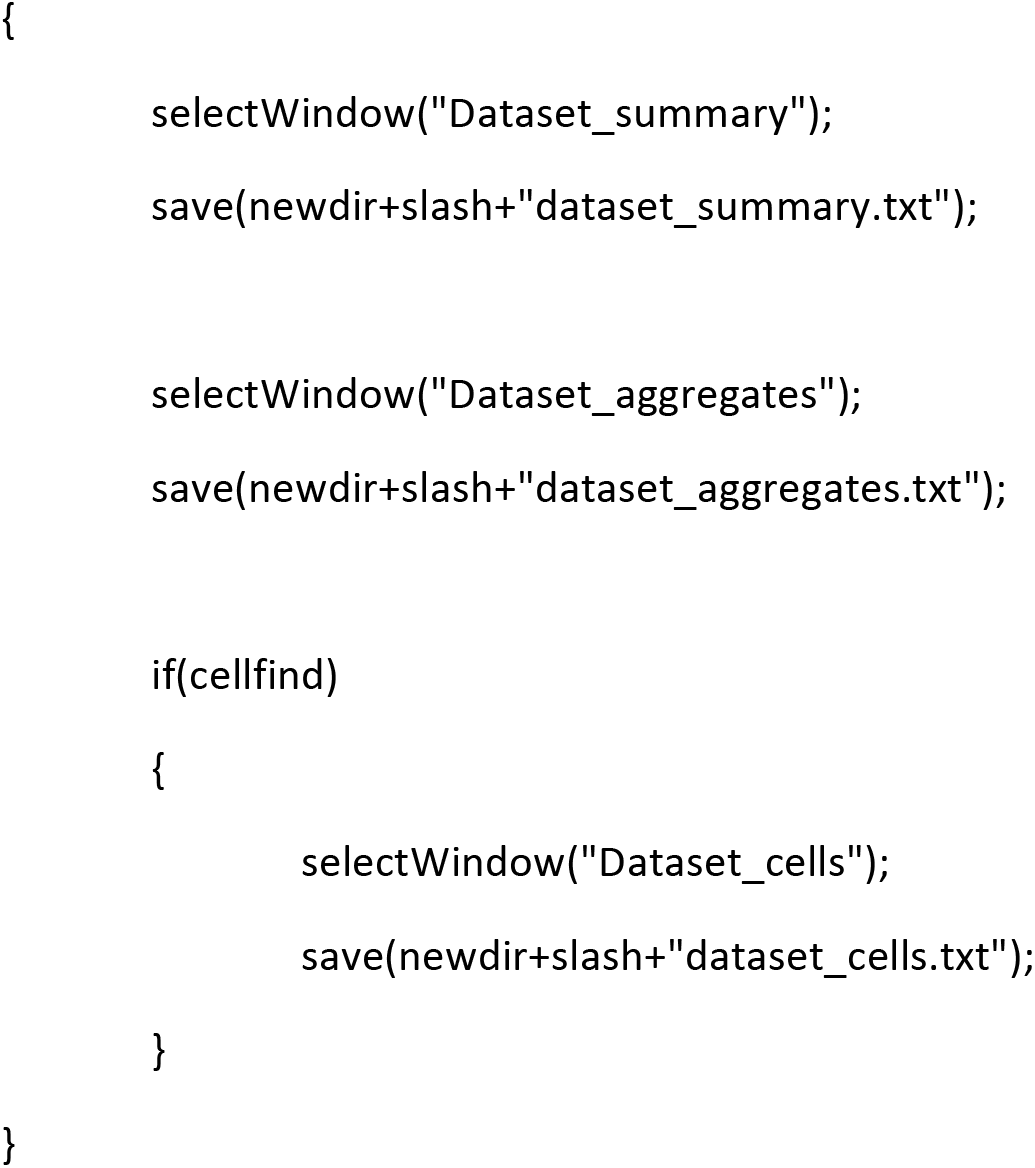

## Supplementary Information 3

### Instructions for AggreCount analysis

Sample images and macro code are available at: https://tufts.box.com/s/ys3kktb5ujdnilyqu8mnup3mrtiej6b6

#### Quick start guide

1. Download the Aggrecount macro

a. Install via the plugin menu “Install…” option
b. Or drag and drop onto the ImageJ toolbar to open in the macro editor
2. Place all images for analysis into one folder

a. All images should have the same channels in the same order
3. Run Aggrecount from either the plugin menu or the “Run” button in the macro editor
4. Select the folder with images for analysis
5. Select “Setup”
6. Follow instructions to designate nuclei, aggregate, and cell body channels

a. If there is no cell body stain, select nuclei channel
7. Select “Get image” to adjust threshold (applied to all images during analysis)

a. Choose an image and adjust threshold
b. Continue until satisfied with threshold

i. Press “Next step”
8. Select “Get image” to view aggregate size and distance from nucleus

a. Note size of noise vs signal for size cut off
b. Note distance from nuclear perimeter

i. This distance designates the perinuclear zone
c. Press “Next step”
9. Select “Get image” to view different cell body processing methods and strictness

a. Note method and strictness from image titles
b. Press “Next step (settings)”
10. Main settings window

a. Threshold values are auto-populated from setup
b. Adjust perinuclear distance, aggregate min/max levels as previously noted
c. Adjust strictness of cell body processing and method as previously noted
d. Adjust other settings as desired (see below for explanations)
e. Batch mode

i. Unchecked

1. Macro will pause after acquiring ROIs and show user
2. Useful to ensure fidelity of processing and ROI capture before running a batch mode analysis
ii. Checked

1. Macro will proceed to analyze all images in folder in the background
2. Useful when processing an entire experiment
11. Data files will be saved after all images have been analyzed in the folder selected

a. Look for “AC_analysis#” folder

#### AggreCount analysis setup

Users may decide to follow the setup process outlined below or manually input settings by selecting “Setup” or “Manual input” after selecting image analysis folder. It is recommended that users utilize the set-up process for first time analyses of an experiment to help select proper settings.

1. Open FIJI (v1.52p or later)
2. Download the AggreCount macro

a. Install using the “Install…” option under the “Plugin” menu in the ImageJ toolbar
b. Alternatively, drag and drop the Aggrecount macro icon onto the ImageJ toolbar to open Aggrecount in the macro editor
3. Place all images for analysis into one folder

a. All images supported by the Bio-Formats importer may be used, however, images must contain multiple channels
4. Select the Aggrecount macro from the “Macros” menu under the “Plugins” menu in the ImageJ toolbar

a. Alternatively, select “Run” from the bottom of the macro editor window
5. Select the folder with images for analysis
6. The macro will automatically open the first image and ask the user to select the channels for nuclei, aggregates, and cell body

a. Change the channel to the immunostain specified by the window and select “Okay”
b. If there is no stain for cell bodies, select the nuclei channel
7. The next window will prompt the user to get an image to adjust thresholding

a. Select “Get image” and choose an image with aggregates
b. Adjust the threshold so aggregates are highlighted in red with as little background as possible and click “Okay”

i. Do not close the threshold window
c. Select “Get image” again and choose an image without aggregates
d. The previous threshold will be shown, adjust appropriately and click “Okay”
e. Select “Get image” again and choose an image with aggregates
f. Adjust threshold as previously mentioned and click okay
g. Either continue cycling through images to adjust threshold or, when satisfied, select “Next step”

i. This threshold will be applied to all images in the analysis
8. The next window will prompt the user to get an image for determining aggregate size cutoff and perinuclear distance cutoff

a. Select “Get image” and choose an image with aggregates
b. The macro will identify aggregates using the previously set threshold and nuclei using an automated thresholding system
c. The image will be displayed with the size of each aggregate in square microns as well as the distance from the nucleus perimeter in pixels
d. Use the “+” and “−” buttons on the keyboard to zoom in and out of the image
e. Note the smallest and largest size of ‘true’ aggregates vs noise as well as the distance from the nucleus

i. The default setting for the perinuclear zone is 10 pixels for 60x magnification images, this may need to be adjusted for images of different magnification
ii. The default setting for aggregate size is 0.1 – 25mm^2^ which is applicable to most aggregate images
f. Select “Okay” when finished noting sizes and distances
g. Either continue cycling through images to note sizes and distances or select “Next step” to proceed
9. The next window will prompt the user to get an image to determine which cell body processing method and strictness will work best

a. The default setting of “Segmentation” with strictness of 5 is optimized to work with most images

i. Dense cells require *lower* strictness for segmentation and *higher* strictness for thresholding
ii. Strictness values are optimized to fall between 1 and 10, however, any non-zero value will be accepted but may result in reduced processing quality
b. Select “Get image” and choose one of the sample images
c. The macro will automatically process this image in 3 different strictness for both thresholding and segmentation types of cell body processing
d. This may take a minute
e. The macro will display the results of each processing method and strictness

i. The best one will be the image that has each cell in its own ROI
ii. ROIs without a nucleus will not be included in the analysis, however, excessive segmentation may slow processing speed
iii. Note the best method and strictness from the image titles
f. Either continue cycling through images to adjust threshold or, when satisfied, select “Next step (settings)” to continue to the main settings window

#### AggreCount main settings window

The main settings window will allow users to adjust the parameters described below. Some of these parameters may be populated from performing the setup. Settings are reset to default settings each time the macro is run. Settings from previous runs are saved in the “aggresettings.txt” file within the AC_analysis folder if “Save results” is checked.

1. Aggregate thresholding lower/upper

a. The threshold values will be auto-populated from the user-selected threshold during setup
b. This threshold values may be adjusted in this window
2. Perinuclear distance cutoff

a. The distance from the nuclear perimeter that defines the perinuclear zone measured in pixels
3. Aggresome min size

a. The size cutoff for designating a perinuclear aggregate an aggresome
4. Aggregate min/max size

a. Size cutoff for aggregate ROI acquisition
b. Aggregate ROIs smaller or larger than these values will not be captured
5. Nuclei size/strictness

a. Minimum size of nuclei (helps exclude noise)
b. Strictness defines the amount of background subtraction during nuclei processing

i. Lower values *decrease* background subtraction, higher values *increase* it
6. Cell size/strictness

a. Minimum size of cell ROIs (helps to exclude noise)
b. Strictness alters cell body processing depending on the method

i. Thresholding strictness increases or decreases background subtraction in the same manner as nuclei strictness
ii. Segmentation strictness defines the “prominence” value in the “Find maxima…” function

1. Smaller strictness values will increase total number of segments while larger values to decrease it
2. Cell segments are checked for nuclei and segments without nuclei are deleted
7. Nuclei/aggregates/Cell bodies channel/slice/frame

a. These fields are auto-populated from earlier in the setup process
b. If the images do not have slices (z-stack) or frames (movie), these values should be 1
c. If the images do not have a cell bodies channel, do not change these values and uncheck “Find cell bodies”
d. NOTE – all images to be analyzed must have all channels in the same order
8. Find cell bodies checkbox

a. If checked, the macro will find cell bodies using the processing method and strictness designated, aggregate distance will be calculated from the nucleus within the same cell and will output cell-by-cell data in the save file dataset_cells
b. If unchecked, the macro will not find cell bodies, aggregate distance will be calculated from the closest nuclei, and will output image summary data and aggregate data but not single cell data
9. Cell processing method

a. Segmentation uses the “Find maxima.” function in ImageJ to locate cell centers and then uses a watershed algorithm to create a Voronoi diagram. This is used to find cell ROIs which are checked for nuclei
b. Thresholding uses the “Enhance contrast.” function to increase fluorescent signal, then adds in nuclei area before applying an automated threshold
10. Save results

a. If checked, the macro will create a new folder in the image analysis folder previously selected named “AC_analysis#” where individual data files and ROI files as well as dataset files will be saved
b. If unchecked, the macro will create tables in ImageJ with all data captured but will not save it

i. The user may manually save this data if wanted
11. Batch mode

a. If checked, the macro will proceed to analyze all images in the folder selected in the background without user input, greatly increasing processing speed
b. If unchecked, the macro will prompt user before each image as well as display ROIs for nuclei, aggregates, and cells after capture
c. It is suggested that users uncheck batch mode before running a batch analysis to ensure fidelity of ROI capture for a subset of images

#### AggreCount analysis (non-batch mode)

1. After proceeding through AC setup or manually entering AC settings, leave “Batch mode?” unchecked and press “OK”
2. For each file that is in the folder, the user will be prompted to analyze it (“Continue”), skip it (“Skip”), or quit the macro (“Cancel”)

a. Non-image files or incompatible image files may appear but will be skipped even if the user selects “Continue”
3. After pressing “Continue”, nuclei will be processed, and an auto-threshold applied. The nuclei ROIs will be displayed for the user to view

a. If nuclei ROIs are too large or there is excess noise, consider increasing nuclei processing strictness in the main menu
b. If nuclei ROIs are too small or nuclei are being excluded, consider decrease nuclei processing strictness in the main menu

i. To change nuclei processing strictness, quit out of the macro, run it again, and select “Manual input” to input desired settings
c. Select “OK” to continue
4. Aggregates will be processed, and a threshold applied (as previously set by the user). The aggregate ROIs will be displayed for the user to view

a. If aggregate thresholding requires further refinement, consider quitting the macro and restarting the setup process
b. Select “OK” to continue
5. Cells will be processed as previously determined by the user (method and strictness). The cell ROIs will be displayed for the user to view

a. Thresholding

i. If ROIs are too large or multiple cells are captured in one ROI, consider increasing cell processing strictness
ii. If ROIs are too small or cells are being excluded, consider decreasing cell processing strictness
b. Segmentation

i. If ROIs are too large and contain multiple cells, consider *decreasing* cell processing strictness
ii. If ROIs are too small and are splitting cells, consider *increasing* cell processing strictness
c. Select “OK” to continue
6. Aggregates and nuclei will be assigned to cells and aggregate localization is determined

a. The dataset tables and the image data table will be updated with the values from the current image
7. The macro will pause after the entire analysis has been finished and will display the image along with all ROIs captured (nuclei, cell bodies and aggregates)

a. Additionally, data will be presented in the “Data” table with cells and aggregates for the user to view
b. Select “OK” to continue
8. The macro will again present files in the folder for the user to analyze or skip

a. The user may proceed through all the images in this manner
b. NOTE – the dataset tables will NOT save unless all files have been analyzed or skipped. To save these tables after canceling the macro run, select “File” -> “Save” on the table window.

#### AggreCount analysis (batch mode)

1. After proceeding through AC setup or manually entering AC settings, check “Batch mode?” and press “OK”
2. All images will be analyzed in the folder selected
3. Four tables will be visible to the user

a. Dataset_summary
b. Dataset_aggregates
c. Dataset_cells (if “Find cells” is checked)
d. Data
4. Each will be updated after each image is processed
5. Each line in Dataset_summary represents an image file so this table is ideal for tracking analysis process
6. WARNING – do not close these tables or select specific lines as it may interfere with the analysis

a. These tables may be minimized
7. If the user wishes to cancel the batch mode analysis, close the Dataset tables
8. After the analysis is finished, the tables are saved to the “AC_analysis#” folder in the folder containing images

#### Analysis output

All data files are saved as tab-delimited .txt files. These may be easily imported into excel or another data processing software. In addition to the data tables described below, ROIs for each image are saved as .zip files within the AC_analysis folder. Each ROI zip file contains ROIs for cells, nuclei, and aggregates in that order.

Dataset_summary

Summary data by image

– File – Filename
– Cells – Number of cells imaged (uses # of nuclei)
– Aggregates per cell – Total aggregates in image divided by total number of cells (nuclei)
– Aggregate area per cell – Total aggregate area in image divided by total number of cells
– Avg aggregate size – Total aggregate area divided by total aggregates
– Cells with aggregates* – Total number of cells with any type of aggregate
– % cells with aggregates* – Percentage of all cells with any type of aggregate
– Cells with aggresome* – Total number of cells with an aggregate above the user defined size cutoff within the perinuclear zone (aggresome)
– % cells with aggresome* – Percentage of all cells with an aggregate above the user defined size cutoff within the perinuclear zone (aggresome)
– Total aggresomes** – Total number of aggresomes in the image
– % aggs perinuclear – Perinuclear aggregates divided by total aggregates
– % aggs cytosolic – Cytosolic aggregates divided by total aggregates
– % aggs nuclear – Nuclear aggregates divided by total aggregates
– Perinuclear aggregates per cell – Total perinuclear aggregates divided by total cells (nuclei)
– Perinuclear aggregate area per cell – Total perinuclear aggregate area divided by total cells (nuclei)
– Cytosolic aggregates per cell – Total cytosolic aggregates divided by total cells (nuclei)
– Cytosolic aggregate area per cell – Total cytosolic aggregate area divided by total cells (nuclei)
– Nuclear aggregates per cell – Total nuclear aggregates divided by total cells (nuclei)
– Nuclear aggregate area per cell – Total nuclear aggregate area divided by total cells (nuclei)

Dataset_cells*

Summary data by cell

– File – Filename of the image that contains the cell
– Nuclei – Number of nuclei contained within a cell ROI
– Aggregates – Total number of aggregates within a cell
– Aggregate area – Total aggregate area within a cell (um^2)
– Avg aggregate size – Total aggregates within the cell divided by total aggregate area within the cell
– Cytosolic aggregates – Total number of cytosolic aggregates within a cell
– Cytosolic agg area – Total cytosolic aggregate area within a cell
– Perinuclear aggregates – Total number of perinuclear aggregates within a cell
– Perinuclear agg area – Total perinuclear aggregate area within a cell
– Nuclear aggregates – Total number of nuclear aggregates within a cell
– Nuclear agg area – Total nuclear aggregate area within a cell
– Area – Total area of a cell ROI
– Mean – Mean fluorescent intensity from a cell ROI
– StDev – Standard deviation of fluorescent intensity from a cell ROI

Dataset_aggregates

Summary data by aggregate

– File – Filename of image that contains the aggregate
– Location – Cellular compartment of an aggregate (nuclear, perinuclear, cytosolic)
– Distance – Distance in pixels between the nearest edge of an aggregate to a nucleus ROI
– Area – Area of the aggregate
– Mean – Mean fluorescent intensity from aggregate ROI
– StDev – Standard deviation of the fluorescent intensity from aggregate ROI

‘Image name’_analysis

Summary data for a specific image. Contains data for both cell and aggregate ROIs

– Class – Classification of ROI: cell, aggregate, aggresome
– Cell # – Cell ID that links aggregates to cells
– Location – Subcellular location of aggregates (cytosolic, perinuclear, nuclear). Cell ROIs will be labeled “NA”
– Area – Area of a ROI
– Mean – Mean fluorescence intensity of a ROI
– StDev – Standard deviation of fluorescence intensity of a ROI
– Nuclei* – Number of nuclei contained within a cell ROI

- Aggregate ROIs will have “0” in this column
– Aggregates* – Total number of aggregates in a cell

- Aggregate ROIs will have “0” in this column
– Aggregate area* – Total sum of aggregate area in a cell

- Aggregate ROIs will have “0” in this column
– Cytosolic agg* – Total number of cytosolic aggregates in a cell

- Aggregate ROIs will have “0” in this column
– Cytosolic agg area* – Total sum of cytosolic aggregate area in a cell

- Aggregate ROIs will have “0” in this column
– Perinuclear agg* – Total number of perinuclear aggregates in cell

- Aggregate ROIs will have “0” in this column
– Perinuclear agg area* – Total sum of perinuclear aggregate area in a cell

- Aggregate ROIs will have “0” in this column
– Nuclear agg* – Total number of nuclear aggregates in a cell

- Aggregate ROIs will have “0” in this column
– Nuclear agg area* – Total sum of nuclear aggregate area in a cell

- Aggregate ROIs will have “0” in this column
– Aggresomes* – Total number of aggresomes in a cell

- Aggregate ROIs will have “0” in this column
– Aggresome area* – Total aggresome area in a cell

- Aggregate ROIs will have “0” in this column
– Distance – Aggregate distance in pixels from a nucleus ROI perimeter

- Cell ROIs will have “0” in this column

*designates fields or data tables that are only present in analyses that use cell ROIs

** designates fields that are only present in analyses that do not use cell ROIs

